# More than a stick in the mud: Eelgrass leaf and root bacterial communities are distinct from those on physical mimics

**DOI:** 10.1101/2021.05.31.446483

**Authors:** Melissa R. Kardish, John. J. Stachowicz

## Abstract

We examine the role of physical structure vs. biotic interactions in structuring host-associated microbial communities on a marine angiosperm, *Zostera marina,* eelgrass. Across several months and sites, we compared microbiomes on physical mimics of eelgrass roots and leaves to those on intact plants. We find large, consistent differences in the microbiome of mimics and plants, especially on roots, but also on leaves. Key taxa that are more abundant on leaves have been associated with microalgal and macroalgal disease and merit further investigation to determine their role in mediating plant-microalgal-pathogen interactions. Root associated taxa were associated with sulfur and nitrogen cycling, potentially ameliorating environmental stresses for the plant. Our work identifies targets for future work on the functional role of the seagrass microbiome in promoting the success of these angiosperms in the sea.

**Originality-Significance Statement:** We show that eelgrass establishes a distinct microbial community from a physical mimic on both its leaves and roots. This is, to our knowledge, the first comparison of seagrass to a mimicked physical environment. Insights from our study establish bacterial targets for future functional studies of seagrass-microbiome interactions.

## Introduction

The role of microbiomes in host ecology is increasingly recognized as a potentially important force driving the ecology and dynamics of a broad range of ecosystems. Host-microbe interactions range in strength and direction (McFall-Ngai *et al*., 2013; Hammer *et al*., 2019) with microbes providing net benefits to their hosts in some cases, parasitizing hosts or other members of the microbial community in others, and participating in many symbiotic relationships in between these extremes (Trivedi *et al*., 2020). Yet hundreds or thousands of microbial taxa associate with any given host, and we generally know little about the extent to which most associations rely on specific host traits such as morphology, physiology or metabolites. In most cases, our knowledge is limited to comparing host-associated microbes with a larger environmental pool, such as soils, and identifying taxa over-represented on hosts compared to the environment (Knights *et al*., 2011; Fahimipour *et al*., 2017; Xiong *et al*., 2021). These approaches often identify hundreds of taxa positively associated with hosts, still leaving a major challenge for developing an understanding of the extent to which particular microbes interact closely with, and impact, hosts. However, this approach does not distinguish the role of the provision of physical structure vs. host-specific biology; incorporating this level of distinction would identify taxa that require not just the structure but the presence of a living host and therefore a greater potential for reciprocal interactions with the host. One way to distinguish the relative importance of physical structure from living organisms is to use physical mimics to assess how microbial communities develop differently in the absence of biotic interactions with the host.

Mimicking environments to learn more about host-microbe interactions -- whether through simple physical models, reconstituting biochemical environments, or even using germ-free organisms -- has been used across a diversity of taxa to assess critical members microbial communities as well as to assess how the overall structure of their communities vary under stress. For example, in terrestrial plants, finely mimicked leaf surfaces have led to insights on where *E. coli* resides on spinach leaves based on water retention on structural mimics (Zhang *et al*., 2014) and an artificial human gut has been used to demonstrate interactions between antiinflammatory bacteria and epithelial cells (Zhang *et al*., 2021). While often used to understand how an organism interacts with the microbes, these mimics can also be used to describe community shifts that would occur without the host, identify key partners, and consider differences in assembly (e.g., Lee *et al*., 2019).

Host-associated microbial communities have been identified across organisms to assemble in distinct non-random ways though the degree of specificity varies (Taylor *et al*., 2004; Ambika Manirajan *et al*., 2016). Microbial associates can be highly specific (e.g., the bobtail squid and *Vibrio fisherii, McFall-Ngai and Ruby, 1991)* or might be transitory/happenstance associations (e.g., high numbers of soil bacteria in Lycaenid butterfly gut microbiomes; (Whitaker *et al*., 2016). In addition a host can interact with a microbiome with different levels of restrictiveness: a host might harbor a highly restricted environment potentially heavily modified by a host (e.g., a gut microbiome; Garland *et al*., 1982; Rinninella *et al*., 2019) or a less restrictive environment where even with environmental modification from the host many microbes could enter the community (e.g., skin microbiome; Byrd *et al*., 2018). Distinguishing among these types of microbial communities may offer insight into the intensity of interactions with a host. For instance, if we distinguish that a less restrictive surface had a microbiome unassociated specifically with a host, we might infer limited direct unique interactions with that host.

We investigate the role of physical structure vs. biotic interactions in structuring the surface microbiomes of seagrass, specifically, the eelgrass, *Zostera marina.* Eelgrass is a marine flowering plant; its roots and rhizomes grow in highly sulfidic sediment and its leaves, while primarily exposed to seawater, can be periodically exposed to air at low tides (Jørgensen, 1982). Seagrass leaves are distinct from terrestrial angiosperms in several ways, including absence of stomata as well as primary exposure to seawater rather than air (Olsen *et al*., 2016). Our previous work showed broad overlap in the composition of leaf and water microbiomes (Fahimipour *et al*., 2017), but did find some microbes preferentially associated with leaves. Comparison of microbiomes among species of seagrass that grow in the same environment show that some harbor distinct microbial communities on their leaves from other species (Garcias-Bonet *et al*., 2020) while some seagrass species have broad overlap in their microbiomes (Ugarelli *et al*., 2017; Kaimenyi *et al*., 2018). At least some taxa are disproportionately found on seagrass compared to water, though it is not clear the extent to which leaf microbiomes differ from those that accumulate on inert surfaces in marine systems, where biofilms develop on surfaces at a fast rate (Fischer *et al*., 2014). Mimicked seagrass leaves have long been used to investigate community structures and show similar macroinvertebrate (Healey and Hovel, 2004), fish (Bell *et al*., 1985) and microalgal communities (Horner, 1987; Pinckney and Micheli, 1998) to natural seagrass and provide an obvious approach for distinguishing substrate generalists from seagrass-specific associates. We adopt this approach to narrow the functionally important microbiome of seagrass leaves from the pool of over a hundred of taxa known to be enriched on leaves relative to surrounding seawater.

Similarly, root surfaces have bacterial communities distinct from adjacent sediments (Fahimipour et al 2017), but it is not yet clear again how much physical structure vs host biology influences this. Roots inhabit anoxic and highly sulfidic sediments that without mitigation can lead to sulfide intrusion decreasing plant growth and health (Hasler-Sheetal and Holmer, 2015). Various mechanisms exist to mitigate this environment and reduce sulfide intrusion into the plant including radial oxygen loss (ROL) from growing roots (Pedersen *et al*., 2004), direct partnerships with sulfide-oxidizing bacteria (Smith *et al*., 2004), and three-way symbiosis with lucinid clams hosting sulfide-oxidizing bacteria (van der Heide *et al*., 2012; de Fouw *et al*., 2018). However direct association of seagrass with sulfur oxidizers is known (Fahimipour *et al*., 2017) and given the leak of oxygen and sugars out of the roots (Sogin *et al*., 2021), it seems likely that the plant plays an important role in root microbe assembly. Examining mimicked root environments is less common in seagrass than use of seagrass leaves and has focused on sediment stabilization processes (Temmink *et al*., 2020) rather than influence on biotic community structure.

Here, we explicitly test whether seagrass roots and leaves assemble microbiomes that are distinct from physical mimics at a range of sites and seasons with a harbor. By comparing live plants with biologically inactive surfaces mimicking some physical aspects of their environment, we test explicitly whether seagrass cultivates a unique microbiome on its leaves and/or roots. Differences in the bacterial communities between physically mimicked environments and plants could indicate bacteria that might be either attracted by specific biological aspects of a plant or that might be selected by plants for biologically important roles. Thus, such an approach can identify the role of the plant in microbial assembly and hint at specific key processes, while also potentially identifying microbial partners that may play key functional roles in the seagrass microbiome for future experimentation.

## Results

The bacterial assemblages associated with leaves and roots differed from those on their corresponding physical mimics in alpha diversity and/or community composition. However, the extent of this differentiation of eelgrass microbiomes from passive substrates was stronger in roots than leaves, and there were still many amplicon sequence variants (ASVs) shared between mimics and live plants (Figures 1 and 2). These differences among substrates were highly consistent across four sites and three time points (see results below), despite previously identified seasonal and site-specific microbial components at these sites (Kardish and Stachowicz in prep.), indicating a strong impact of live plants on the microbiome. Thus, we focus our presentation of results on the consistent effects of substrate across sites and time (each of which is controlled for in our statistical models).

**Figure 1:**
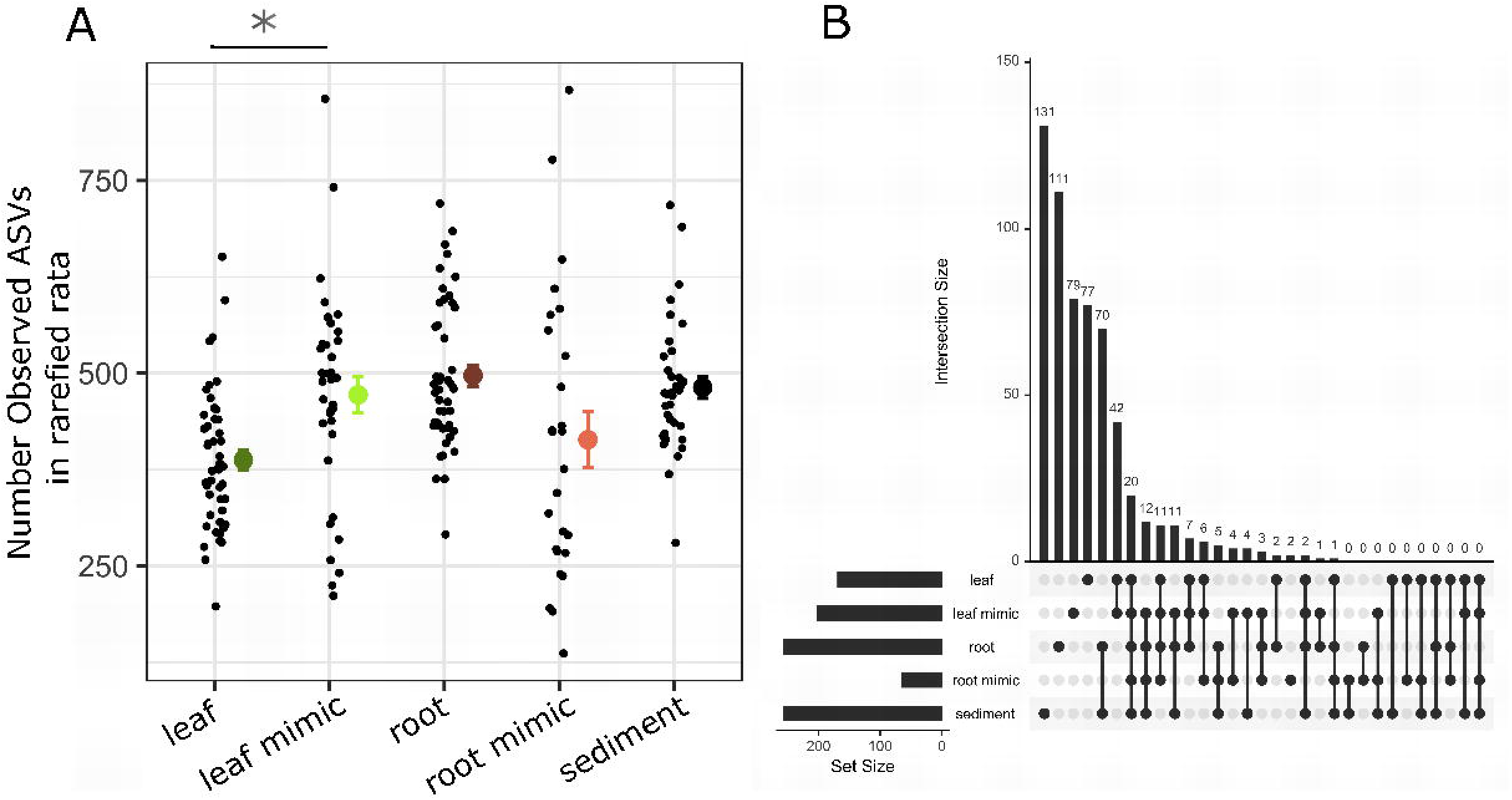
(A) Mean amplicon sequence variant (ASV) richness found in each type of sample we measured. Raw data as well as means and standard errors are presented. Leaf mimic bacterial communities had a higher mean richness than leaf bacterial communities (p < 0.001); root mimic, root, and sediment bacterial communities did not differ in mean community richness. (B) Overlap among core bacterial communities showing shared ASVs present in each sample type in at least 50% of samples at a 1% detection rate. Diagram is a barplot of shared community memberships, equivalent to a Venn diagram.

**Figure 2:**
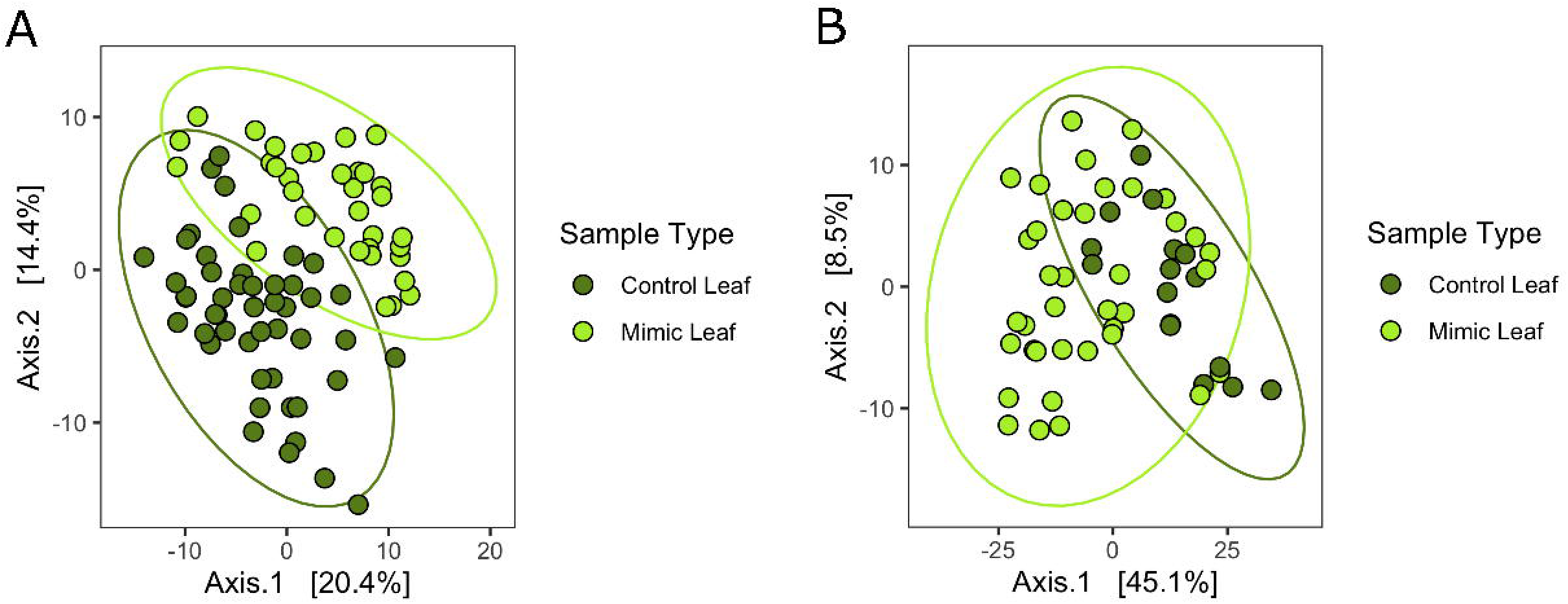
(A) Ordination of bacterial community structure based on principal coordinate analysis of phylogenetic-isometric log-ratio transformed distances. (B) Ordination of predicted Metacyc pathways structure based on principal coordinate analysis of centered log-ratio transformed distances. Bright green points are communities on leaf mimics and dark green points are communities on leaves. Leaf and leaf mimic communities in both analyses are distinct from each other (PERMANOVA p < 0.001).

### Leaves

Leaf mimics had greater ASV richness than leaves (negative binomial glm with crossed random effects for month and site, estimate = 0.19738, standard error = 0.05802, z-value 3.402, p = 0.0007; Figure 1a). The ASV composition of the leaf and mimic communities was different (PERMANOVA, F = 12.09, p = 0.001, r^2^ = 0.13, Figure 2a), though there was no difference in variance among-leaf vs. among-mimic communities (betadisper ANOVA, p = 0.73). When examining core ASVs (present in at least 50% of samples of a type at at least 1% detection rate), we found that roughly half the ASVs found on leaves were not found on any other substrate (77 of 168 ASVs; 46%) while most of the remaining were shared with those on leaf mimics (88 of 168, 52%). (Figure 1b). This degree of overlap in core taxa was the greatest of any pairwise comparison among sample types; The same patterns were present when we examined all ASVs rather than just the core (Supplemental Figure 1). Predicted Metacyc pathways, based on a crossdomain database of metabolic pathways and enzymes (Caspi *et al*., 2014), also differed between leaves and mimics (PERMANOVA, r^2^ = 0.05488, F = 4.7035, p = 0.001), though this effect was weaker than for the sequence based compositional differences (Figure 2b).

Through analysis of specific ASVs that varied between leaves and leaf mimics via DESEQ2, we found 92 ASVs were relatively more abundant on leaves and 49 were relatively more abundant on mimics. Only three families contained more than ten ASVs that varied between mimics and leaves (Table 1): Flavobacteriaceae (eight higher on leaves, eight higher on mimics), Rhodobacteraceae (23 higher on leaves, 13 higher on mimics), and Saprospiraceae (16 higher on leaves, none higher on mimics). Within these families several genera were represented by multiple ASVs. These included *Kordia* (three ASVs higher on leaves, none on mimics), *Ulvibacter* (two higher on leaves, two higher on mimics), *Octadecabacter* (one higher on leaves, one higher on mimics), *Sedimentitalea* (one higher on leaves, one on mimics), *Tateyamaria* (one higher on leaves, one on mimics), *Yoonia-Loktanella* (two higher on leaves), *Lewinella* (three higher on leaves) and *Rubidimonas* (two higher on leaves). Many other families contained fewer than 3 ASVs that varied between leaves and mimics (Table 1, Supplementary Table 1), and within these nine genera contained multiple ASVs that varied between leaves and mimics (Supplementary Table 2). All taxa that varied can be found in Supplemental Table 3.

**Table 1:** For both leaves and root communities, the five families that had the most ASVs vary between mimics and seagrass substrate. See supplemental Tables 1 and 6 for complete lists for leaves and roots respectively.

| Leaves |  |  |
| --- | --- | --- |
| Family | Higher on leaves | Higher on mimics |
| Rhodobacteraceae | 23 | 13 |
| Flavobacteriaceae | 8 | 8 |
| Saprospiraceae | 16 | 0 |
| Granulosicoccaceae | 4 | 2 |
| Alteromonadaceae | 5 | 0 |

| Roots |  |  |  |  |  |  |
| --- | --- | --- | --- | --- | --- | --- |
| Family | Higher on roots | Higher on mimics | Higher on mimics | Higher on sediment | Higher on roots | Higher on sediment |
| Desulfocapsaceae | 40 | 0 | 8 | 25 | 27 | 9 |
| Flavobacteriaceae | 24 | 13 | 19 | 20 | 21 | 17 |
| Desulfosarcinaceae | 34 | 0 | 0 | 35 | 8 | 26 |
| Bacteroidetes_BD2-2 | 31 | 1 | 0 | 25 | 16 | 14 |
| Spirochaetaceae | 23 | 0 | 1 | 21 | 15 | 11 |

When we examined predicted pathways that changed between the leaf and leaf mimic microbiomes, we identified 53 pathways that changed, 16 upregulated in leaf microbiomes and 37 upregulated on mimic microbiomes (Supplemental Table 4). These predicted pathways included differences in amino acid degradation, starch degradation, and denitrification; however, these predicted pathways did not always indicate expected differences among plants and mimic communities or produce clear candidate predicted pathways, likely at least in part due to limits in prediction of environmental microbial pathways, so we have focused on taxonomic differences. *Roots*

Despite strong compositional differences between roots and root mimics (PERMANOVA, F = 31.511, p = 0.001, r^2^ = 0.307; Fig 3a), these substrates did not differ in ASV richness (negative binomial glm with crossed random effects for site and month, p = 0.28, Fig 1a) or variance (betadisper ANOVA, p = 0.29). When including sediments in the comparisons, richness did not differ among the three groups in richness (negative binomial glm with crossed random effects for site and month, p = 0.31) but microbiome variance among samples was less among sediment samples than either of the other two groups (betadisper ANOVA, p < 0.001, Tukey’s HSD sediment vs mimic p = 0.0001, vs roots p = 0.004); ASV composition on roots, mimics and sediment were compositionally distinct (PERMANOVA r^2^ = 0.3480348, F = 37.90152, p = 0.001, see Supplemental Table 5 for pairwise comparisons). When we examine overlap in predicted Metacyc pathways, we found that there was a significant difference between roots, sediments and mimics (PERMANOVA, r^2^ = 0.18903, F = 12.534, p = 0.001), though this effect was weaker than for the sequence based compositional differences (Figure 3b, See Supplemental Table 5 for pairwise differences).

**Figure 3:**
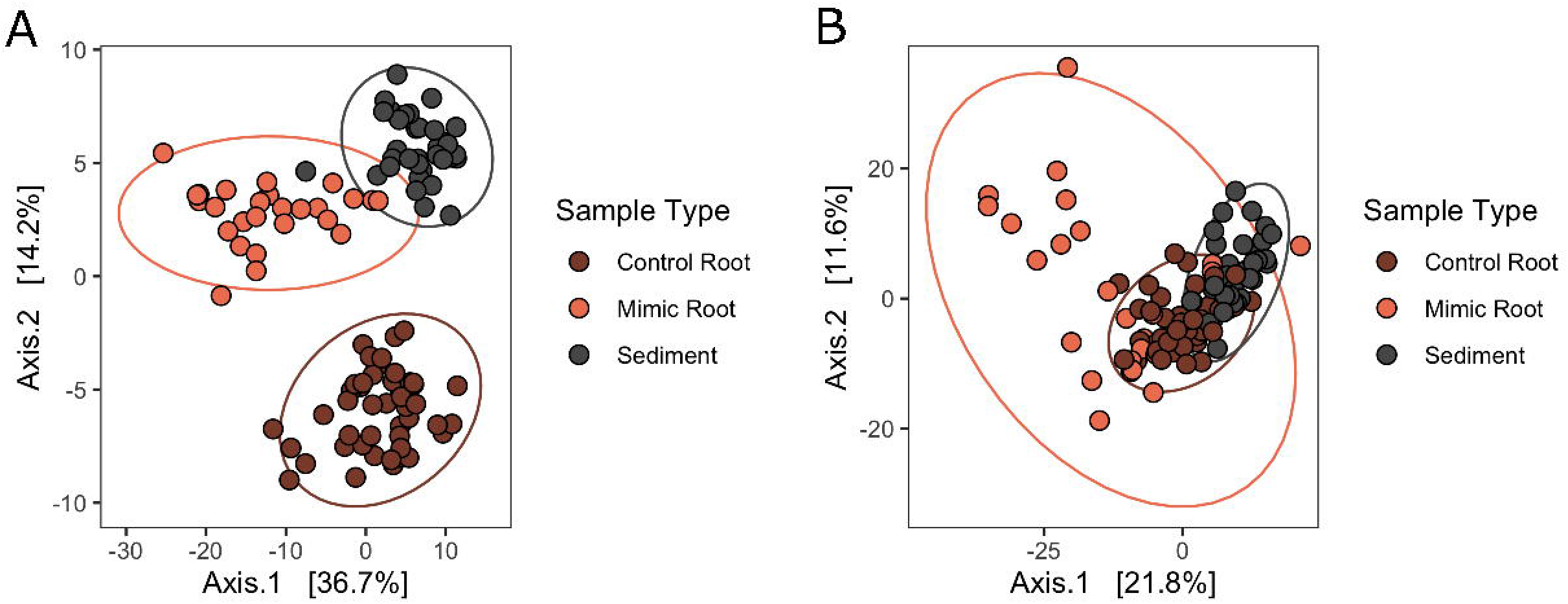
(A) Ordination of bacterial community structure based on principal coordinate analysis of phylogenetic-isometric log-ratio transformed distances. (B) Ordination of predicted Metacyc pathways structure based on principal coordinate analysis of centered log-ratio transformed distances. Red-orange points are communities on root mimics, dark brown points are communities on roots, and grey points are communities in sediments. All communities are distinct from each other in each analysis (PERMANOVA p < 0.001).

When examining core ASVs (present in at least 50% of samples of a given type at at least 1% detection rate), we found that roots and sediments largely harbored distinct bacterial communities, by ASV (Figure 1b), though root mimics had few ASVs unique to its core microbiome (only 2 ASVs unique to root mimics); again, we saw the same patterns when including all ASVs in these analyses, and not just the core (Supplemental Figure 1). Of 256 ASVs in the core root microbiome, 111 or 43% were found only on roots and 181 (71%) were found only on roots and in sediments. Only 52 core root ASVs (20%) were shared between roots and root mimics. When we examined all ASVs (without core restrictions), root mimics had more taxa unique to their sample type, indicating considerable variability in communities assembled on root mimics and a large contribution of rare ASVs (Supp figure 1).

We found many ASVs varied in abundance between these groups (457 between sediments and mimics, 505 between roots and sediments, and 486 between roots and mimics). Of these, the majority were at higher relative abundances on roots or sediments compared to mimics (comparing roots to mimics, 437 were higher on roots, and 49 were higher on mimics; comparing sediment to mimics, 359 were higher in sediments, 98 were higher on mimics; comparing roots to sediments 265 were higher on roots, 240 were higher in sediments; see Table 1 for families with the most representatives, Supplemental Table 6 for more details and Supplemental Table 7 for all ASVs that varied). The families that had the largest number of taxa vary among sample types included Spirochaetaceae (71 ASVs), Bactoroidetes BD2-2 (87 ASVs), Desulfosarcinaceae (103 ASVs), Desusulfocapsaceae (109 ASVs), and Flavobacteriaceae (114 ASVs). Within the families Spirochaetaceae, Bactoroidetes, and Desulfosarcinaceae most ASV were at greater relative abundance on roots than mimics (Table 1). In Flavobacteriaceae, roughly equal numbers of ASVs were more abundant in roots vs mimics vs. sediment. A few families showed an abundance of ASVs on roots compared to both sediments and mimics including: Desulfobacteraceae, Lachnospiraceae, Marinilabiliaceae, Moduliflexaceae, and Prolixibacteraceae (Supplemental Table 6).

While the predicted pathways that varied were numerous and not particularly remarkable (as indicated in Supplemental Table 8), we found that indicated pathways were generally indicated to be upregulated on mimics in pairwise comparisons (133 predicted pathways higher in mimics compared to 32 in sediments, and 137 higher on mimics compared to 35 on roots). Again we anticipate that due to limits in prediction of environmental microbial pathways these may be limited (especially seeing lower numbers of predicted pathways enriched where we saw more taxa enriched), so we have focused on the taxonomic differences we identified.

## Discussion

We found large and consistent differences in the microbiome between seagrass and structural mimics both on above- and belowground surfaces. This builds on previous work that showed distinction between microbiomes on water and leaf surfaces and sediment and root surfaces (Fahimipour *et al*., 2017), showing definitively that microbiomes respond not just to plant physical structure but also the biological activity associated with the host. Previous work also showed strong geographic variation in the microbiome of seagrasses at medium and large scales (Fahimipour *et al*., 2017; Hurtado-McCormick *et al*., 2019) yet, we find consistent differences between mimics and plants across four close sites and three time periods from early to late summer. These findings were consistent for both roots and leaves. This confirms a need for understanding of how these communities are cultivated/built, their interactions with the plants, and ultimate influence on plant fitness.

### Leaves are differentiated from passive mimics

While we saw a >50% overlap in core taxa between the mimics and leaves, we found that they were compositionally distinct (both taxonomically and functionally) and even showed differences in alpha diversity (more ASVs were found on the average mimic than the average leaf). Epiphytic algae also have higher alpha diversity on mimicked compared to natural seagrass and differences in the relative abundance of major microalgal groups between mimic and leaves (Pinckney and Micheli, 1998). These differences in communities between leaves and mimics likely represent either microbial preferences for different surfaces or selection by plants. The reduced alpha diversity on real leaves suggests that there may be some selection by leaves, but also some bacterial preferences as the leaf microbiome is not simply a subset of that on mimics (Fig 1b). While these mimics were not perfect physical mimics, we did find they captured a large portion of the eelgrass leaf community.

This contrasts with mimicked environments from other marine organismal phyllospheres. Recent work in kelp indicated an enrichment of common seawater taxa on artificial substrate (agar infused with and without kelp), no difference in taxonomic diversity on artificial substrates compared to kelp, and increases in aerobic taxa on the surface of kelp blades (Weigel and Pfister, 2021). We found none of these patterns on seagrass and seagrass mimics, instead finding no enrichment for common taxa on seagrass compared to mimics, higher taxonomic diversity on mimics, and no compelling evidence that compositional differences we observed were due to differences in aerobic conditions. The distinction from our study could be driven by the accumulation of epiphytic algal communities on both seagrass leaves and mimics that could render the mimic surfaces highly aerobic; there also is less carbon released by seagrasses than macroalgae (Barrón *et al*., 2014) which likely further distinguishes microbe-host interactions in seagrass from those in kelp.

Terrestrial work mimicking plants has largely focused on even more precisely recreating leaf environments when using artificial surfaces in comparisons (Doan and Leveau, 2015). Use of mimics to test the effect of terrestrial plant leaf morphology on microbiota has indicated that the physical structure plays an important role in influencing the microbiome via moisture retention (Doan *et al*., 2020), a mechanism that is irrelevant to the submerged microbiome of aquatic plant leaves. Like leaves of terrestrial plants, seagrass leaves exude amino acids (Jørgensen *et al*., 1981) and dissolved organic carbon (Wetzel and Penhale, 1979) which create a uniquely rich environment potentially shaping their microbial communities including some predicted amino acid degradation pathways we identified (Supplemental Table 4). These and other potentially seagrass-curated microbial partners could allow us to identify mechanisms in the future that have allowed seagrasses to persist in these extreme environments with a host of new biotic interactions as well ranging from microalgae to marine microbial communities. Future experiments could also investigate interactions involving microbial attachment and facilitation or deterrence by plant exudates as has been explored in terrestrial systems (Zhang *et al*., 2014; Doan and Leveau, 2015; Warning and Datta, 2017) to compare differences that have arisen with transitions to sea.

Finally, while our limited functional evidence does not indicate clear functional differences, the specific taxa that we observe on leaves may have speculatively important roles that are worthy of further investigation. The repeated enrichment of certain ASVs on leaves versus mimics suggests that they might be good targets for experimentation. Some — such as *Kordia spp. —* are likely algicidal bacteria that has been isolated during red tides (Sohn *et al*., 2004) and could be cultivated by seagrass to manage epiphyte loads; we have also previously identified different *Kordia* ASVs at higher relative abundance under warmed and cooled temperature treatments (Schenck et al. submitted). The two Saprospiraceae that we saw multiple representatives within a genera only present on leaves *(Lewinella spp.* and *Rubidimonas spp.)* are both genera comprised of aerobic heterotrophs previously isolated from marine environments that can metabolize complex starches (McIlroy and Nielsen, 2014) and have previously been associated higher levels in macroalgal diseases (Zozaya-Valdés *et al*., 2017). While these bacteria are associated more strongly with seagrass, further investigation is needed to determine how they interact with seagrass, seagrass diseases and seagrass epiphytes -- as they may be cultivated partners in removing unwanted epiphytes or could be negatively affecting eelgrass as well.

### Root microbes are vastly different from those in sediment and on root mimics

That root microbiomes differed from mimics was not surprising given that roots exude both organic (e.g., sugars) and inorganic (e.g., oxygen) compounds that have major influences on microbiota. In fact, sugar concentrations associated with seagrass roots can be exceedingly high yet may not metabolized by bacteria due to the presence of inhibitory phenolic compounds (Sogin *et al*., 2021). Similarly, the combination of oxygen leakage from root tips and surrounding sulfidic sediments promotes sulfur oxidizing bacteria (Brodersen *et al*., 2018; Martin *et al*., 2019). Given these plant-caused environmental differences from surrounding sediments, it does not seem surprising that root mimics microbiomes did not resemble those on roots. However, as in leaves, we captured the surface microbiome of a physical structure buried in sediment. There was no clear “core” of root mimic communities and while there was no evidence of a difference in alpha diversity, our taxonomic analyses showed that taxa that varied between mimics and sediments or roots were generally higher in relative abundance on root surfaces or in sediments than on the mimics. Given the lower abundances, we were surprised to see the opposite result in functional predictions where predicted pathways were generally upregulated on mimics. However, given the more limited functional predictions (including the lack of upregulation sulfate reduction related predicted pathways on roots and sediments compared to mimics despite increases in taxonomic relative abundance of sulfate reducing bacteria) and rapidly changing taxonomy of some of these groups (Waite *et al*., 2020), the bacteria on mimics might be better described than other groups. However, based on taxonomy we have identified several families containing ASVs worthy of further investigation including Desulfobacteraceae and Prolixibacteraceae, both of which were overrepresented on roots compared to mimics and sediments and involved in sulfate-reduction and nitrogen cycling respectively.

Terrestrial experiments investigating rhizosphere microbial communities through creation of artificial environments have added root exudates through capillaries (e.g., addition of oxalic acid into soils frees carbon; Keiluweit *et al*., 2015), and similar experiments could test the roles of exudates and oxygen extrusion independently and in combination to further determine the mechanisms by which microbial communities on the surfaces of roots are assembled, particularly how they sustain these plants in highly anoxic sediments. Based on the taxa we identified, it is likely that seagrass root bacterial communities, like those of terrestrial plants, are structured by a resource exchange between hosts and microbes. While there are many differences in what these resources are and partnerships look like (e.g., eelgrass do not have associations with arbuscular mycorrhizal fungi (Nielsen *et al*., 1999; Ettinger and Eisen, 2019) and land plants live in well oxygenated soils instead of sulfidic anoxic sediments), aquatic and land angiosperms participate in a resource exchange that drives microbial community structures in the rhizosphere (for review of land plants see Jacoby *et al*., 2017). This suggests that it is likely that root microbes are not only affected by the eelgrass, but affect eelgrass itself, perhaps through sulfur metabolism (Fahimipour *et al*., 2017). While no studies have directly tested the effect of root microbiome on seagrass growth, the oxidation of sulfides by lucinid bivalve – bacterial symbiosis has been implicated as a major influence on seagrass success in anoxic sediments (van der Heide *et al*., 2012). Direct tests implicating changing rhizosphere microbial communities with changes in plant performance would be necessary to explicitly test these roles but starting with isolates from many of the families we saw preferentially on seagrass roots compared to mimics and sediments seem likely candidates to have specific adaptations that might deal with both potential phenolic challenges.

### Overall conclusions

We found robust evidence of differences between mimics and plant tissues in various environments in seagrass. These large differences suggest that there is a unique environment created by the plant that creates these distinct communities either as part of active partnerships or through inhibition of certain microbes or unique characteristics of that environment preferred by some microbes. Unlike in terrestrial plant surfaces, water retention is unlikely to play a role in driving microbe assembly in aquatic leaves. Seagrass leaf microbiomes may be structured more like microbiomes in terrestrial and marine root microbiomes, by exudates of plants and their influence on the environment. Further experiments with more detailed or realistic mimics could isolate the mechanisms by which these microbiomes are structured and function.

Finally, despite the differences emphasized here, there is vast overlap between these communities even on simple substrates, especially on leaves and their mimics. This designates a manageable number of taxa to further examine to see what factors are driving their unique assemblies. With a continued and rising interest in microbial communities in general and the role of microbial communities associated with seagrasses specifically, extensions of this work point towards associations that should be further explored for understanding holobiont dynamics across species ranges.

## Experimental Procedures

### Field methods

In July 2015, we deployed artificial substrates to characterize the microbiome of a leaf and root mimics compared with those on live plants (see Supp Figure 2 for site locations). We conducted this experiment at four eelgrass beds within Bodega Harbor within 2 km of each other yet vary in distance from the mouth of the harbor, which sets up a gradient of increasing temperature (2° C mean temperature difference among sites), decreasing water flow, and progressively finer sediment grain size that result in each site harboring distinct eelgrass microbial communities on seagrass that vary among seasons (Kardish and Stachowicz unpublished data). We used 0.75 m long, 4 mm wide green polypropylene ribbons attached to a vexar mesh anchored into sediments to mimic artificial leaf substrates, a standard technique that has been used for over 40 years to mimic the physical habitat provided by eelgrass to isolate the role of physical structure in structuring the epiphytic, epifaunal and fish communities that inhabit eelgrass (Barber *et al*., 1979). These ribbons mimic the physical structure of seagrass leaves in a bed with a similar length, width and accumulation of epiphytes. We also deployed artificial root substrates (4 in. twist ties twisted around the same vexar mesh as ribbons) that went to the depth where most root biomass is found at these sites (approximately 2-5 cm) to examine the community that accumulates on a physical structure at a similar depth in sediment. Neither artificial substrate was preinoculated with microbial communities though they were planted (attached to a vexar mesh) inside a seagrass bed immediately adjacent to live plants. We sampled undisturbed plants near the mimics for comparison. We also sampled plants that had been taken from these sites, taken back to the Bodega Marine lab, attached to the same vexar screens as the mimics and planted back into the field. These results parallel the results presented in the main text comparing mimics and undisturbed plants; parallel analyses can be found in Appendix A. We sampled after one, two and three months after deployment, which is sufficient time for transplanted eelgrass to take on microbial characteristics of a new site (Kardish and Stachowicz unpublished data). For leaves and leaf mimics, we took a 2 cm clip of leaf or ribbon at approximately 15 cm above the sediment surface. For the root mimics, we took a 2 cm clip from the bottom of the twist tie (at approximately the same depth as root samples). For roots, we detached ~10 roots from the rhizome. For sediment samples, we took a small sediment sample from approximately 2 cm under the sediment surface (a similar depth to roots sampled). Samples were immediately placed on dry ice and were frozen at −80° C within a few hours of sampling until extraction, and all instruments were alcohol sterilized in between samples. At each of the four sites, across three timepoints, we sampled three sediment samples, three mimic samples (for each of leaf and root mimics), and four plant samples (leaf and root). Due to the identifiable nature of these sample types, we were unable to blind ourselves to sample type during sampling or extraction.

### Molecular Methods and Bioinformatic analysis

We extracted DNA with the MoBio PowerSoil DNA kit from leaves, roots, and sediments. To get the surface of the leaves and roots only, we vortexed each frozen sample with 500ul of MilliQ water and then added that liquid to the bead tubes and proceeded with the standard extraction protocol (full protocol available at github.mkardish/Transplants/Lab_Protocols). For sediments, we added a small amount of sediment (approximately 0.25 mg) directly to the bead tube. We amplified and sequenced the V4-V5 region of the 16S rRNA gene on an Illumina MiSeq to identify bacteria present at the Integrated Microbiome Resource at Dalhousie University with primers 515F and 926R (Walters *et al*., 2016; Comeau *et al*., 2017).

### Bioinformatic Analysis

We ran all bioinformatic and statistical analyses in R (version 4.0.3). We used a standard dada2 pipeline to error check our reads and to identify amplicon sequence variants (Callahan *et al.*, 2016). We used only forward reads in our subsequent analyses (280 base pairs). We identified ASV taxonomy based on the SILVA database (Quast *et al*., 2013) and built a phylogeny of ASVs using alignments built with DECIPHER (Wright, 2015) then a tree built with FastTree2 (Price *et al*., 2010) then converted to ultrametric (Britton *et al*., 2007). We then rooted the bacterial tree with an archaeal outgroup (Callahan *et al*., 2016).

We also examined the functional potential of the metagenomes of our samples using PICRUSt2 (Douglas *et al*., 2020). While these predictions come with major caveats for environmental samples due to underrepresentation in the database, we used this approach to infer potential metabolic pathways based on similarities to known metabolisms and compare these among tissue types.

### Sampling and sequencing success

We identified 7,696 bacterial ASVs across 192 leaf, root, mimic and sediment samples after quality filtering samples to 3,752,142 reads. Root samples contained between 390 and 1, 013 bacterial ASVs on their surface (we measured 47 root samples with read depth between 11, 522 and 49,735 reads), Root mimics samples contained between 81 and 790 bacterial ASVs on their surface (we measured 26 root mimic samples with read depth between 2,037 and 37,793 reads), Sediment samples contained between 270 and 843 bacterial ASVs on their surface (we measured 36 sediment root samples with read depth between 10,771 and 40,343 reads), leaf samples contained between 191 and 841 bacterial ASVs on their surface (we measured 48 leaf samples with read depth between 5,961 and 61,896 reads) and leaf mimic samples which contained between 195 and 717 bacterial ASVs (35 leaf mimic samples with between 4,587 and 31,648 reads per sample)

### Statistics

We analyzed the compositional changes in our dataset based on phylogenetic similarity among samples by normalizing samples via a phylogenetic isometric log transform described in (Silverman *et al*., 2017) and implemented in the R-package “philr”. This allows a compositional transformation of the phylogenetic data -- comparing differential weights at nodes throughout the bacterial tree as opposed to just ASVs. We then calculated the Euclidean distance among samples before using PERMANOVA to determine differences among sample types controlling for month and site by constraining permutations. We tested homogeneity of group dispersions with the betadispr function in ‘vegan’.

To measure bacterial richness, we rarified all samples to 2950 reads samples which we repeated 200 times (McMurdie and Holmes, 2014) and used each sample’s average “Observed ASVs” in our analysis as our measure of bacterial richness in a sample. We tested differences in Observed ASVs using the negative binomial mixed model with crossed random effects implemented in lme4: Observed ASVs ~ Sample Type + (Sample Type | Site)+(Sample Type | Month) (Bates *et al*., 2015). We also visualized overlap in these observed ASVs using upSet which allowed us to identify the numbers of overlapping and non-overlapping ASVs across sample types (Conway *et al*., 2017).

To identify which ASVs varied between samples we performed a likelihood ratio test in DESeq2 comparing models of ~ Site + Month + Sample Type with ~ Site + Month (separately for above and belowground samples) after geometric mean centering raw ASV abundances (Love *et al*., 2014). We then examined the contrast between sample types to identify which ASVs varied in each compartment.

We treated functional data compositionally as well, using PERMANOVA to analyze differences in pathway composition among samples after a centered log-ratio transformation. We then used DESeq2 with the same models as for taxonomic differences to test for predicted pathway differences among sample types.

### Data Accessibility

All scripts used to analyze this data are available at www.github.com/mkardish/Mimics and sequences have been deposited under the NCBI BioProject ID PRJNA731931.

## Acknowledgements

We would like to thank J. Eisen and E. Grosholz for their comments on this manuscript. We would like to thank A. Alexiev for assistance with extractions and A. Firl for assistance in sequencing. We would like to thank the Stachowicz Lab for their expertise and assistance in field work. This work was funded by the UC Davis Center for Population Biology, the National Science Foundation Graduate Research Fellowship (to M. Kardish) and a grant from the Gordon and Betty Moore Foundation. This work used the Extreme Science and Engineering Discovery Environment (XSEDE) on the Comet at SDSC through an allocation to M. Kardish TG-DEB160008.

