## Supplemental Figures and Tables for "More than a stick in the mud: Eelgrass leaf and root bacterial communities are distinct from those on physical mimics"

Supplemental Figure 1: Overlap among all ASVs present in each sample type. Diagram is a barplot of shared community memberships, equivalent to a Venn diagram.

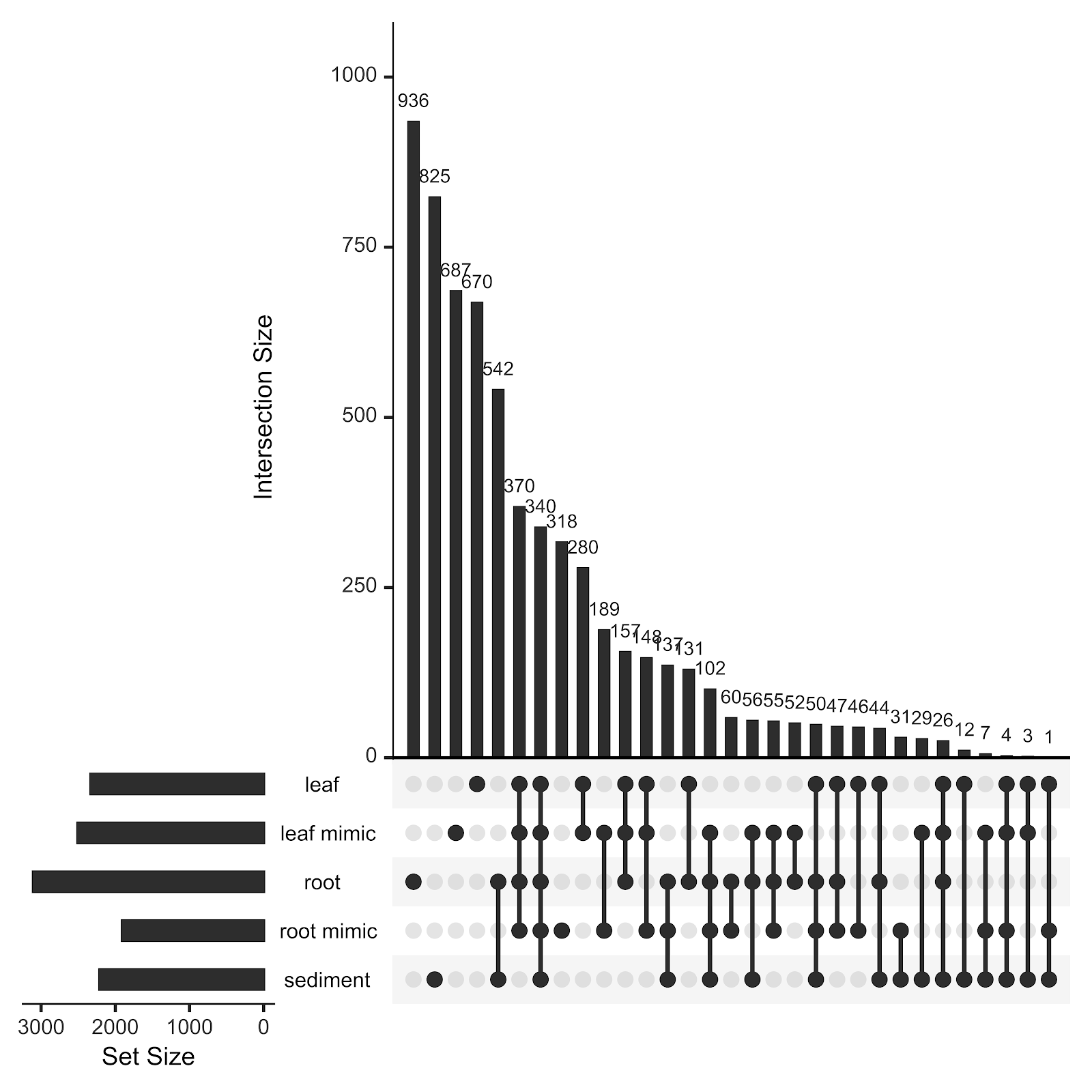

Supplemental Figure 2: Map of Sampling sites in Bodega Harbor, Bodega Bay, CA, USA.

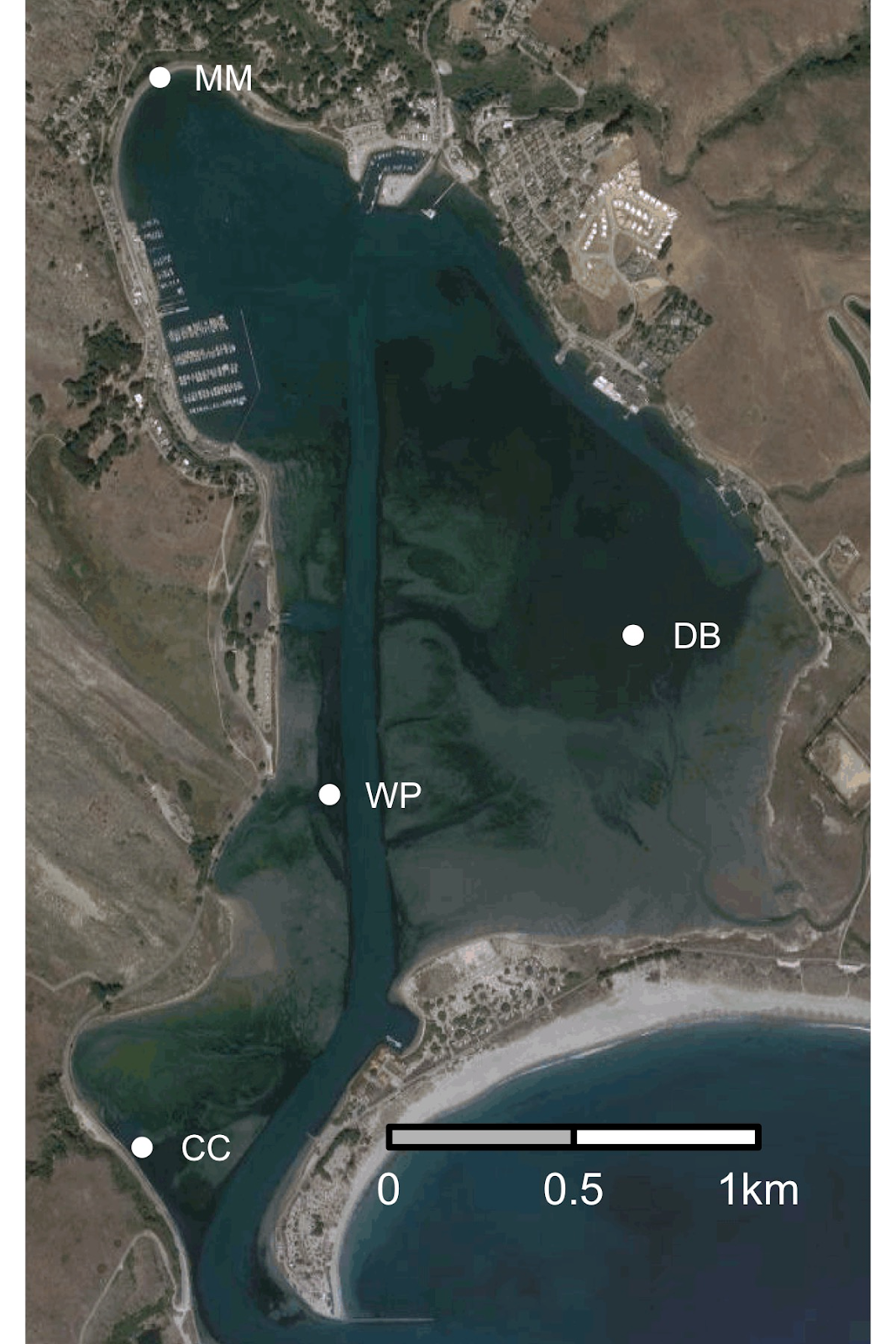

**Supplemental Tables legends (followed by supplemental tables themselves)**

Supplemental Table 1: For leaf bacterial communities, the family-level identification of ASVs that varied significantly between mimics and seagrass substrate determined by DESeq2.

Supplemental Table 2:  For leaf bacterial communities, the genus-level identification of ASVs that varied significantly between mimics and seagrass substrate determined by DESeq2.

Supplemental Table 3: For leaf bacterial communities, all ASVs that varied significantly between mimics and seagrass substrate determined by DESeq2, including magnitude of differences.

Supplemental Table 4:  For leaf bacterial communities, all Metacyc predicted pathways that varied significantly between mimics and seagrass substrate determined by DESeq2, including magnitude of differences.

Supplemental Table 5: Results of pairwise PERMANOVA tests distinguishing compositional differences among roots, root mimics, and sediments in both ASV composition and composition of predicted Metacyc pathways.

Supplemental Table 6: For belowground bacterial communities, the family-level identification of ASVs that varied significantly  among mimics, seagrass and sediment determined by DESeq2.

Supplemental Table 7: For belowground bacterial communities, all ASVs that varied significantly among mimics, seagrass and sediment determined by DESeq2, including magnitude of differences.

Supplemental Table 8:  For belowground bacterial communities, all Metacyc predicted pathways that varied significantly among mimics, seagrass and sediment determined by DESeq2, including magnitude of differences.

**Tables:**

**Supplemental Table 1**

| **Family** | **Higher on leaves** | **Higher on mimics** |
| --- | --- | --- |
| Alteromonadaceae | 5 | 0 |
| Arenicellaceae | 1 | 1 |
| Blastocatellaceae | 0 | 1 |
| Cellvibrionaceae | 1 | 0 |
| Colwelliaceae | 2 | 0 |
| Crocinitomicaceae | 5 | 0 |
| Cryomorphaceae | 3 | 0 |
| Desulfocapsaceae | 0 | 1 |
| DEV007 | 0 | 2 |
| Flavobacteriaceae | 8 | 8 |
| Fokiniaceae | 1 | 0 |
| Gimesiaceae | 0 | 1 |
| Granulosicoccaceae | 4 | 2 |
| Hyphomicrobiaceae | 0 | 1 |
| Hyphomonadaceae | 2 | 1 |
| Kangiellaceae | 1 | 0 |
| Marinomonadaceae | 1 | 0 |
| Methylophagaceae | 1 | 0 |
| Methylophilaceae | 3 | 1 |
| Micavibrionaceae | 0 | 2 |
| Microtrichaceae | 0 | 1 |
| Nitrincolaceae | 2 | 0 |
| NS9_marine_group | 1 | 0 |
| Oleiphilaceae | 1 | 0 |
| Phormidesmiaceae | 0 | 1 |
| Pirellulaceae | 4 | 1 |
| Rhizobiaceae | 1 | 2 |
| Rhodobacteraceae | 23 | 13 |
| Rhodothermaceae | 1 | 0 |
| Rickettsiaceae | 1 | 0 |
| Rubinisphaeraceae | 0 | 2 |
| Rubritaleaceae | 0 | 1 |
| Saprospiraceae | 16 | 0 |
| Sphingomonadaceae | 0 | 2 |
| Spirosomaceae | 1 | 0 |
| Spongiibacteraceae | 1 | 0 |
| Sulfurovaceae | 0 | 2 |
| Terasakiellaceae | 1 | 0 |
| Thiomicrospiraceae | 1 | 0 |
| Trueperaceae | 0 | 1 |
| Unknown_Family | 0 | 1 |
| Woeseiaceae | 0 | 1 |

Supplemental Table 2

| **Family** | **Genus** | **Higher on leaves** | **Higher on mimics** |
| --- | --- | --- | --- |
| Alteromonadaceae | Glaciecola | 4 | 0 |
| Alteromonadaceae | Salinimonas | 1 | 0 |
| Arenicellaceae | Arenicella | 1 | 1 |
| Blastocatellaceae | Blastocatella | 0 | 1 |
| Cellvibrionaceae | Agaribacterium | 1 | 0 |
| Colwelliaceae | Colwellia | 2 | 0 |
| Crocinitomicaceae | Crocinitomix | 1 | 0 |
| Crocinitomicaceae | Fluviicola | 3 | 0 |
| Cryomorphaceae | Vicingus | 1 | 0 |
| Flavobacteriaceae | Aquibacter | 0 | 1 |
| Flavobacteriaceae | Aurantivirga | 1 | 0 |
| Flavobacteriaceae | Changchengzhania | 1 | 0 |
| Flavobacteriaceae | Kordia | 3 | 0 |
| Flavobacteriaceae | Maribacter | 0 | 1 |
| Flavobacteriaceae | Polaribacter | 1 | 0 |
| Flavobacteriaceae | Psychroserpens | 0 | 1 |
| Flavobacteriaceae | Ulvibacter | 2 | 2 |
| Fokiniaceae | MD3-55 | 1 | 0 |
| Granulosicoccaceae | Granulosicoccus | 4 | 2 |
| Hyphomicrobiaceae | Filomicrobium | 0 | 1 |
| Hyphomonadaceae | Hellea | 1 | 0 |
| Hyphomonadaceae | Hyphomonas | 0 | 1 |
| Hyphomonadaceae | Litorimonas | 1 | 0 |
| Marinomonadaceae | Marinomonas | 1 | 0 |
| Methylophilaceae | Methylotenera | 3 | 1 |
| Microtrichaceae | Sva0996_marine_group | 0 | 1 |
| Oleiphilaceae | Oleiphilus | 1 | 0 |
| Phormidesmiaceae | Phormidesmis_ANT.LACV5.1 | 0 | 1 |
| Pirellulaceae | Blastopirellula | 3 | 1 |
| Pirellulaceae | Rhodopirellula | 1 | 0 |
| Rhizobiaceae | Pseudahrensia | 1 | 2 |
| Rhodobacteraceae | Celeribacter | 0 | 1 |
| Rhodobacteraceae | Jannaschia | 0 | 1 |
| Rhodobacteraceae | Octadecabacter | 1 | 1 |
| Rhodobacteraceae | Phaeobacter | 0 | 1 |
| Rhodobacteraceae | Roseovarius | 0 | 1 |
| Rhodobacteraceae | Sedimentitalea | 1 | 1 |
| Rhodobacteraceae | Sulfitobacter | 0 | 1 |
| Rhodobacteraceae | Tateyamaria | 1 | 1 |
| Rhodobacteraceae | Thiobacimonas | 0 | 1 |
| Rhodobacteraceae | Yoonia-Loktanella | 2 | 0 |
| Rickettsiaceae | Candidatus_Megaira | 1 | 0 |
| Rubinisphaeraceae | Planctomicrobium | 0 | 1 |
| Rubritaleaceae | Persicirhabdus | 0 | 1 |
| Saprospiraceae | Lewinella | 3 | 0 |
| Saprospiraceae | Phaeodactylibacter | 1 | 0 |
| Saprospiraceae | Portibacter | 1 | 0 |
| Saprospiraceae | Rubidimonas | 2 | 0 |
| Sphingomonadaceae | Parasphingopyxis | 0 | 1 |
| Spirosomaceae | Taeseokella | 1 | 0 |
| Sulfurovaceae | Sulfurovum | 0 | 2 |
| Thiomicrospiraceae | endosymbionts | 1 | 0 |
| Trueperaceae | Truepera | 0 | 1 |
| Woeseiaceae | Woeseia | 0 | 1 |

Supplemental table 3:

| **Family** | **Genus** | **Species** | **baseMean** | **log2FoldChange** | **lfcSE** | **stat** | **pvalue** | **padj** |
| --- | --- | --- | --- | --- | --- | --- | --- | --- |
| Blastocatellaceae | Blastocatella | NA | 8.634 | -3.969 | 0.953 | 13.066 | <0.001 | 0.025 |
| Pirellulaceae | Rhodopirellula | NA | 12.059 | 3.583 | 0.763 | 17.452 | <0.001 | 0.004 |
| Pirellulaceae | Blastopirellula | NA | 26.01 | 2.218 | 0.453 | 16.77 | <0.001 | 0.005 |
| Pirellulaceae | Blastopirellula | NA | 6.362 | -3.07 | 0.656 | 16.339 | <0.001 | 0.006 |
| Pirellulaceae | Blastopirellula | NA | 21.227 | 2.555 | 0.594 | 14.095 | <0.001 | 0.016 |
| Phormidesmiaceae | Phormidesmis_ANT.LACV5.1 | NA | 58.235 | -2.408 | 0.557 | 12.483 | <0.001 | 0.031 |
| Rubinisphaeraceae | Planctomicrobium | NA | 8.024 | -4.426 | 0.76 | 21.124 | <0.001 | 0.001 |
| Rubinisphaeraceae | NA | NA | 29.24 | -1.711 | 0.341 | 21.782 | <0.001 | <0.001 |
| Gimesiaceae | NA | NA | 8.264 | -4.048 | 0.964 | 11.668 | 0.001 | 0.045 |
| Pirellulaceae | Blastopirellula | NA | 36.821 | 8.175 | 0.608 | 116.911 | <0.001 | <0.001 |
| Sulfurovaceae | Sulfurovum | NA | 35.1 | -2.351 | 0.432 | 26.554 | <0.001 | <0.001 |
| Sulfurovaceae | Sulfurovum | NA | 25.418 | -2.611 | 0.594 | 17.23 | <0.001 | 0.004 |
| Rubritaleaceae | Persicirhabdus | NA | 17.136 | -3.59 | 0.674 | 16.305 | <0.001 | 0.006 |
| DEV007 | NA | NA | 2.813 | -4.186 | 1.106 | 12.838 | <0.001 | 0.027 |
| DEV007 | NA | NA | 10.793 | -4.078 | 0.752 | 15.57 | <0.001 | 0.008 |
| Granulosicoccaceae | Granulosicoccus | NA | 28.28 | 7.429 | 0.607 | 97.66 | <0.001 | <0.001 |
| Granulosicoccaceae | Granulosicoccus | coccoides | 58.104 | 2.262 | 0.45 | 17.754 | <0.001 | 0.003 |
| Granulosicoccaceae | Granulosicoccus | NA | 22.483 | 4.112 | 0.789 | 18.559 | <0.001 | 0.002 |
| Granulosicoccaceae | Granulosicoccus | NA | 105.233 | -1.658 | 0.342 | 17.461 | <0.001 | 0.004 |
| Granulosicoccaceae | Granulosicoccus | NA | 119.671 | 2.337 | 0.413 | 22.214 | <0.001 | <0.001 |
| Granulosicoccaceae | Granulosicoccus | NA | 14.532 | -4.847 | 1.192 | 15.868 | <0.001 | 0.007 |
| Arenicellaceae | Arenicella | NA | 41.303 | 6.26 | 0.618 | 62.286 | <0.001 | <0.001 |
| Arenicellaceae | Arenicella | NA | 9.769 | -4.871 | 1.062 | 17.243 | <0.001 | 0.004 |
| Woeseiaceae | Woeseia | NA | 6.946 | -5.372 | 1.229 | 11.785 | 0.001 | 0.043 |
| Spongiibacteraceae | NA | NA | 58.424 | 3.862 | 0.537 | 34.029 | <0.001 | <0.001 |
| Unknown_Family | NA | NA | 53.935 | -1.182 | 0.296 | 13.938 | <0.001 | 0.017 |
| Methylophilaceae | Methylotenera | NA | 19.786 | -3.236 | 0.849 | 13.234 | <0.001 | 0.023 |
| Methylophilaceae | Methylotenera | NA | 61.362 | 5.868 | 0.586 | 62.869 | <0.001 | <0.001 |
| Methylophilaceae | Methylotenera | NA | 13.359 | 5.89 | 1.13 | 16.955 | <0.001 | 0.005 |
| Methylophilaceae | Methylotenera | NA | 83.462 | 5.768 | 0.62 | 50.412 | <0.001 | <0.001 |
| Alteromonadaceae | Salinimonas | NA | 13.51 | 4.726 | 1.031 | 12.954 | <0.001 | 0.026 |
| Alteromonadaceae | Glaciecola | NA | 11.982 | 5.014 | 1.081 | 15.418 | <0.001 | 0.009 |
| Alteromonadaceae | Glaciecola | NA | 17.041 | 4.317 | 0.903 | 11.855 | 0.001 | 0.042 |
| Alteromonadaceae | Glaciecola | NA | 37.762 | 6.247 | 0.629 | 61.608 | <0.001 | <0.001 |
| Alteromonadaceae | Glaciecola | punicea | 13.312 | 4.5 | 0.919 | 12.536 | <0.001 | 0.031 |
| Methylophagaceae | NA | NA | 8.874 | 5.272 | 1.02 | 19.437 | <0.001 | 0.001 |
| Colwelliaceae | Colwellia | polaris | 58.733 | 6.264 | 0.683 | 45.932 | <0.001 | <0.001 |
| Colwelliaceae | Colwellia | NA | 38.304 | 6.45 | 0.695 | 53.797 | <0.001 | <0.001 |
| Kangiellaceae | NA | NA | 22.289 | 4.652 | 0.662 | 27.473 | <0.001 | <0.001 |
| Marinomonadaceae | Marinomonas | NA | 15.231 | 5.285 | 0.987 | 16.701 | <0.001 | 0.005 |
| Nitrincolaceae | NA | NA | 17.265 | 7.078 | 1.149 | 24.056 | <0.001 | <0.001 |
| Nitrincolaceae | NA | NA | 8.395 | 4.688 | 1.013 | 14.109 | <0.001 | 0.016 |
| Oleiphilaceae | Oleiphilus | NA | 16.753 | 7.231 | 0.661 | 83.582 | <0.001 | <0.001 |
| Cellvibrionaceae | Agaribacterium | NA | 11.934 | 4.209 | 0.84 | 17.029 | <0.001 | 0.004 |
| Thiomicrospiraceae | endosymbionts | NA | 5.656 | 5.261 | 1.24 | 13.717 | <0.001 | 0.019 |
| Trueperaceae | Truepera | NA | 9.138 | -2.689 | 0.554 | 17.499 | <0.001 | 0.004 |
| Flavobacteriaceae | NA | NA | 20.98 | -5.507 | 0.911 | 16.74 | <0.001 | 0.005 |
| Flavobacteriaceae | Psychroserpens | damuponensis | 159.42 | -1.161 | 0.294 | 14.13 | <0.001 | 0.016 |
| Flavobacteriaceae | NA | NA | 84.944 | -1.752 | 0.411 | 16.497 | <0.001 | 0.006 |
| Flavobacteriaceae | Aquibacter | NA | 136.29 | -1.432 | 0.311 | 19.408 | <0.001 | 0.001 |
| Flavobacteriaceae | Ulvibacter | NA | 38.294 | 4.716 | 0.778 | 28.698 | <0.001 | <0.001 |
| Flavobacteriaceae | Changchengzhania | NA | 56.506 | 2.425 | 0.504 | 15.731 | <0.001 | 0.008 |
| Flavobacteriaceae | Ulvibacter | NA | 41.038 | 5.34 | 0.749 | 22.334 | <0.001 | <0.001 |
| Flavobacteriaceae | Ulvibacter | NA | 32.722 | -7.339 | 1 | 32.877 | <0.001 | <0.001 |
| Flavobacteriaceae | Ulvibacter | NA | 16.79 | -7.85 | 0.869 | 17.871 | <0.001 | 0.003 |
| Flavobacteriaceae | Kordia | NA | 29.596 | 6.553 | 1.169 | 15.66 | <0.001 | 0.008 |
| Flavobacteriaceae | Kordia | jejudonensis | 84.147 | 5.963 | 0.578 | 66.903 | <0.001 | <0.001 |
| Flavobacteriaceae | Kordia | NA | 6.941 | 4.927 | 1.165 | 12.53 | <0.001 | 0.031 |
| Flavobacteriaceae | Polaribacter | NA | 126.711 | 8.478 | 0.705 | 79.666 | <0.001 | <0.001 |
| Flavobacteriaceae | Aurantivirga | NA | 28.861 | 9.283 | 1.085 | 16.011 | <0.001 | 0.007 |
| Crocinitomicaceae | Crocinitomix | NA | 19.245 | 6.306 | 0.883 | 27.094 | <0.001 | <0.001 |
| Saprospiraceae | Phaeodactylibacter | NA | 5.174 | 4.593 | 1.116 | 13.229 | <0.001 | 0.023 |
| Saprospiraceae | NA | NA | 31.823 | 4.771 | 1.005 | 14.881 | <0.001 | 0.011 |
| Saprospiraceae | Lewinella | NA | 7.277 | 4.275 | 0.952 | 13.473 | <0.001 | 0.021 |
| Saprospiraceae | Lewinella | NA | 12.896 | 3.2 | 0.82 | 12.708 | <0.001 | 0.029 |
| Saprospiraceae | Lewinella | persica | 23.037 | 3.204 | 0.555 | 21.879 | <0.001 | <0.001 |
| Saprospiraceae | NA | NA | 7.088 | 5.254 | 1.081 | 17.731 | <0.001 | 0.003 |
| Saprospiraceae | NA | NA | 4.064 | 3.836 | 1.055 | 11.465 | 0.001 | 0.05 |
| Saprospiraceae | Portibacter | NA | 34.143 | 2.237 | 0.531 | 13.556 | <0.001 | 0.021 |
| Saprospiraceae | NA | NA | 14.664 | 5.055 | 0.685 | 41.695 | <0.001 | <0.001 |
| Saprospiraceae | NA | NA | 40.698 | 2.5 | 0.581 | 12.285 | <0.001 | 0.034 |
| Saprospiraceae | NA | NA | 5.978 | 4.909 | 1.177 | 12.193 | <0.001 | 0.035 |
| Saprospiraceae | NA | NA | 19.903 | 5.377 | 0.865 | 18.326 | <0.001 | 0.003 |
| Saprospiraceae | NA | NA | 16.506 | 6.402 | 0.908 | 33.039 | <0.001 | <0.001 |
| Saprospiraceae | Rubidimonas | NA | 46.119 | 5.803 | 0.735 | 32.33 | <0.001 | <0.001 |
| Saprospiraceae | Rubidimonas | NA | 16.484 | 6.004 | 0.837 | 34.507 | <0.001 | <0.001 |
| Cryomorphaceae | Vicingus | NA | 39.92 | 8.815 | 1.106 | 15.058 | <0.001 | 0.01 |
| NS9_marine_group | NA | NA | 7.303 | 5.443 | 1.099 | 14.389 | <0.001 | 0.014 |
| Crocinitomicaceae | NA | NA | 11.541 | 5.341 | 1.069 | 15.945 | <0.001 | 0.007 |
| Flavobacteriaceae | NA | NA | 4.78 | -4.952 | 0.905 | 25.497 | <0.001 | <0.001 |
| Flavobacteriaceae | Maribacter | NA | 60.234 | -1.936 | 0.44 | 12.808 | <0.001 | 0.027 |
| Cryomorphaceae | NA | NA | 11.39 | 2.924 | 0.771 | 12.212 | <0.001 | 0.035 |
| Crocinitomicaceae | Fluviicola | NA | 8.382 | 4.419 | 0.965 | 15.854 | <0.001 | 0.007 |
| Crocinitomicaceae | Fluviicola | NA | 59.221 | 4.42 | 0.631 | 30.869 | <0.001 | <0.001 |
| Crocinitomicaceae | Fluviicola | NA | 13.2 | 5.254 | 1.068 | 13.396 | <0.001 | 0.022 |
| Cryomorphaceae | NA | NA | 10.5 | 4.733 | 0.759 | 27.193 | <0.001 | <0.001 |
| Microtrichaceae | Sva0996_marine_group | NA | 6.494 | -4.211 | 1.047 | 11.61 | 0.001 | 0.046 |
| Terasakiellaceae | NA | NA | 20.178 | 5.687 | 0.858 | 29.384 | <0.001 | <0.001 |
| Rhodobacteraceae | Thiobacimonas | NA | 23.578 | -1.724 | 0.454 | 13.363 | <0.001 | 0.022 |
| Hyphomicrobiaceae | Filomicrobium | NA | 14.015 | -3.249 | 0.556 | 22.389 | <0.001 | <0.001 |
| Hyphomonadaceae | Hyphomonas | NA | 6.878 | -3.616 | 0.796 | 13.309 | <0.001 | 0.023 |
| Hyphomonadaceae | Hellea | balneolensis | 32.065 | 3.115 | 0.42 | 42.073 | <0.001 | <0.001 |
| Hyphomonadaceae | Litorimonas | NA | 19.341 | 3.782 | 0.691 | 19.64 | <0.001 | 0.001 |
| Rhodobacteraceae | NA | NA | 13.374 | 3.927 | 0.774 | 17.581 | <0.001 | 0.004 |
| Rhodobacteraceae | NA | NA | 156.8 | 5.239 | 0.369 | 116.098 | <0.001 | <0.001 |
| Rhodobacteraceae | NA | NA | 26.224 | 6.988 | 0.746 | 35.258 | <0.001 | <0.001 |
| Rhodobacteraceae | NA | NA | 49.826 | 5.501 | 0.611 | 50.96 | <0.001 | <0.001 |
| Rhodobacteraceae | Jannaschia | NA | 18.178 | -6.647 | 0.856 | 34.293 | <0.001 | <0.001 |
| Rhodobacteraceae | NA | NA | 11.039 | -5.772 | 0.912 | 12.84 | <0.001 | 0.027 |
| Rhodobacteraceae | Octadecabacter | NA | 134.373 | 4.597 | 0.722 | 24.126 | <0.001 | <0.001 |
| Rhodobacteraceae | Octadecabacter | NA | 182.503 | -1.684 | 0.26 | 34.455 | <0.001 | <0.001 |
| Rhodobacteraceae | Celeribacter | NA | 45.875 | -2.037 | 0.391 | 23.139 | <0.001 | <0.001 |
| Rhodobacteraceae | NA | NA | 28.868 | 3.607 | 0.712 | 15.82 | <0.001 | 0.007 |
| Rhodobacteraceae | NA | NA | 297.305 | 1.386 | 0.23 | 31.552 | <0.001 | <0.001 |
| Rhodobacteraceae | NA | NA | 23.106 | 7.592 | 0.722 | 71.166 | <0.001 | <0.001 |
| Rhodobacteraceae | NA | NA | 70.863 | 2.018 | 0.396 | 20.735 | <0.001 | 0.001 |
| Rhodobacteraceae | NA | NA | 14.836 | 6.254 | 1.052 | 20.536 | <0.001 | 0.001 |
| Rhodobacteraceae | Sulfitobacter | litoralis | 67.491 | -1.478 | 0.35 | 13.263 | <0.001 | 0.023 |
| Rhodobacteraceae | Sedimentitalea | NA | 30.39 | -5.095 | 0.721 | 32.182 | <0.001 | <0.001 |
| Rhodobacteraceae | Phaeobacter | NA | 17.936 | -6.293 | 0.781 | 22.546 | <0.001 | <0.001 |
| Rhodobacteraceae | NA | NA | 26.276 | 5.002 | 0.747 | 23.799 | <0.001 | <0.001 |
| Rhodobacteraceae | Roseovarius | aestuarii | 8.958 | -5.594 | 1.01 | 24.136 | <0.001 | <0.001 |
| Rhodobacteraceae | NA | NA | 9.473 | 5.571 | 1.118 | 13.972 | <0.001 | 0.017 |
| Rhodobacteraceae | NA | NA | 23.751 | 3.868 | 0.534 | 39.091 | <0.001 | <0.001 |
| Rhodobacteraceae | Sedimentitalea | NA | 55.962 | 3.298 | 0.457 | 35.005 | <0.001 | <0.001 |
| Rhodobacteraceae | NA | NA | 6.103 | -4.458 | 0.794 | 26.924 | <0.001 | <0.001 |
| Rhodobacteraceae | NA | NA | 27.698 | 4.835 | 0.552 | 48.63 | <0.001 | <0.001 |
| Rhodobacteraceae | NA | NA | 54.699 | 2.74 | 0.433 | 30.576 | <0.001 | <0.001 |
| Rhodobacteraceae | NA | NA | 22.331 | 6.23 | 1.218 | 14.552 | <0.001 | 0.013 |
| Rhodobacteraceae | NA | NA | 38.101 | 2.069 | 0.45 | 13.874 | <0.001 | 0.018 |
| Rhodobacteraceae | Yoonia-Loktanella | NA | 28.841 | 2.622 | 0.596 | 12.907 | <0.001 | 0.027 |
| Rhodobacteraceae | Yoonia-Loktanella | NA | 33.033 | 1.722 | 0.361 | 19.916 | <0.001 | 0.001 |
| Rhodobacteraceae | NA | NA | 91.674 | 2.711 | 0.319 | 56.853 | <0.001 | <0.001 |
| Rhizobiaceae | Pseudahrensia | NA | 16.213 | 3.444 | 0.764 | 12.287 | <0.001 | 0.034 |
| Rhizobiaceae | Pseudahrensia | NA | 20.369 | -6.875 | 0.646 | 82.222 | <0.001 | <0.001 |
| Rhodobacteraceae | NA | NA | 5.136 | -4.7 | 0.936 | 20.204 | <0.001 | 0.001 |
| Rhodobacteraceae | NA | NA | 52.283 | -2.369 | 0.365 | 34.517 | <0.001 | <0.001 |
| Rhizobiaceae | Pseudahrensia | NA | 12.006 | -6.025 | 0.694 | 25.431 | <0.001 | <0.001 |
| Sphingomonadaceae | NA | NA | 33.438 | -2.647 | 0.722 | 11.976 | 0.001 | 0.039 |
| Saprospiraceae | NA | NA | 42.309 | 6.654 | 0.899 | 31.117 | <0.001 | <0.001 |
| Spirosomaceae | Taeseokella | NA | 38.22 | 3.031 | 0.441 | 37.073 | <0.001 | <0.001 |
| Rhodothermaceae | NA | NA | 20.236 | 2.904 | 0.477 | 24.9 | <0.001 | <0.001 |
| Desulfocapsaceae | NA | NA | 18.128 | -2.126 | 0.538 | 12.336 | <0.001 | 0.034 |
| Fokiniaceae | MD3-55 | NA | 11.348 | 5.372 | 0.572 | 65.274 | <0.001 | <0.001 |
| Rhodobacteraceae | Tateyamaria | NA | 77.176 | 3.393 | 0.416 | 45.07 | <0.001 | <0.001 |
| Rhodobacteraceae | Tateyamaria | NA | 11.514 | -8.193 | 1.125 | 11.903 | 0.001 | 0.041 |
| Rhodobacteraceae | NA | NA | 23.077 | 4.569 | 0.575 | 40.098 | <0.001 | <0.001 |
| Sphingomonadaceae | Parasphingopyxis | NA | 11.706 | -5.257 | 0.837 | 23.948 | <0.001 | <0.001 |
| Micavibrionaceae | NA | NA | 3.442 | -4.79 | 1.156 | 16.063 | <0.001 | 0.007 |
| Micavibrionaceae | NA | NA | 11.927 | -3.664 | 0.9 | 13.377 | <0.001 | 0.022 |
| Rickettsiaceae | Candidatus_Megaira | NA | 41.926 | 6.587 | 0.791 | 39.498 | <0.001 | <0.001 |

Supplemental Table 4

| **Pathway** | **log2-fold Change** |
| --- | --- |
| nitrifier denitrification | 3.57517042 |
| superpathway of polyamine biosynthesis III | 2.81827993 |
| CMP-pseudaminate biosynthesis | 2.62893542 |
| nylon-6 oligomer degradation | 1.64149213 |
| formaldehyde oxidation I | 1.19448623 |
| formaldehyde assimilation II (RuMP Cycle) | 1.18703389 |
| thiazole biosynthesis II (Bacillus) | 1.17319111 |
| coenzyme M biosynthesis I | 1.06840996 |
| superpathway of thiamin diphosphate biosynthesis II | 0.98492463 |
| methyl ketone biosynthesis | 0.93025842 |
| L-arginine degradation II (AST pathway) | 0.87076592 |
| glucose and glucose-1-phosphate degradation | 0.71012766 |
| ectoine biosynthesis | 0.68464435 |
| norspermidine biosynthesis | 0.59741725 |
| ADP-L-glycero-&beta;-D-manno-heptose biosynthesis | 0.59480138 |
| superpathway of polyamine biosynthesis I | 0.50117322 |
| catechol degradation II (meta-cleavage pathway) | -0.5051274 |
| L-tryptophan degradation XII (Geobacillus) | -0.5311326 |
| catechol degradation I (meta-cleavage pathway) | -0.5785306 |
| acetylene degradation | -0.599038 |
| 2-aminophenol degradation | -0.6086563 |
| catechol degradation to &beta;-ketoadipate | -0.6123384 |
| superpathway of pyridoxal 5'-phosphate biosynthesis and salvage | -0.6160838 |
| superpathway of sulfur oxidation (Acidianus ambivalens) | -0.7083875 |
| reductive acetyl coenzyme A pathway | -0.8598703 |
| meta cleavage pathway of aromatic compounds | -0.8753625 |
| adenosylcobalamin biosynthesis II (late cobalt incorporation) | -0.9123838 |
| androstenedione degradation | -0.9491612 |
| superpathway of salicylate degradation | -0.9500477 |
| methanogenesis from acetate | -0.9597901 |
| catechol degradation III (ortho-cleavage pathway) | -0.968522 |
| aromatic compounds degradation via &beta;-ketoadipate | -0.968522 |
| formaldehyde assimilation I (serine pathway) | -0.9774884 |
| superpathway of 2,3-butanediol biosynthesis | -0.9804931 |
| D-galactarate degradation I | -1.0428591 |
| superpathway of D-glucarate and D-galactarate degradation | -1.0428591 |
| pyruvate fermentation to acetone | -1.0446446 |
| isopropanol biosynthesis | -1.0943399 |
| superpathway of (R,R)-butanediol biosynthesis | -1.1560577 |
| superpathway of L-aspartate and L-asparagine biosynthesis | -1.2056707 |
| glycerol degradation to butanol | -1.2387094 |
| superpathway of N-acetylneuraminate degradation | -1.2987337 |
| superpathway of N-acetylglucosamine, N-acetylmannosamine and N-acetylneuraminate degradation | -1.407782 |
| creatinine degradation II | -1.4875672 |
| D-glucarate degradation I | -1.4888527 |
| 1,5-anhydrofructose degradation | -1.5852009 |
| allantoin degradation to glyoxylate III | -1.6402326 |
| mono-trans, poly-cis decaprenyl phosphate biosynthesis | -1.6413193 |
| cob(II)yrinate a,c-diamide biosynthesis I (early cobalt insertion) | -2.4457325 |
| methylaspartate cycle | -2.4547742 |
| coenzyme B biosynthesis | -2.718622 |
| chondroitin sulfate degradation I (bacterial) | -3.8564041 |
| starch degradation III | -5.3764083 |

Supplemental Table 5

|  |  |  | df | Sum Of Squares | R^2^ | F-Statistic | Pr(>F) |
| --- | --- | --- | --- | --- | --- | --- | --- |
| Based on taxonomy | Root vs Mimic | Sample Type | 1 | 4716.5 | 0.30739 | 31.511 | **0.001** |
|  |  | Residual | 71 | 10627.2 | 0.69261 |  |  |
|  |  | Total | 72 | 15343.7 | 1 |  |  |
|  | Mimic vs. Sediment | Sample Type | 1 | 5659.8 | 0.42429 | 44.22 | **0.001** |
|  |  | Residual | 60 | 7679.5 | 0.57571 |  |  |
|  |  | Total | 61 | 13339.3 | 1 |  |  |
|  | Root vs. Sediment | Sample Type | 1 | 3234.5 | 0.24806 | 26.722 | **0.001** |
|  |  | Residual | 81 | 9804.5 | 0.75194 |  |  |
|  |  | Total | 82 | 13039.1 | 1 |  |  |
| Based on predicted function | Root vs Mimic | Sample Type | 1 | 4991 | 0.11386 | 9.1228 | **0.001** |
|  |  | Residual | 71 | 38841 | 0.88614 |  |  |
|  |  | Total | 72 | 43831 | 1 |  |  |
|  | Mimic vs. Sediment | Sample Type | 1 | 8250 | 0.19562 | 14.592 | **0.001** |
|  |  | Residual | 60 | 33924 | 0.80438 |  |  |
|  |  | Total | 61 | 42174 | 1 |  |  |
|  | Root vs. Sediment | Sample Type | 1 | 4865 | 0.1482 | 14.093 | **0.001** |
|  |  | Residual | 81 | 27963 | 0.8518 |  |  |
|  |  | Total | 82 | 32829 | 1 |  |  |

Supplemental Table 6

| **Family** | **Higher on roots** | **Higher on mimics** | **Higher on mimics** | **Higher on sediment** | **Higher on roots** | **Higher on sediment** |
| --- | --- | --- | --- | --- | --- | --- |
| 4572-13 | 3 | 0 | 0 | 2 | 1 | 2 |
| Acanthopleuribacteraceae | 2 | 0 | 0 | 1 | 1 | 1 |
| Anaerolineaceae | 12 | 0 | 1 | 11 | 3 | 10 |
| Arcobacteraceae | 2 | 0 | 1 | 0 | 2 | 0 |
| Arenicellaceae | 0 | 1 | 0 | 1 | 0 | 1 |
| Bacteroidetes_BD2-2 | 31 | 1 | 0 | 25 | 16 | 14 |
| Calditrichaceae | 10 | 1 | 0 | 11 | 1 | 10 |
| Cellvibrionaceae | 1 | 0 | 1 | 0 | 1 | 0 |
| Christensenellaceae | 3 | 0 | 0 | 3 | 2 | 1 |
| Chromatiaceae | 4 | 0 | 0 | 4 | 0 | 4 |
| Crocinitomicaceae | 1 | 0 | 2 | 0 | 2 | 0 |
| Cyclobacteriaceae | 3 | 0 | 0 | 3 | 0 | 3 |
| Desulfatiglandaceae | 4 | 0 | 0 | 4 | 0 | 4 |
| Desulfobacteraceae | 9 | 0 | 0 | 5 | 7 | 2 |
| Desulfobulbaceae | 8 | 0 | 1 | 7 | 4 | 5 |
| Desulfocapsaceae | 40 | 0 | 8 | 25 | 27 | 9 |
| Desulfococcaceae | 1 | 0 | 0 | 1 | 1 | 0 |
| Desulfolunaceae | 1 | 0 | 0 | 1 | 0 | 1 |
| Desulfomonilaceae | 0 | 1 | 0 | 1 | 0 | 1 |
| Desulfosarcinaceae | 34 | 0 | 0 | 35 | 8 | 26 |
| Desulfovibrionaceae | 3 | 0 | 0 | 1 | 3 | 0 |
| Ectothiorhodospiraceae | 1 | 1 | 0 | 2 | 0 | 2 |
| Fermentibacteraceae | 3 | 0 | 0 | 3 | 1 | 2 |
| Fibrobacteraceae | 1 | 0 | 0 | 1 | 1 | 0 |
| Flavobacteriaceae | 24 | 13 | 19 | 20 | 21 | 17 |
| Fusibacteraceae | 1 | 0 | 0 | 1 | 1 | 0 |
| Gemmatimonadaceae | 1 | 0 | 0 | 1 | 0 | 1 |
| Geopsychrobacteraceae | 2 | 0 | 1 | 0 | 2 | 0 |
| Halieaceae | 9 | 0 | 1 | 9 | 1 | 9 |
| Halomonadaceae | 0 | 2 | 2 | 0 | 2 | 0 |
| Hungateiclostridiaceae | 3 | 0 | 0 | 3 | 1 | 2 |
| Hyphomonadaceae | 0 | 2 | 2 | 0 | 2 | 0 |
| Ignavibacteriaceae | 1 | 0 | 0 | 1 | 0 | 1 |
| Kiritimatiellaceae | 3 | 1 | 0 | 3 | 2 | 1 |
| Lachnospiraceae | 8 | 0 | 3 | 3 | 8 | 0 |
| Latescibacteraceae | 1 | 0 | 0 | 3 | 0 | 3 |
| Lentimicrobiaceae | 7 | 0 | 0 | 7 | 1 | 3 |
| Leptospiraceae | 1 | 0 | 0 | 1 | 1 | 0 |
| Marinifilaceae | 6 | 0 | 0 | 2 | 6 | 0 |
| Marinilabiliaceae | 9 | 0 | 0 | 8 | 6 | 3 |
| Marinomonadaceae | 1 | 0 | 1 | 0 | 1 | 0 |
| Melioribacteraceae | 7 | 0 | 0 | 6 | 3 | 4 |
| Methylophagaceae | 2 | 0 | 1 | 1 | 2 | 0 |
| Methylophilaceae | 1 | 0 | 2 | 0 | 2 | 0 |
| Moduliflexaceae | 19 | 0 | 0 | 14 | 16 | 3 |
| MSBL8 | 5 | 0 | 1 | 4 | 2 | 2 |
| Nitrincolaceae | 1 | 0 | 0 | 1 | 1 | 0 |
| NS11-12_marine_group | 0 | 1 | 1 | 0 | 1 | 0 |
| Pedosphaeraceae | 1 | 0 | 0 | 1 | 0 | 1 |
| PHOS-HE36 | 2 | 0 | 0 | 3 | 0 | 3 |
| Pirellulaceae | 9 | 1 | 2 | 10 | 2 | 9 |
| Prolixibacteraceae | 16 | 0 | 1 | 11 | 10 | 3 |
| Puniceicoccaceae | 2 | 0 | 1 | 0 | 2 | 0 |
| Rhizobiaceae | 4 | 0 | 3 | 0 | 5 | 0 |
| Rhodobacteraceae | 5 | 4 | 17 | 0 | 17 | 0 |
| Rickettsiaceae | 1 | 0 | 1 | 0 | 1 | 0 |
| Rubinisphaeraceae | 0 | 1 | 2 | 0 | 2 | 0 |
| S15A-MN91 | 1 | 0 | 0 | 1 | 1 | 0 |
| Sandaracinaceae | 1 | 0 | 0 | 1 | 0 | 1 |
| Saprospiraceae | 5 | 5 | 6 | 5 | 6 | 3 |
| SB-5 | 11 | 0 | 0 | 8 | 6 | 5 |
| Sedimenticolaceae | 8 | 2 | 1 | 9 | 4 | 6 |
| SG8-4 | 3 | 0 | 0 | 2 | 1 | 2 |
| Shewanellaceae | 0 | 1 | 1 | 0 | 1 | 0 |
| Spirochaetaceae | 23 | 0 | 1 | 21 | 15 | 11 |
| Spirosomaceae | 0 | 1 | 1 | 0 | 1 | 0 |
| Spongiibacteraceae | 2 | 1 | 2 | 1 | 2 | 1 |
| Sulfurimonadaceae | 6 | 1 | 1 | 1 | 6 | 1 |
| Sulfurovaceae | 1 | 0 | 1 | 0 | 1 | 0 |
| Syntrophotaleaceae | 1 | 0 | 0 | 1 | 1 | 0 |
| Thermoanaerobaculaceae | 13 | 0 | 0 | 14 | 0 | 14 |
| Thioalkalispiraceae | 1 | 1 | 0 | 4 | 0 | 4 |
| Thiohalorhabdaceae | 1 | 0 | 0 | 1 | 0 | 1 |
| Thiomicrospiraceae | 7 | 2 | 0 | 9 | 1 | 8 |
| Thiotrichaceae | 2 | 5 | 5 | 4 | 5 | 4 |
| Unknown_Family | 9 | 0 | 1 | 10 | 1 | 10 |
| Vibrionaceae | 1 | 0 | 1 | 0 | 1 | 0 |
| Woeseiaceae | 3 | 0 | 0 | 3 | 0 | 3 |

Supplemental Table 7

See excel file

Supplemental Table 8

| **Pathway** | **Root vs. Mimic** | **Mimic vs. Sediment** | **Root vs. Sediment** |
| --- | --- | --- | --- |
| &beta;-alanine biosynthesis II | -2.707 | 6.111 | 3.404 |
| 1,4-dihydroxy-2-naphthoate biosynthesis I | -0.734 | NA | NA |
| 1,4-dihydroxy-6-naphthoate biosynthesis I | 0.895 | -0.833 | NA |
| 1,4-dihydroxy-6-naphthoate biosynthesis II | 0.936 | -0.922 | NA |
| 2-amino-3-carboxymuconate semialdehyde degradation to 2-oxopentenoate | -1.392 | 2.038 | 0.646 |
| 2-aminophenol degradation | -2.279 | 1.971 | NA |
| 2-methylcitrate cycle I | -1.101 | NA | -0.666 |
| 2-methylcitrate cycle II | -0.814 | NA | -0.5 |
| 2-nitrobenzoate degradation I | -1.351 | 1.919 | 0.568 |
| 3-phenylpropanoate and 3-(3-hydroxyphenyl)propanoate degradation | -1.066 | 3.175 | 2.11 |
| 3-phenylpropanoate degradation | -2.55 | 7.214 | 4.664 |
| 4-coumarate degradation (anaerobic) | NA | 0.894 | 0.511 |
| 4-deoxy-L-threo-hex-4-enopyranuronate degradation | NA | NA | 0.568 |
| 4-hydroxyphenylacetate degradation | -0.697 | 1.697 | 1 |
| 4-methylcatechol degradation (ortho cleavage) | -2.808 | 2.565 | NA |
| acetylene degradation | NA | NA | 0.599 |
| adenosylcobalamin biosynthesis I (early cobalt insertion) | -0.714 | 2.256 | 1.542 |
| adenosylcobalamin biosynthesis II (late cobalt incorporation) | -0.868 | 2.381 | 1.512 |
| ADP-L-glycero-&beta;-D-manno-heptose biosynthesis | 0.625 | -1.079 | NA |
| aerobactin biosynthesis | -0.913 | 2.622 | 1.709 |
| allantoin degradation IV (anaerobic) | -5.19 | 12.389 | 7.199 |
| allantoin degradation to glyoxylate III | -1.345 | 1.089 | NA |
| androstenedione degradation | NA | -1.1 | -1.165 |
| arginine, ornithine and proline interconversion | 0.831 | NA | 0.59 |
| aromatic biogenic amine degradation (bacteria) | -0.658 | 0.859 | NA |
| aromatic compounds degradation via &beta;-ketoadipate | -2.45 | 2.489 | NA |
| benzoyl-CoA degradation II (anaerobic) | 2.335 | -2.511 | NA |
| Bifidobacterium shunt | -1.207 | 1.348 | NA |
| biotin biosynthesis II | -1.18 | 5.052 | 3.871 |
| catechol degradation I (meta-cleavage pathway) | NA | NA | -0.548 |
| catechol degradation III (ortho-cleavage pathway) | -2.45 | 2.489 | NA |
| catechol degradation to &beta;-ketoadipate | -1.833 | 2.268 | NA |
| catechol degradation to 2-oxopent-4-enoate II | -0.686 | 1.26 | 0.574 |
| chitin derivatives degradation | NA | 0.947 | 1.405 |
| chlorophyllide a biosynthesis I (aerobic, light-dependent) | -0.72 | 1.652 | 0.932 |
| chlorophyllide a biosynthesis II (anaerobic) | -0.768 | 1.592 | 0.823 |
| chlorophyllide a biosynthesis III (aerobic, light independent) | -0.768 | 1.592 | 0.823 |
| chlorosalicylate degradation | -3.061 | 5.32 | 2.259 |
| chondroitin sulfate degradation I (bacterial) | -1.706 | 2.696 | 0.99 |
| CMP-legionaminate biosynthesis I | 0.934 | -1.577 | -0.643 |
| CMP-pseudaminate biosynthesis | 2.677 | 3.179 | 5.856 |
| cob(II)yrinate a,c-diamide biosynthesis I (early cobalt insertion) | NA | 1.762 | 1.522 |
| cob(II)yrinate a,c-diamide biosynthesis II (late cobalt incorporation) | -0.532 | 1.015 | NA |
| coenzyme B biosynthesis | -3.071 | 6.334 | 3.263 |
| coenzyme M biosynthesis I | -0.653 | NA | NA |
| creatinine degradation I | -0.731 | 1.411 | 0.68 |
| creatinine degradation II | -1.043 | 1.966 | 0.923 |
| D-fructuronate degradation | -0.69 | 0.755 | NA |
| D-galactarate degradation I | -1.061 | 0.528 | -0.533 |
| D-galacturonate degradation I | NA | 0.635 | NA |
| D-glucarate degradation I | -1.669 | NA | -1.318 |
| dTDP-N-acetylthomosamine biosynthesis | -1.002 | 0.717 | NA |
| ectoine biosynthesis | -0.56 | 0.587 | NA |
| enterobacterial common antigen biosynthesis | -6.585 | 6.736 | NA |
| enterobactin biosynthesis | -2.12 | 1.54 | -0.58 |
| ergothioneine biosynthesis I (bacteria) | -5.705 | 4.133 | -1.572 |
| ethylmalonyl-CoA pathway | NA | 1.609 | 1.126 |
| factor 420 biosynthesis | -3.584 | 7.65 | 4.066 |
| formaldehyde assimilation I (serine pathway) | -1.108 | NA | -1.106 |
| formaldehyde assimilation II (RuMP Cycle) | -0.539 | 1.394 | 0.855 |
| formaldehyde oxidation I | -0.531 | 1.373 | 0.843 |
| galactose degradation I (Leloir pathway) | 0.553 | NA | NA |
| gallate degradation I | -1.414 | 2.933 | 1.519 |
| gallate degradation II | -1.449 | 2.968 | 1.52 |
| GDP-D-glycero-&alpha;-D-manno-heptose biosynthesis | 1.444 | -2.19 | -0.746 |
| glucose and glucose-1-phosphate degradation | -0.788 | 0.798 | NA |
| glucose degradation (oxidative) | -2.795 | 1.294 | -1.501 |
| glutaryl-CoA degradation | 0.7 | -1.314 | -0.614 |
| glycerol degradation to butanol | -1.06 | 1.965 | 0.905 |
| glycine betaine degradation I | NA | 1.479 | 1.012 |
| glycogen degradation I (bacterial) | 0.541 | -0.531 | NA |
| glycogen degradation II (eukaryotic) | -0.826 | 1.763 | 0.937 |
| glyoxylate cycle | -0.581 | 0.505 | NA |
| heterolactic fermentation | -1.206 | 1.333 | NA |
| hexitol fermentation to lactate, formate, ethanol and acetate | -3.151 | 3.158 | NA |
| incomplete reductive TCA cycle | 0.575 | -0.655 | NA |
| isoprene biosynthesis II (engineered) | 1.375 | -1.53 | NA |
| isopropanol biosynthesis | NA | -0.788 | -1.081 |
| ketogluconate metabolism | -2.08 | 2.904 | 0.823 |
| L-1,2-propanediol degradation | -3.483 | 7.5 | 4.017 |
| L-arabinose degradation IV | -1.091 | 10.119 | 9.027 |
| L-arginine degradation II (AST pathway) | -2.36 | 2.826 | NA |
| L-glutamate degradation V (via hydroxyglutarate) | 0.929 | -1.452 | -0.522 |
| L-histidine degradation II | -0.905 | 1.758 | 0.853 |
| L-isoleucine biosynthesis IV | 0.648 | -0.637 | NA |
| L-lysine biosynthesis II | -3.403 | 5.364 | 1.961 |
| L-lysine fermentation to acetate and butanoate | 0.561 | 1.344 | 1.905 |
| L-methionine biosynthesis I | NA | 0.651 | NA |
| L-methionine salvage cycle III | -6.548 | 9.365 | 2.817 |
| L-rhamnose degradation I | -0.601 | 0.575 | NA |
| L-tryptophan degradation IX | -0.755 | 0.943 | NA |
| L-tryptophan degradation to 2-amino-3-carboxymuconate semialdehyde | -1.013 | 0.945 | NA |
| L-tryptophan degradation XII (Geobacillus) | -1.826 | 1.525 | NA |
| L-tyrosine degradation I | -0.69 | 0.791 | NA |
| L-valine degradation I | -2.825 | 6.469 | 3.644 |
| lactose and galactose degradation I | -5.366 | 8.296 | 2.93 |
| mannan degradation | NA | 0.979 | 0.8 |
| meta cleavage pathway of aromatic compounds | -2.222 | 2.442 | NA |
| methanogenesis from acetate | 1.383 | -1.405 | NA |
| methanol oxidation to carbon dioxide | -1.267 | 1.652 | NA |
| methyl ketone biosynthesis | -0.595 | NA | -0.695 |
| methylaspartate cycle | NA | NA | 0.622 |
| methylgallate degradation | -1.426 | 2.944 | 1.518 |
| methylphosphonate degradation I | NA | 1.155 | 0.657 |
| mevalonate pathway I | 0.62 | -0.789 | NA |
| mevalonate pathway II (archaea) | 2.767 | -2.41 | NA |
| mono-trans, poly-cis decaprenyl phosphate biosynthesis | -4.413 | 7.054 | 2.641 |
| mycothiol biosynthesis | -1.242 | 0.619 | -0.623 |
| myo-, chiro- and scillo-inositol degradation | -1.407 | 2.468 | 1.061 |
| myo-inositol degradation I | -1.294 | 2.424 | 1.13 |
| NAD biosynthesis II (from tryptophan) | -0.799 | 0.725 | NA |
| NAD salvage pathway II | -3.019 | 2.894 | NA |
| nicotinate degradation I | -5.063 | 8.194 | 3.131 |
| nitrate reduction VI (assimilatory) | -1.019 | 1.286 | NA |
| nitrifier denitrification | -2.899 | 1.945 | -0.954 |
| norspermidine biosynthesis | -0.953 | 1.536 | 0.583 |
| nylon-6 oligomer degradation | -0.85 | 0.947 | NA |
| octane oxidation | -0.822 | 0.989 | NA |
| palmitate biosynthesis II (bacteria and plants) | NA | -0.905 | -1.09 |
| peptidoglycan biosynthesis II (staphylococci) | -5.146 | 11.401 | 6.254 |
| peptidoglycan biosynthesis IV (Enterococcus faecium) | -3.081 | 3.053 | NA |
| peptidoglycan biosynthesis V (&beta;-lactam resistance) | -3.874 | 6.005 | 2.131 |
| phenylacetate degradation I (aerobic) | -2.362 | 2.021 | NA |
| phospholipases | -1.595 | 1.442 | NA |
| polymyxin resistance | -3.01 | 1.664 | -1.346 |
| ppGpp biosynthesis | -0.775 | 1.179 | NA |
| protocatechuate degradation I (meta-cleavage pathway) | -1.35 | 3.374 | 2.025 |
| protocatechuate degradation II (ortho-cleavage pathway) | -0.963 | 1.364 | NA |
| purine nucleotides degradation II (aerobic) | NA | 0.761 | 0.626 |
| purine ribonucleosides degradation | NA | 0.974 | 1.037 |
| pyrimidine deoxyribonucleotides biosynthesis from CTP | 1.42 | -2.896 | -1.476 |
| pyrimidine deoxyribonucleotides de novo biosynthesis III | 0.531 | NA | NA |
| pyrimidine deoxyribonucleotides de novo biosynthesis IV | 1.396 | -2.894 | -1.498 |
| pyruvate fermentation to acetate and lactate II | 0.607 | NA | NA |
| pyruvate fermentation to acetone | -1.116 | NA | -0.677 |
| pyruvate fermentation to butanoate | 1.011 | -0.849 | NA |
| reductive acetyl coenzyme A pathway | 1.077 | -1.026 | NA |
| S-adenosyl-L-methionine cycle I | NA | 1.435 | 1.331 |
| S-methyl-5-thio-&alpha;-D-ribose 1-phosphate degradation | -6.882 | 9.53 | 2.648 |
| spirilloxanthin and 2,2'-diketo-spirilloxanthin biosynthesis | -0.97 | 2.188 | 1.218 |
| sucrose degradation II (sucrose synthase) | 0.736 | -2.09 | -1.354 |
| sucrose degradation III (sucrose invertase) | -1.619 | 2.032 | NA |
| superpathway of (Kdo)2-lipid A biosynthesis | -0.563 | -0.78 | -1.344 |
| superpathway of (R,R)-butanediol biosynthesis | -0.822 | NA | -1.035 |
| superpathway of &beta;-D-glucuronide and D-glucuronate degradation | -0.856 | 0.788 | NA |
| superpathway of 2,3-butanediol biosynthesis | -0.542 | NA | -0.89 |
| superpathway of aerobic toluene degradation | -1.618 | 1.714 | NA |
| superpathway of bacteriochlorophyll a biosynthesis | -0.717 | 1.655 | 0.938 |
| superpathway of C1 compounds oxidation to CO2 | NA | 3.717 | 3.424 |
| superpathway of chorismate metabolism | -1.082 | NA | -0.77 |
| superpathway of Clostridium acetobutylicum acidogenic fermentation | 0.964 | -0.791 | NA |
| superpathway of D-glucarate and D-galactarate degradation | -1.061 | 0.528 | -0.533 |
| superpathway of demethylmenaquinol-6 biosynthesis I | -0.559 | NA | NA |
| superpathway of demethylmenaquinol-6 biosynthesis II | 1.413 | 0.828 | 2.241 |
| superpathway of demethylmenaquinol-8 biosynthesis | -0.555 | NA | NA |
| superpathway of demethylmenaquinol-9 biosynthesis | -0.559 | NA | NA |
| superpathway of fucose and rhamnose degradation | -1.558 | 3.619 | 2.061 |
| superpathway of geranylgeranyldiphosphate biosynthesis I (via mevalonate) | 0.629 | -0.792 | NA |
| superpathway of glycerol degradation to 1,3-propanediol | -0.507 | 3.473 | 2.966 |
| superpathway of glycol metabolism and degradation | -1.911 | 4.578 | 2.667 |
| superpathway of glyoxylate bypass and TCA | -0.568 | NA | NA |
| superpathway of hexitol degradation (bacteria) | -1.444 | 1.087 | NA |
| superpathway of hexuronide and hexuronate degradation | -0.937 | 1.092 | NA |
| superpathway of L-arginine and L-ornithine degradation | -5.65 | 7.165 | 1.516 |
| superpathway of L-arginine, putrescine, and 4-aminobutanoate degradation | -5.65 | 7.165 | 1.516 |
| superpathway of L-aspartate and L-asparagine biosynthesis | 0.576 | NA | 0.74 |
| superpathway of L-threonine metabolism | -6.335 | 9.146 | 2.811 |
| superpathway of menaquinol-10 biosynthesis | -0.508 | NA | NA |
| superpathway of menaquinol-11 biosynthesis | -0.52 | NA | NA |
| superpathway of menaquinol-12 biosynthesis | -0.52 | NA | NA |
| superpathway of menaquinol-13 biosynthesis | -0.52 | NA | NA |
| superpathway of menaquinol-6 biosynthesis I | -0.508 | NA | NA |
| superpathway of menaquinol-8 biosynthesis II | 1.413 | -0.787 | 0.626 |
| superpathway of menaquinol-9 biosynthesis | -0.508 | NA | NA |
| superpathway of methylglyoxal degradation | -2.183 | 3.746 | 1.563 |
| superpathway of N-acetylglucosamine, N-acetylmannosamine and N-acetylneuraminate degradation | -0.896 | NA | -0.715 |
| superpathway of N-acetylneuraminate degradation | -0.663 | NA | -0.566 |
| superpathway of phenylethylamine degradation | -2.206 | 4.726 | 2.52 |
| superpathway of phylloquinol biosynthesis | -0.712 | NA | NA |
| superpathway of purine deoxyribonucleosides degradation | NA | 0.937 | 0.905 |
| superpathway of pyridoxal 5'-phosphate biosynthesis and salvage | NA | 2.341 | 2.37 |
| superpathway of pyrimidine deoxyribonucleosides degradation | NA | 0.648 | 0.768 |
| superpathway of salicylate degradation | -2.333 | 2.405 | NA |
| superpathway of sulfolactate degradation | NA | 1.819 | 1.324 |
| superpathway of sulfur oxidation (Acidianus ambivalens) | 1.533 | -1.735 | NA |
| superpathway of thiamin diphosphate biosynthesis II | 1.049 | -1.069 | NA |
| superpathway of UDP-glucose-derived O-antigen building blocks biosynthesis | -0.577 | 1.115 | 0.537 |
| superpathway of vanillin and vanillate degradation | -1.328 | 3.437 | 2.11 |
| TCA cycle VII (acetate-producers) | NA | 0.716 | NA |
| teichoic acid (poly-glycerol) biosynthesis | -1.136 | 4.14 | 3.004 |
| thiazole biosynthesis II (Bacillus) | 1.319 | -1.274 | NA |
| toluene degradation III (aerobic) (via p-cresol) | -2.388 | 2.255 | NA |
| toluene degradation IV (aerobic) (via catechol) | -2.366 | 2.877 | 0.511 |
| tRNA processing | NA | -0.591 | NA |
| UDP-2,3-diacetamido-2,3-dideoxy-&alpha;-D-mannuronate biosynthesis | 0.615 | -0.585 | NA |
| vanillin and vanillate degradation I | -1.328 | 3.437 | 2.11 |
| vanillin and vanillate degradation II | -1.325 | 3.426 | 2.101 |
| vitamin E biosynthesis (tocopherols) | -2.895 | 7.085 | 4.19 |
