## Appendix A for "More than a stick in the mud: Eelgrass leaf and root bacterial communities are distinct from those on physical mimics"

We also had controls that we transplanted back to sites with mimics (in addition to control plants taken from nearby where mimics were placed). Appendix A repeats the analyses in the main text and produces very similar results in terms of degrees of differences compared to the transplants in the main text. This suggests that these results are robust to our manipulations.

*Sampling and sequencing success*

We identified 43,118 bacterial ASVs across 247 leaf, root, mimic, and sediment samples after quality filtering samples to 5,165,712 reads. Root samples contained between 328 and 1,026 bacterial ASVs on their surface (we measured 75 root samples with read depth between 7,501 and 79,940 reads), Root mimics samples contained between 85 and 797 bacterial ASVs on their surface (we measured 26 root mimic samples with read depth between 2,056 and 37,853 reads), Sediment samples contained between 271 and 843 bacterial ASVs on their surface (we measured 36 sediment root samples with read depth between 10,775 and 40,346 reads), leaf samples contained between 261 and 716 bacterial ASVs on their surface (we measured 75 leaf samples with read depth between 5,770 and 50,564 reads) and leaf mimic samples which contained between 196 and 724 bacterial ASVs (35 leaf mimic samples with between 4,591 and 31,702 reads per sample).

**Results**

Leaf mimics have a higher number of ASVs than leaves (negative binomial glm, ANOVA, p = 0.048; Appendix Figure 1a). When examining core ASVs (present in at least 50% of samples of a type at at least 1% detection rate), we found that leaves and leaf mimics largely harbored distinct bacterial communities, by ASV, while also sharing substantial overlap (Appendix Figure 1b) and more so than other sample types. Of the 212 ASVS in the core leaf microbiome, 107 or 50% were found only on leaves and 97 of the remaining (56% of core leaf ASVs) overlapped with leaf mimic ASVs. While there was not a difference in variance within each sample type (betadisper ANOVA, p = 0.71), the composition of the two groups was different (PERMANOVA, F = 19.61, p = 0.001, r2 = 0.15, Appendix Figure 2a). When we examine overlap in predicted Metacyc pathways, we found that there was a significant difference between leaves and mimics (PERMANOVA, r2 = 0.05411, F = 6.1787, p = 0.001), though this effect was weaker than for the sequence based compositional differences (Appendix Figure 2b).

Through analysis of specific ASVs that varied between leaves and leaf mimics via DESEQ2, we found 209 ASV that showed higher relative abundance on leaves and 101 that showed higher relative abundance on mimics. Only four families contained more than ten ASVs that varied between mimics and leaves: Flavobacteriaceae (15 higher on leaves, 17 higher on mimics), Pirellulaceae (eight higher on leaves, four higher on mimics), Rhodobacteraceae (40 higher on leaves, 25 higher on mimics), and Saprospiraceae (39 higher on leaves, one higher on mimics);  most families contained fewer than 3 ASVs that varied between leaves and mimics  (Supplementary Table 1). Within these families several genera were represented by multiple ASVs. These included *Kordia* (three ASVs higher on leaves), *Maribacter* (one higher on leaves, two higher on mimics) *Ulvibacter* (two higher on leaves, two higher on mimics), *Winogradskyella* (one higher on leaves, two higher on mimics), *Blastopirellula* (5 higher on leaves, 2 higher on mimics), *Rhodopirellula* (two higher on leaves), *Octadecabacter* (one higher on leaves, one higher on mimics), *Roseobacter* (one higher on leaves, two higher on mimics), *Sedimentitalea* (one higher on leaves, one on mimics), *Sulfitobacter* (three higher on leaves, three higher on mimics), *Tateyamaria* (one higher on leaves, one on mimics), *Yoonia-Loktanella* (four higher on leaves), *Lewinella* (five higher on leaves), *Portibacter* (four higher on mimics), and *Rubidimonas* (three higher on leaves). Six other genera (not in these families) contained multiple ASVs that varied between leaves and mimics (Supplementary Table 2).

When we examined pathways that changed between the leaf and leaf mimic microbiomes, we identified 82 pathways that changed, 16 upregulated in leaf microbiomes and 66 upregulated on mimic microbiomes (Supplemental Table 2). These pathways were generally unremarkable (likely at least in part due to limits in prediction of environmental microbial pathways), though did indicate that aerobic environments might not solely limited to leaf microbiomes with an upregulation of the superpathway of sulfur oxidation on mimic leaf surfaces compared to leaf surfaces.

In roots, less surprisingly, we also found differences between the mimic and root communities. We found that despite fewer ASVs found summed across samples in mimics (Appendix Figure 1b), there were generally the same number of ASVs on roots and root mimics (Appendix Figure 1a, negative binomial glm ANOVA, p = 0.093, p = 0.32 when comparing sediments as well). We found that there was no difference in variance among roots and root mimics (betadisper ANOVA p = 0.87), however sediments showed less variance than either of the other two groups (betadisper ANOVA, p < 0.001, Tukey’s HSD sediment vs mimic p < 0.001, vs roots p < 0.001). When examining core ASVs (present in at least 50% of samples of a given type at at least 1% detection rate), we found that roots and sediments largely harbored distinct bacterial communities, by ASV (Appendix Figure 1b), though root mimics had fewer unique core microbiome (only 2 ASVs unique to root mimics). Of 266 ASVS in the core root microbiome, 82 or 31% were found only on roots and 131 (49%) were found only on roots and in sediments. Only 38 core root ASVs (14%) were shared between roots and root mimics.  When we examined all ASVs (without core restrictions), root mimics had more taxa unique to their sample type, indicating considerable  variability in communities assembled on root mimics (Supp Appendix Figure 1). ASV composition on roots, mimics and sediment were compositionally distinct (PERMANOVA r2 = 0.151 F = 17.672 p =  0.001, see Supplemental Table 3 for pairwise comparisons). We found many ASVs varied in abundance between these groups (417 between sediments and mimics, 454 between roots and sediments, and 385 between roots and mimics). Of these, the majority were at higher abundances on roots or sediments compared to mimics (comparing roots to mimics, 437 were higher on roots, and 49 were higher on mimics; comparing sediment to mimics, 359 were higher in sediments, 98 were higher on mimics; comparing roots to sediments 265 were higher on roots, 240 were higher in sediments; see Supplemental Table 5 for more details). The families that had the largest number of taxa vary among sample types included Spirochaetaceae (32 ASVs), Thiotrichaceae (33 ASVs), Bactoroidetes BD2-2 (47 ASVs), Pirellulaceae (56 ASVs), Desulfosarcinaceae (63 ASVs), Desusulfocapsaceae (69 ASVs), Saprospiraceae (73 ASVs), Rhodobacteraceae (125 ASVs) and Flavobacteriaceae (130 ASVs). Number of ASVs at higher or lower relative abundances in these different treatments can be found in Appendix Table 7.While the pathways that varied were numerous and not particularly remarkable (as indicated in Supplemental Table 6), we found that indicated pathways were generally indicated to be upregulated on mimics in pairwise comparisons (141 pathways higher in mimics compared to 32 in sediments, and 69 higher on mimics compared to 11 on roots).

Appendix Figures and Tables:

Appendix Figure 1: (A) Mean amplicon sequence variant (ASV) richness found in each type of sample we measured. Raw data as well as means and standard errors are presented. Leaf mimic bacterial communities had a higher mean richness than leaf bacterial communities (p < 0.001); root mimic, root, and sediment bacterial communities did not differ in mean community richness. (B) Overlap among core bacterial communities showing shared ASVs present in each sample type in at least 50% of samples at a 1% detection rate. Diagram is a barplot of shared community memberships, equivalent to a Venn diagram.

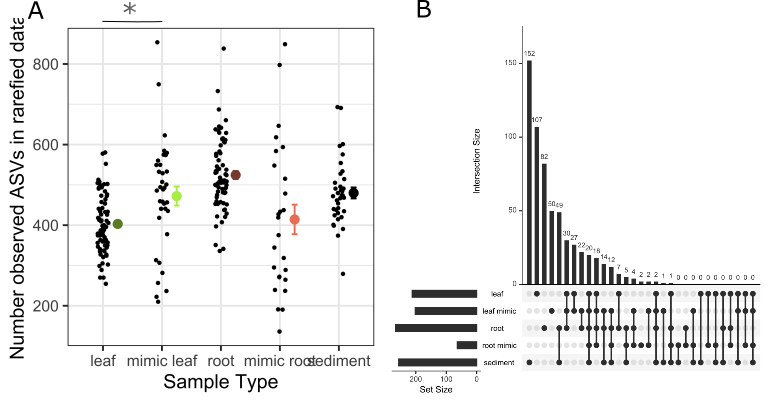

Appendix Figure 2: (A) Ordination of bacterial community structure based on principal coordinate analysis of phylogenetic-isometric log-ratio transformed distances. (B) Ordination of predicted Metacyc pathways structure based on principal coordinate analysis of centered log-ratio transformed distances. Bright green points are communities on leaf mimics and dark green points are communities on leaves. Leaf and leaf mimic communities in both analyses are distinct from each other (PERMANOVA p < 0.001).

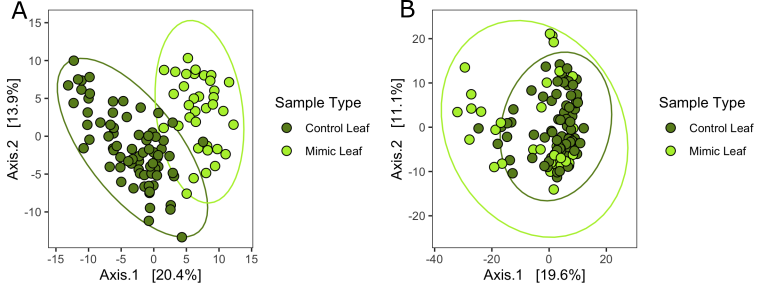

Appendix Figure 3: (A) Ordination of bacterial community structure based on principal coordinate analysis of phylogenetic-isometric log-ratio transformed distances. (B) Ordination of predicted Metacyc pathways structure based on principal coordinate analysis of centered log-ratio transformed distances. Red-orange points are communities on root mimics, dark brown points are communities on roots, and grey points are communities in sediments. All communities are distinct from each other in each analysis (PERMANOVA p < 0.001).

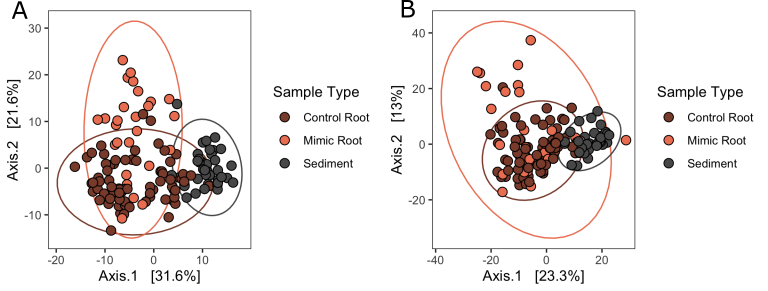

Appendix Table 1: For both leaves and root communities, the five families that had the most ASVs vary between mimics and seagrass substrate. See supplemental Tables 1 and 6 for complete lists for leaves and roots respectively.

| **Family** | **Higher on leaves** | **Higher on mimics** | |  |  |  |
| --- | --- | --- | --- | --- | --- | --- |
| Rhodobacteraceae | 40 | 25 |  |  |  |  |
| Saprospiraceae | 39 | 1 |  |  |  |  |
| Flavobacteriaceae | 15 | 17 |  |  |  |  |
| Pirellulaceae | 8 | 4 |  |  |  |  |
| Rhizobiaceae | 1 | 8 |  |  |  |  |
| **Family** | **Higher on roots** | **Higher on mimics** | **Higher on mimics** | **Higher on sediment** | **Higher on roots** | **Higher on sediment** |
| Flavobacteriaceae | 38 | 3 | 32 | 12 | 37 | 8 |
| Rhodobacteraceae | 36 | 3 | 41 | 0 | 45 | 0 |
| Desulfocapsaceae | 24 | 0 | 6 | 15 | 14 | 10 |
| Saprospiraceae | 19 | 3 | 21 | 4 | 23 | 3 |
| Desulfosarcinaceae | 14 | 3 | 0 | 23 | 0 | 23 |

**Supplemental Tables legends (followed by supplemental tables themselves)**

Supplemental Table 1: For leaf bacterial communities, the family-level identification of ASVs that varied significantly between mimics and seagrass substrate determined by DESeq2.

Supplemental Table 2:  For leaf bacterial communities, the genus-level identification of ASVs that varied significantly between mimics and seagrass substrate determined by DESeq2.

Supplemental Table 3: For leaf bacterial communities, all ASVs that varied significantly between mimics and seagrass substrate determined by DESeq2, including magnitude of differences.

Supplemental Table 4:  For leaf bacterial communities, all Metacyc predicted pathways that varied significantly between mimics and seagrass substrate determined by DESeq2, including magnitude of differences.

Supplemental Table 5: Results of pairwise PERMANOVA tests distinguishing compositional differences among roots, root mimics, and sediments in both ASV composition and composition of predicted Metacyc pathways.

Supplemental Table 6: For belowground bacterial communities, the family-level identification of ASVs that varied significantly among mimics, seagrass and sediment determined by DESeq2.

Supplemental Table 7: For belowground bacterial communities, all ASVs that varied significantly among mimics, seagrass and sediment determined by DESeq2, including magnitude of differences.

Supplemental Table 8:  For belowground bacterial communities, all Metacyc predicted pathways that varied significantly among mimics, seagrass and sediment determined by DESeq2, including magnitude of differences.

Appendix Supp Figure 1

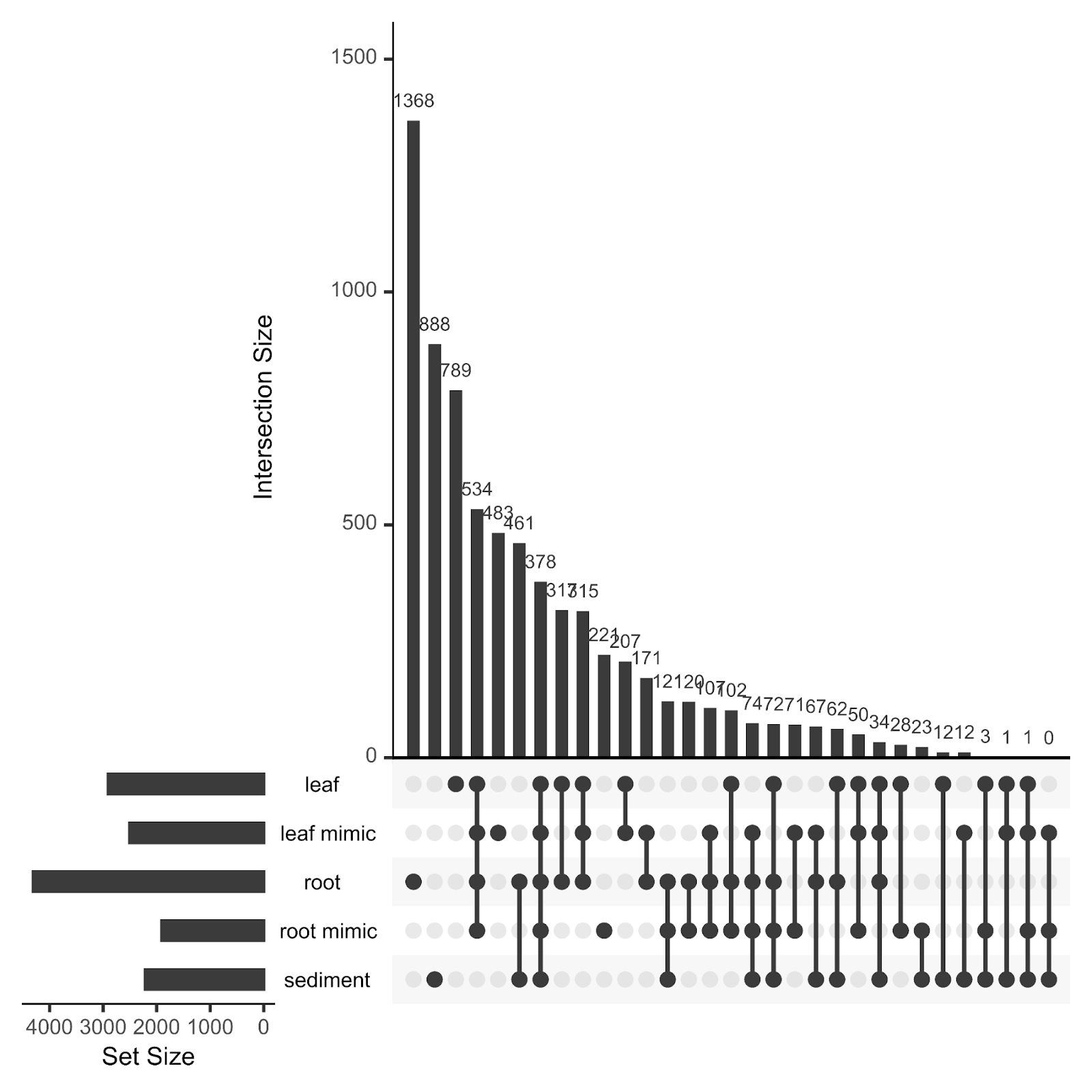

Appendix Supplemental Table 1

| Family | Higher on leaves | Higher on mimics |
| --- | --- | --- |
| 37-13 | 1 | 0 |
| A4b | 1 | 0 |
| Alteromonadaceae | 8 | 0 |
| Arenicellaceae | 1 | 0 |
| Bdellovibrionaceae | 2 | 0 |
| Bernardetiaceae | 1 | 0 |
| Blastocatellaceae | 0 | 1 |
| Caldilineaceae | 2 | 0 |
| Cellvibrionaceae | 3 | 0 |
| Chromatiaceae | 2 | 1 |
| Colwelliaceae | 4 | 0 |
| Crocinitomicaceae | 8 | 0 |
| Cryomorphaceae | 8 | 0 |
| Cyanobiaceae | 0 | 1 |
| Cyclobacteriaceae | 2 | 0 |
| Desulfobulbaceae | 0 | 1 |
| Desulfocapsaceae | 1 | 5 |
| DEV007 | 3 | 3 |
| Flammeovirgaceae | 1 | 0 |
| Flavobacteriaceae | 15 | 17 |
| Fokiniaceae | 1 | 0 |
| Francisellaceae | 1 | 0 |
| Gimesiaceae | 0 | 1 |
| Granulosicoccaceae | 5 | 2 |
| Halieaceae | 3 | 1 |
| Halomonadaceae | 0 | 1 |
| Hyphomicrobiaceae | 0 | 1 |
| Hyphomonadaceae | 3 | 2 |
| Ilumatobacteraceae | 0 | 1 |
| Kangiellaceae | 1 | 0 |
| KD3-93 | 1 | 0 |
| Legionellaceae | 0 | 1 |
| Magnetospiraceae | 1 | 0 |
| Marinomonadaceae | 1 | 0 |
| Methylophagaceae | 2 | 0 |
| Methylophilaceae | 5 | 1 |
| Micavibrionaceae | 0 | 2 |
| Microtrichaceae | 2 | 2 |
| Nitrincolaceae | 2 | 0 |
| Nitrosococcaceae | 1 | 0 |
| NS11-12_marine_group | 1 | 0 |
| NS9_marine_group | 1 | 0 |
| Oleiphilaceae | 1 | 0 |
| Opitutaceae | 1 | 0 |
| Phormidesmiaceae | 0 | 1 |
| Phycisphaeraceae | 3 | 0 |
| Pirellulaceae | 8 | 4 |
| Porticoccaceae | 1 | 0 |
| Prolixibacteraceae | 2 | 0 |
| Pseudohongiellaceae | 2 | 0 |
| Rhizobiaceae | 1 | 8 |
| Rhizobiales_Incertae_Sedis | 0 | 1 |
| Rhodobacteraceae | 40 | 25 |
| Rhodothermaceae | 1 | 0 |
| Rickettsiaceae | 2 | 0 |
| Rubinisphaeraceae | 1 | 3 |
| Rubritaleaceae | 1 | 2 |
| Saprospiraceae | 39 | 1 |
| Schleiferiaceae | 1 | 0 |
| Shewanellaceae | 0 | 1 |
| Sphingomonadaceae | 3 | 3 |
| Spirosomaceae | 1 | 0 |
| Spongiibacteraceae | 2 | 0 |
| Sulfurimonadaceae | 1 | 0 |
| Sulfurovaceae | 0 | 3 |
| Tenderiaceae | 1 | 0 |
| Terasakiellaceae | 1 | 0 |
| Thioglobaceae | 1 | 0 |
| Thiotrichaceae | 1 | 1 |
| Unknown_Family | 0 | 2 |
| Woeseiaceae | 0 | 2 |

Appendix Supplemental Table 2

| **Family** | **Genus** | **Higher on leaves** | **Higher on mimics** |
| --- | --- | --- | --- |
| Alteromonadaceae | Glaciecola | 6 | 0 |
| Alteromonadaceae | Paraglaciecola | 1 | 0 |
| Alteromonadaceae | Salinimonas | 1 | 0 |
| Arenicellaceae | Arenicella | 1 | 0 |
| Bdellovibrionaceae | Bdellovibrio | 1 | 0 |
| Bdellovibrionaceae | OM27_clade | 1 | 0 |
| Bernardetiaceae | Garritya | 1 | 0 |
| Blastocatellaceae | Blastocatella | 0 | 1 |
| Cellvibrionaceae | Agaribacterium | 1 | 0 |
| Cellvibrionaceae | Candidatus_Endobugula | 1 | 0 |
| Chromatiaceae | Candidatus_Thiobios | 2 | 0 |
| Chromatiaceae | Halochromatium | 0 | 1 |
| Colwelliaceae | Colwellia | 3 | 0 |
| Colwelliaceae | Thalassotalea | 1 | 0 |
| Crocinitomicaceae | Crocinitomix | 4 | 0 |
| Crocinitomicaceae | Fluviicola | 3 | 0 |
| Cryomorphaceae | Vicingus | 3 | 0 |
| Cyanobiaceae | Synechococcus_CC9902 | 0 | 1 |
| Cyclobacteriaceae | Ekhidna | 1 | 0 |
| Cyclobacteriaceae | Fabibacter | 1 | 0 |
| Desulfobulbaceae | Desulfobulbus | 0 | 1 |
| Flavobacteriaceae | Actibacter | 0 | 1 |
| Flavobacteriaceae | Aquibacter | 0 | 2 |
| Flavobacteriaceae | Aurantivirga | 2 | 0 |
| Flavobacteriaceae | Changchengzhania | 1 | 0 |
| Flavobacteriaceae | Formosa | 1 | 0 |
| Flavobacteriaceae | Jejudonia | 0 | 1 |
| Flavobacteriaceae | Kordia | 3 | 0 |
| Flavobacteriaceae | Lutibacter | 0 | 1 |
| Flavobacteriaceae | Maribacter | 1 | 2 |
| Flavobacteriaceae | NS2b_marine_group | 1 | 0 |
| Flavobacteriaceae | NS3a_marine_group | 1 | 0 |
| Flavobacteriaceae | Polaribacter | 1 | 0 |
| Flavobacteriaceae | Psychroserpens | 0 | 1 |
| Flavobacteriaceae | Robiginitalea | 0 | 1 |
| Flavobacteriaceae | Ulvibacter | 2 | 2 |
| Flavobacteriaceae | Wenyingzhuangia | 1 | 0 |
| Flavobacteriaceae | Winogradskyella | 1 | 2 |
| Fokiniaceae | MD3-55 | 1 | 0 |
| Granulosicoccaceae | Granulosicoccus | 5 | 2 |
| Halieaceae | Halioglobus | 0 | 1 |
| Halieaceae | OM60(NOR5)_clade | 2 | 0 |
| Halieaceae | Pseudohaliea | 1 | 0 |
| Halomonadaceae | Halomonas | 0 | 1 |
| Hyphomicrobiaceae | Filomicrobium | 0 | 1 |
| Hyphomonadaceae | Hellea | 1 | 0 |
| Hyphomonadaceae | Hyphomonas | 0 | 1 |
| Hyphomonadaceae | Litorimonas | 1 | 0 |
| Hyphomonadaceae | Robiginitomaculum | 1 | 0 |
| Ilumatobacteraceae | Ilumatobacter | 0 | 1 |
| Magnetospiraceae | Magnetospira | 1 | 0 |
| Marinomonadaceae | Marinomonas | 1 | 0 |
| Methylophagaceae | Marine_Methylotrophic_Group_3 | 1 | 0 |
| Methylophilaceae | Methylotenera | 5 | 1 |
| Microtrichaceae | Sva0996_marine_group | 1 | 2 |
| Nitrosococcaceae | Cm1-21 | 1 | 0 |
| Oleiphilaceae | Oleiphilus | 1 | 0 |
| Opitutaceae | Diplosphaera | 1 | 0 |
| Phormidesmiaceae | Phormidesmis_ANT.LACV5.1 | 0 | 1 |
| Phycisphaeraceae | Phycisphaera | 1 | 0 |
| Phycisphaeraceae | SM1A02 | 2 | 0 |
| Pirellulaceae | Blastopirellula | 5 | 2 |
| Pirellulaceae | Pir4_lineage | 0 | 1 |
| Pirellulaceae | Pirellula | 0 | 1 |
| Pirellulaceae | Rhodopirellula | 2 | 0 |
| Pirellulaceae | Rubripirellula | 1 | 0 |
| Porticoccaceae | C1-B045 | 1 | 0 |
| Prolixibacteraceae | Draconibacterium | 2 | 0 |
| Pseudohongiellaceae | Pseudohongiella | 2 | 0 |
| Rhizobiaceae | Ahrensia | 0 | 1 |
| Rhizobiaceae | Hoeflea | 0 | 1 |
| Rhizobiaceae | Pseudahrensia | 1 | 4 |
| Rhizobiales_Incertae_Sedis | Anderseniella | 0 | 1 |
| Rhodobacteraceae | Aliiroseovarius | 1 | 0 |
| Rhodobacteraceae | Celeribacter | 0 | 1 |
| Rhodobacteraceae | Jannaschia | 0 | 1 |
| Rhodobacteraceae | Leisingera | 0 | 1 |
| Rhodobacteraceae | Limibaculum | 1 | 0 |
| Rhodobacteraceae | Octadecabacter | 1 | 1 |
| Rhodobacteraceae | Phaeobacter | 0 | 1 |
| Rhodobacteraceae | Planktomarina | 0 | 1 |
| Rhodobacteraceae | Planktotalea | 1 | 0 |
| Rhodobacteraceae | Roseobacter | 1 | 2 |
| Rhodobacteraceae | Roseovarius | 0 | 1 |
| Rhodobacteraceae | Sedimentitalea | 1 | 1 |
| Rhodobacteraceae | Sulfitobacter | 3 | 3 |
| Rhodobacteraceae | Tateyamaria | 1 | 1 |
| Rhodobacteraceae | Thiobacimonas | 0 | 1 |
| Rhodobacteraceae | Tropicimonas | 0 | 1 |
| Rhodobacteraceae | Yoonia-Loktanella | 4 | 0 |
| Rickettsiaceae | Candidatus_Megaira | 2 | 0 |
| Rubinisphaeraceae | Fuerstia | 1 | 0 |
| Rubinisphaeraceae | Planctomicrobium | 0 | 2 |
| Rubritaleaceae | Haloferula | 0 | 1 |
| Rubritaleaceae | Persicirhabdus | 0 | 1 |
| Rubritaleaceae | Roseibacillus | 1 | 0 |
| Saprospiraceae | Aureispira | 1 | 0 |
| Saprospiraceae | Lewinella | 5 | 0 |
| Saprospiraceae | Phaeodactylibacter | 1 | 0 |
| Saprospiraceae | Portibacter | 4 | 0 |
| Saprospiraceae | Rubidimonas | 3 | 0 |
| Schleiferiaceae | Schleiferia | 1 | 0 |
| Shewanellaceae | Shewanella | 0 | 1 |
| Sphingomonadaceae | Altererythrobacter | 0 | 1 |
| Sphingomonadaceae | Erythrobacter | 2 | 0 |
| Sphingomonadaceae | Parasphingopyxis | 0 | 1 |
| Sphingomonadaceae | Sphingorhabdus | 1 | 0 |
| Spirosomaceae | Taeseokella | 1 | 0 |
| Spongiibacteraceae | Dasania | 1 | 0 |
| Sulfurimonadaceae | Sulfurimonas | 1 | 0 |
| Sulfurovaceae | Sulfurovum | 0 | 3 |
| Tenderiaceae | Candidatus_Tenderia | 1 | 0 |
| Thioglobaceae | SUP05_cluster | 1 | 0 |
| Thiotrichaceae | Cocleimonas | 0 | 1 |
| Thiotrichaceae | Leucothrix | 1 | 0 |
| Unknown_Family | Marinicella | 0 | 1 |
| Woeseiaceae | Woeseia | 0 | 2 |

Appendix Supplemental Table 3

| **Family** | **Genus** | **Species** | **baseMean** | **log2FoldChange** | **lfcSE** | **stat** | **pvalue** | **padj** |
| --- | --- | --- | --- | --- | --- | --- | --- | --- |
| Blastocatellaceae | Blastocatella | NA | 6.434 | -3.014 | 0.859 | 9.377 | 0.002 | 0.014 |
| Pirellulaceae | Rubripirellula | NA | 13.893 | 7.241 | 1.285 | 8.033 | 0.005 | 0.025 |
| Pirellulaceae | Rhodopirellula | NA | 16.519 | 3.544 | 0.647 | 19.603 | 0 | 0 |
| Pirellulaceae | Blastopirellula | NA | 39.708 | 2.587 | 0.311 | 48.476 | 0 | 0 |
| Pirellulaceae | Pirellula | NA | 4.711 | -2.877 | 0.768 | 9.371 | 0.002 | 0.014 |
| Pirellulaceae | Rhodopirellula | NA | 14.974 | 2.126 | 0.423 | 19.064 | 0 | 0 |
| Pirellulaceae | Blastopirellula | NA | 5.49 | -3.413 | 0.714 | 17.136 | 0 | 0 |
| Pirellulaceae | Blastopirellula | NA | 28.816 | 1.94 | 0.481 | 12.162 | 0 | 0.004 |
| Pirellulaceae | Pir4_lineage | NA | 1.763 | -3.891 | 1.577 | 7.325 | 0.007 | 0.034 |
| Phycisphaeraceae | Phycisphaera | NA | 13.048 | 2.782 | 0.599 | 13.549 | 0 | 0.002 |
| Phycisphaeraceae | SM1A02 | NA | 8.912 | 2.29 | 0.719 | 6.506 | 0.011 | 0.048 |
| Phycisphaeraceae | SM1A02 | NA | 10.453 | 2.816 | 0.705 | 13.357 | 0 | 0.002 |
| A4b | NA | NA | 3.747 | 2.879 | 0.781 | 10.396 | 0.001 | 0.008 |
| Phormidesmiaceae | Phormidesmis_ANT.LACV5.1 | NA | 43.465 | -2.54 | 0.643 | 9.411 | 0.002 | 0.014 |
| Cyanobiaceae | Synechococcus_CC9902 | NA | 2.218 | -3.287 | 1.151 | 7.02 | 0.008 | 0.039 |
| Rubinisphaeraceae | Planctomicrobium | NA | 8.154 | -2.872 | 0.57 | 15.903 | 0 | 0.001 |
| Rubinisphaeraceae | Planctomicrobium | NA | 3.569 | -3.076 | 1.177 | 6.779 | 0.009 | 0.043 |
| Rubinisphaeraceae | NA | NA | 32.582 | -1.449 | 0.308 | 17.168 | 0 | 0 |
| Rubinisphaeraceae | Fuerstia | NA | 17.75 | 3.07 | 0.573 | 21.4 | 0 | 0 |
| Gimesiaceae | NA | NA | 5.592 | -3.822 | 1.026 | 10.255 | 0.001 | 0.009 |
| Pirellulaceae | Blastopirellula | NA | 53.818 | 8.559 | 0.534 | 164.158 | 0 | 0 |
| Pirellulaceae | Blastopirellula | NA | 20.059 | 3.628 | 0.734 | 19.854 | 0 | 0 |
| Pirellulaceae | Blastopirellula | NA | 12.862 | -2.702 | 0.795 | 6.967 | 0.008 | 0.04 |
| Pirellulaceae | Blastopirellula | NA | 3.239 | 4.279 | 1.473 | 7.067 | 0.008 | 0.038 |
| Sulfurimonadaceae | Sulfurimonas | NA | 7.374 | 2.775 | 0.668 | 13.374 | 0 | 0.002 |
| Sulfurovaceae | Sulfurovum | NA | 29.021 | -1.865 | 0.42 | 19.997 | 0 | 0 |
| Sulfurovaceae | Sulfurovum | NA | 19.689 | -2.74 | 0.611 | 20.037 | 0 | 0 |
| Sulfurovaceae | Sulfurovum | NA | 7.568 | -3.331 | 1.275 | 6.901 | 0.009 | 0.041 |
| Rubritaleaceae | Persicirhabdus | NA | 14.955 | -2.204 | 0.553 | 11.999 | 0.001 | 0.004 |
| Rubritaleaceae | Roseibacillus | NA | 5.282 | 3.648 | 0.715 | 19.178 | 0 | 0 |
| Rubritaleaceae | Haloferula | NA | 8.628 | -2.009 | 0.611 | 7.741 | 0.005 | 0.029 |
| DEV007 | NA | NA | 3.709 | -2.602 | 0.952 | 7.062 | 0.008 | 0.038 |
| DEV007 | NA | NA | 5.643 | 3.991 | 0.772 | 19.152 | 0 | 0 |
| DEV007 | NA | NA | 1.345 | -3.882 | 1.217 | 9.424 | 0.002 | 0.014 |
| DEV007 | NA | NA | 5.969 | 3.758 | 0.89 | 14.693 | 0 | 0.001 |
| DEV007 | NA | NA | 3.107 | 4.233 | 1.357 | 8.373 | 0.004 | 0.022 |
| DEV007 | NA | NA | 7.53 | -4.445 | 0.813 | 21.601 | 0 | 0 |
| Chromatiaceae | Candidatus_Thiobios | NA | 12.243 | 2.649 | 0.654 | 13 | 0 | 0.003 |
| Granulosicoccaceae | Granulosicoccus | NA | 7.316 | 2.589 | 0.744 | 8.285 | 0.004 | 0.023 |
| Granulosicoccaceae | Granulosicoccus | NA | 70.11 | 8.672 | 0.479 | 217.091 | 0 | 0 |
| Granulosicoccaceae | Granulosicoccus | coccoides | 88.566 | 2.875 | 0.356 | 38.95 | 0 | 0 |
| Granulosicoccaceae | Granulosicoccus | NA | 38.921 | 4.95 | 0.623 | 32.006 | 0 | 0 |
| Granulosicoccaceae | Granulosicoccus | NA | 98.839 | -1.145 | 0.285 | 12.325 | 0 | 0.003 |
| Granulosicoccaceae | Granulosicoccus | NA | 199.733 | 2.775 | 0.36 | 37.587 | 0 | 0 |
| Granulosicoccaceae | Granulosicoccus | NA | 11.799 | -8.909 | 1.188 | 7.164 | 0.007 | 0.036 |
| Arenicellaceae | Arenicella | NA | 40.558 | 6.123 | 0.529 | 80.905 | 0 | 0 |
| Tenderiaceae | Candidatus_Tenderia | NA | 48.719 | 1.9 | 0.308 | 27.633 | 0 | 0 |
| Nitrosococcaceae | Cm1-21 | NA | 12.848 | 3.067 | 0.593 | 20.113 | 0 | 0 |
| Woeseiaceae | Woeseia | NA | 6.635 | -4.683 | 0.872 | 17.224 | 0 | 0 |
| Woeseiaceae | Woeseia | NA | 10.216 | -2.693 | 0.951 | 7.189 | 0.007 | 0.036 |
| Chromatiaceae | Halochromatium | NA | 9.886 | -2.297 | 0.652 | 10.278 | 0.001 | 0.009 |
| Chromatiaceae | Candidatus_Thiobios | NA | 12.874 | 3.304 | 0.75 | 13.093 | 0 | 0.003 |
| Spongiibacteraceae | NA | NA | 79.124 | 4.305 | 0.421 | 66.924 | 0 | 0 |
| Spongiibacteraceae | Dasania | NA | 5.253 | 4.259 | 1.067 | 13.016 | 0 | 0.003 |
| Unknown_Family | NA | NA | 46.475 | -1 | 0.278 | 11.761 | 0.001 | 0.005 |
| Halieaceae | Pseudohaliea | NA | 61.517 | 2.424 | 0.622 | 6.78 | 0.009 | 0.043 |
| Halieaceae | Halioglobus | NA | 11.167 | -2.47 | 0.806 | 9.191 | 0.002 | 0.015 |
| Halieaceae | OM60(NOR5)_clade | NA | 5.148 | 4.4 | 1.53 | 6.643 | 0.01 | 0.046 |
| Halieaceae | OM60(NOR5)_clade | NA | 13.032 | 4.966 | 0.95 | 7.873 | 0.005 | 0.027 |
| Pseudohongiellaceae | Pseudohongiella | NA | 9.859 | 2.463 | 0.767 | 8.088 | 0.004 | 0.025 |
| Pseudohongiellaceae | Pseudohongiella | NA | 5.867 | 4.888 | 1.097 | 15.393 | 0 | 0.001 |
| Halomonadaceae | Halomonas | NA | 7.736 | -1.838 | 0.721 | 6.965 | 0.008 | 0.04 |
| Methylophilaceae | Methylotenera | NA | 19.433 | -2.776 | 0.639 | 18.896 | 0 | 0 |
| Methylophilaceae | Methylotenera | NA | 48.788 | 6.084 | 0.551 | 72.969 | 0 | 0 |
| Methylophilaceae | Methylotenera | NA | 16.319 | 7.072 | 1.294 | 17.882 | 0 | 0 |
| Methylophilaceae | Methylotenera | NA | 10.263 | 6.059 | 0.78 | 21.601 | 0 | 0 |
| Methylophilaceae | Methylotenera | NA | 74.915 | 1.757 | 0.275 | 31.366 | 0 | 0 |
| Methylophilaceae | Methylotenera | NA | 101.147 | 5.619 | 0.402 | 114.354 | 0 | 0 |
| Alteromonadaceae | Glaciecola | NA | 7.855 | 3.486 | 0.847 | 13.305 | 0 | 0.002 |
| Alteromonadaceae | Salinimonas | NA | 14.407 | 4.924 | 0.714 | 27.346 | 0 | 0 |
| Alteromonadaceae | Glaciecola | NA | 13.459 | 5.044 | 0.89 | 17.452 | 0 | 0 |
| Alteromonadaceae | Glaciecola | NA | 28.536 | 5.272 | 0.759 | 18.424 | 0 | 0 |
| Alteromonadaceae | Glaciecola | NA | 67.191 | 6.54 | 0.546 | 82.122 | 0 | 0 |
| Alteromonadaceae | Glaciecola | punicea | 19.681 | 4.648 | 0.718 | 21.564 | 0 | 0 |
| Alteromonadaceae | Glaciecola | NA | 5.141 | 3.312 | 1.121 | 7.232 | 0.007 | 0.035 |
| Alteromonadaceae | Paraglaciecola | aestuariivivens | 17.429 | 2.727 | 0.63 | 11.502 | 0.001 | 0.005 |
| Methylophagaceae | NA | NA | 13.535 | 5.669 | 1.001 | 20.499 | 0 | 0 |
| Methylophagaceae | Marine_Methylotrophic_Group_3 | NA | 3.056 | 4.159 | 1.271 | 8.757 | 0.003 | 0.018 |
| Colwelliaceae | Thalassotalea | NA | 2.35 | 4.308 | 1.234 | 11 | 0.001 | 0.006 |
| Colwelliaceae | Colwellia | polaris | 49.793 | 5.937 | 0.492 | 86.222 | 0 | 0 |
| Colwelliaceae | Colwellia | NA | 56.557 | 6.761 | 0.725 | 47.785 | 0 | 0 |
| Colwelliaceae | Colwellia | NA | 10.742 | 6.413 | 1.438 | 12.726 | 0 | 0.003 |
| Kangiellaceae | NA | NA | 30.73 | 4.06 | 0.618 | 23.604 | 0 | 0 |
| Marinomonadaceae | Marinomonas | NA | 29.874 | 6.298 | 0.621 | 61.838 | 0 | 0 |
| Nitrincolaceae | NA | NA | 11.053 | 5.848 | 1.035 | 20.299 | 0 | 0 |
| Nitrincolaceae | NA | NA | 8.767 | 4.503 | 0.99 | 11.195 | 0.001 | 0.006 |
| Thiotrichaceae | Cocleimonas | NA | 8.189 | -2.181 | 0.762 | 7.588 | 0.006 | 0.031 |
| Thiotrichaceae | Leucothrix | NA | 87.708 | 3.565 | 0.64 | 20.471 | 0 | 0 |
| Thioglobaceae | SUP05_cluster | NA | 8.529 | 2.308 | 0.619 | 11.108 | 0.001 | 0.006 |
| Shewanellaceae | Shewanella | NA | 7.961 | -1.77 | 0.634 | 7.543 | 0.006 | 0.031 |
| Cellvibrionaceae | Candidatus_Endobugula | NA | 4.083 | 5.047 | 1.255 | 12.462 | 0 | 0.003 |
| Oleiphilaceae | Oleiphilus | NA | 29.446 | 7.314 | 0.808 | 46.885 | 0 | 0 |
| Cellvibrionaceae | Agaribacterium | NA | 14.523 | 4.49 | 0.619 | 35.387 | 0 | 0 |
| Cellvibrionaceae | NA | NA | 7.665 | 4.14 | 1.188 | 7.995 | 0.005 | 0.025 |
| Porticoccaceae | C1-B045 | NA | 3.894 | 2.936 | 0.935 | 6.452 | 0.011 | 0.05 |
| Legionellaceae | NA | NA | 1.611 | -3.981 | 1.45 | 11.381 | 0.001 | 0.005 |
| Unknown_Family | Marinicella | NA | 41.506 | -1.523 | 0.453 | 10.433 | 0.001 | 0.008 |
| Francisellaceae | NA | NA | 21.001 | 8.184 | 1.305 | 8.74 | 0.003 | 0.019 |
| Flavobacteriaceae | Actibacter | NA | 27.912 | -1.208 | 0.43 | 7.45 | 0.006 | 0.032 |
| Flavobacteriaceae | NA | NA | 14.867 | -4.987 | 0.903 | 17.028 | 0 | 0 |
| Flavobacteriaceae | Wenyingzhuangia | NA | 3.6 | 4.241 | 1.358 | 8.274 | 0.004 | 0.023 |
| Flavobacteriaceae | Psychroserpens | damuponensis | 163.122 | -0.832 | 0.222 | 13.295 | 0 | 0.002 |
| Flavobacteriaceae | Winogradskyella | eximia | 5.059 | -3.408 | 1.064 | 13.056 | 0 | 0.003 |
| Flavobacteriaceae | Formosa | NA | 7.625 | 26.169 | 1.376 | 9.074 | 0.003 | 0.016 |
| Flavobacteriaceae | NA | NA | 62.359 | -2.037 | 0.379 | 29.102 | 0 | 0 |
| Flavobacteriaceae | Aquibacter | NA | 4.978 | -4.899 | 1.132 | 15.574 | 0 | 0.001 |
| Flavobacteriaceae | NS2b_marine_group | NA | 22.179 | 4.756 | 0.869 | 18.764 | 0 | 0 |
| Flavobacteriaceae | Aquibacter | NA | 136.221 | -0.718 | 0.244 | 8.657 | 0.003 | 0.019 |
| Flavobacteriaceae | Winogradskyella | NA | 37.176 | -4.605 | 1.046 | 7.263 | 0.007 | 0.035 |
| Flavobacteriaceae | Winogradskyella | echinorum | 169.052 | 1.134 | 0.29 | 12.586 | 0 | 0.003 |
| Flavobacteriaceae | Ulvibacter | NA | 46.538 | 5.25 | 0.646 | 46.06 | 0 | 0 |
| Flavobacteriaceae | Changchengzhania | NA | 81.885 | 2.788 | 0.456 | 24.249 | 0 | 0 |
| Flavobacteriaceae | Ulvibacter | NA | 31.51 | 4.21 | 0.794 | 14.424 | 0 | 0.001 |
| Flavobacteriaceae | Ulvibacter | NA | 23.023 | -6.897 | 1.091 | 21.659 | 0 | 0 |
| Flavobacteriaceae | Ulvibacter | NA | 13.125 | -3.424 | 0.905 | 10.393 | 0.001 | 0.008 |
| Flavobacteriaceae | Jejudonia | NA | 7.608 | -3.467 | 0.879 | 8.39 | 0.004 | 0.022 |
| Flavobacteriaceae | Kordia | NA | 16.918 | 7.12 | 1.127 | 23.686 | 0 | 0 |
| Flavobacteriaceae | Kordia | jejudonensis | 84.057 | 6.287 | 0.506 | 83.773 | 0 | 0 |
| Flavobacteriaceae | Kordia | NA | 8.547 | 4.704 | 1.028 | 15.263 | 0 | 0.001 |
| Flavobacteriaceae | Lutibacter | NA | 8.27 | -2.362 | 0.905 | 6.684 | 0.01 | 0.045 |
| Flavobacteriaceae | Polaribacter | NA | 83.88 | 8.138 | 0.742 | 59.917 | 0 | 0 |
| Flavobacteriaceae | Aurantivirga | NA | 34.717 | 30 | 1.464 | 6.823 | 0.009 | 0.042 |
| Flavobacteriaceae | Aurantivirga | NA | 43.018 | 7.59 | 0.685 | 60.947 | 0 | 0 |
| Flavobacteriaceae | NA | NA | 10.502 | -2.345 | 0.682 | 8.488 | 0.004 | 0.021 |
| Prolixibacteraceae | Draconibacterium | NA | 14.124 | 2.401 | 0.538 | 13.836 | 0 | 0.002 |
| Prolixibacteraceae | Draconibacterium | NA | 5.147 | 3.852 | 0.995 | 11.114 | 0.001 | 0.006 |
| Crocinitomicaceae | Crocinitomix | NA | 17.445 | 7.287 | 0.987 | 32.168 | 0 | 0 |
| Crocinitomicaceae | Crocinitomix | NA | 29.794 | 1.697 | 0.491 | 7.229 | 0.007 | 0.035 |
| Crocinitomicaceae | Crocinitomix | NA | 3.657 | 2.899 | 0.967 | 7.262 | 0.007 | 0.035 |
| Crocinitomicaceae | Crocinitomix | NA | 6.543 | 3.504 | 1.265 | 6.627 | 0.01 | 0.046 |
| Schleiferiaceae | Schleiferia | NA | 2.695 | 4.457 | 1.158 | 13.028 | 0 | 0.003 |
| KD3-93 | NA | NA | 9.621 | 4.956 | 1.117 | 13.099 | 0 | 0.003 |
| 37-13 | NA | NA | 7.434 | 3.614 | 0.746 | 16.292 | 0 | 0.001 |
| Saprospiraceae | NA | NA | 10.913 | 3.175 | 0.736 | 9.105 | 0.003 | 0.016 |
| Saprospiraceae | Phaeodactylibacter | NA | 10.234 | 6.374 | 0.695 | 57.915 | 0 | 0 |
| Saprospiraceae | NA | NA | 7.622 | 5.277 | 0.864 | 27.418 | 0 | 0 |
| Saprospiraceae | NA | NA | 3.708 | 3.874 | 1.207 | 8.944 | 0.003 | 0.017 |
| Saprospiraceae | Lewinella | NA | 10.492 | 4.041 | 0.742 | 18.784 | 0 | 0 |
| Saprospiraceae | Lewinella | NA | 9.958 | 2.117 | 0.704 | 8.038 | 0.005 | 0.025 |
| Saprospiraceae | Aureispira | NA | 22.142 | 3.47 | 0.97 | 9.511 | 0.002 | 0.013 |
| Saprospiraceae | NA | NA | 26.584 | 6.396 | 0.816 | 36.542 | 0 | 0 |
| Saprospiraceae | NA | NA | 6.653 | -2.635 | 0.693 | 14.908 | 0 | 0.001 |
| Saprospiraceae | Lewinella | persica | 26.305 | 3.478 | 0.405 | 49.519 | 0 | 0 |
| Saprospiraceae | NA | NA | 3.911 | 4.566 | 1.105 | 13.673 | 0 | 0.002 |
| Saprospiraceae | NA | NA | 22.334 | 4.472 | 0.657 | 31.39 | 0 | 0 |
| Saprospiraceae | NA | NA | 23.994 | 2.307 | 0.797 | 6.626 | 0.01 | 0.046 |
| Saprospiraceae | NA | NA | 4.948 | 4.978 | 1.625 | 6.644 | 0.01 | 0.046 |
| Saprospiraceae | NA | NA | 9.042 | 5.089 | 0.877 | 23.782 | 0 | 0 |
| Saprospiraceae | NA | NA | 3.998 | 4.524 | 1.171 | 11.272 | 0.001 | 0.006 |
| Saprospiraceae | NA | NA | 5.664 | 4.783 | 0.948 | 19.147 | 0 | 0 |
| Saprospiraceae | Portibacter | NA | 8.745 | 4.355 | 1.073 | 9.747 | 0.002 | 0.012 |
| Saprospiraceae | Portibacter | NA | 36.801 | 2.67 | 0.414 | 29.718 | 0 | 0 |
| Saprospiraceae | Portibacter | NA | 13.002 | 3.033 | 0.675 | 14.467 | 0 | 0.001 |
| Saprospiraceae | NA | NA | 18.712 | 5.968 | 0.566 | 74.104 | 0 | 0 |
| Saprospiraceae | Lewinella | NA | 4.4 | 3.593 | 0.982 | 12.147 | 0 | 0.004 |
| Saprospiraceae | NA | NA | 24.814 | 2.37 | 0.514 | 13.523 | 0 | 0.002 |
| Saprospiraceae | NA | NA | 36.746 | 2.649 | 0.471 | 21.386 | 0 | 0 |
| Saprospiraceae | Lewinella | NA | 2.819 | 4.134 | 1.397 | 7.611 | 0.006 | 0.031 |
| Saprospiraceae | NA | NA | 19.155 | 5.564 | 0.784 | 30.945 | 0 | 0 |
| Saprospiraceae | NA | NA | 7.515 | 4.502 | 1.274 | 8.288 | 0.004 | 0.023 |
| Saprospiraceae | NA | NA | 23.024 | 4.718 | 0.706 | 25.509 | 0 | 0 |
| Saprospiraceae | NA | NA | 21.959 | 7.175 | 0.579 | 103.085 | 0 | 0 |
| Saprospiraceae | Rubidimonas | NA | 14.501 | 5.359 | 1.29 | 11.091 | 0.001 | 0.006 |
| Saprospiraceae | Rubidimonas | NA | 109.575 | 6.939 | 0.688 | 51.708 | 0 | 0 |
| Saprospiraceae | Rubidimonas | NA | 18.514 | 6.16 | 0.67 | 51.32 | 0 | 0 |
| Saprospiraceae | NA | NA | 11.54 | 2.452 | 0.626 | 9.817 | 0.002 | 0.011 |
| Saprospiraceae | NA | NA | 15.491 | 3.611 | 1.102 | 11.235 | 0.001 | 0.006 |
| Saprospiraceae | NA | NA | 20.482 | 2.702 | 1.001 | 13.127 | 0 | 0.002 |
| Saprospiraceae | Portibacter | NA | 2.484 | 3.88 | 1.39 | 6.779 | 0.009 | 0.043 |
| Saprospiraceae | NA | NA | 9.948 | 3.838 | 0.766 | 12.41 | 0 | 0.003 |
| Cryomorphaceae | Vicingus | NA | 11.109 | 3.707 | 0.846 | 13.168 | 0 | 0.002 |
| Cryomorphaceae | Vicingus | NA | 32.457 | 7.346 | 0.847 | 23.021 | 0 | 0 |
| Cryomorphaceae | Vicingus | NA | 4.908 | 4.196 | 1.205 | 8.651 | 0.003 | 0.019 |
| Cryomorphaceae | NA | NA | 4.305 | 3.02 | 1.073 | 6.56 | 0.01 | 0.047 |
| NS9_marine_group | NA | NA | 11.853 | 6.496 | 0.878 | 34.587 | 0 | 0 |
| Crocinitomicaceae | NA | NA | 6.366 | 4.661 | 1.126 | 13.043 | 0 | 0.003 |
| Flavobacteriaceae | NA | NA | 3.327 | -4.745 | 0.957 | 19.698 | 0 | 0 |
| Flavobacteriaceae | Maribacter | NA | 50.778 | -1.311 | 0.34 | 12.07 | 0.001 | 0.004 |
| Flavobacteriaceae | Maribacter | NA | 66.931 | 0.906 | 0.252 | 11.399 | 0.001 | 0.005 |
| Flavobacteriaceae | Maribacter | NA | 26.899 | -1.917 | 0.563 | 8.112 | 0.004 | 0.024 |
| Flavobacteriaceae | Robiginitalea | NA | 16.938 | -1.239 | 0.397 | 8.381 | 0.004 | 0.022 |
| Cryomorphaceae | NA | NA | 7.531 | 3.737 | 0.782 | 11.425 | 0.001 | 0.005 |
| Cryomorphaceae | NA | NA | 12.525 | 3.11 | 0.581 | 21.487 | 0 | 0 |
| Crocinitomicaceae | Fluviicola | NA | 11.315 | 4.608 | 0.936 | 13.531 | 0 | 0.002 |
| Crocinitomicaceae | Fluviicola | NA | 58.146 | 3.489 | 0.523 | 26.35 | 0 | 0 |
| Crocinitomicaceae | Fluviicola | NA | 18.887 | 4.978 | 0.779 | 24.776 | 0 | 0 |
| Cryomorphaceae | NA | NA | 12.221 | 4.501 | 0.454 | 76.481 | 0 | 0 |
| Cryomorphaceae | NA | NA | 3.592 | 4.352 | 0.946 | 17.693 | 0 | 0 |
| NS11-12_marine_group | NA | NA | 4.99 | 3.29 | 0.904 | 9.233 | 0.002 | 0.015 |
| Caldilineaceae | NA | NA | 40.907 | 3.326 | 0.499 | 32.77 | 0 | 0 |
| Caldilineaceae | NA | NA | 4.929 | 2.726 | 0.856 | 8.028 | 0.005 | 0.025 |
| Ilumatobacteraceae | Ilumatobacter | nonamiensis | 9.81 | -2.21 | 0.713 | 6.894 | 0.009 | 0.041 |
| Microtrichaceae | Sva0996_marine_group | NA | 8.358 | -2.223 | 0.523 | 13.066 | 0 | 0.003 |
| Microtrichaceae | NA | NA | 23.994 | 2.486 | 0.515 | 13.648 | 0 | 0.002 |
| Microtrichaceae | Sva0996_marine_group | NA | 21.773 | 4.342 | 0.573 | 22.638 | 0 | 0 |
| Microtrichaceae | Sva0996_marine_group | NA | 4.327 | -4.728 | 1.128 | 11.759 | 0.001 | 0.005 |
| Bdellovibrionaceae | OM27_clade | NA | 1.491 | 3.325 | 1.189 | 7.513 | 0.006 | 0.032 |
| Desulfobulbaceae | Desulfobulbus | NA | 8.679 | -1.793 | 0.548 | 8.815 | 0.003 | 0.018 |
| Magnetospiraceae | Magnetospira | NA | 1.403 | 2.953 | 1 | 6.877 | 0.009 | 0.041 |
| Terasakiellaceae | NA | NA | 50.386 | 5.978 | 0.673 | 47.836 | 0 | 0 |
| Rhodobacteraceae | NA | NA | 4.761 | 3.165 | 1.095 | 8.287 | 0.004 | 0.023 |
| Rhodobacteraceae | Thiobacimonas | NA | 17.034 | -2.585 | 0.419 | 33.568 | 0 | 0 |
| Hyphomicrobiaceae | Filomicrobium | NA | 11.358 | -2.824 | 0.516 | 21.894 | 0 | 0 |
| Hyphomonadaceae | Hyphomonas | NA | 6.509 | -3.295 | 0.666 | 11.57 | 0.001 | 0.005 |
| Hyphomonadaceae | NA | NA | 15.484 | -1.484 | 0.468 | 9.011 | 0.003 | 0.016 |
| Hyphomonadaceae | Hellea | balneolensis | 39.771 | 3.474 | 0.336 | 77.082 | 0 | 0 |
| Hyphomonadaceae | Litorimonas | NA | 21.269 | 3.428 | 0.508 | 29.41 | 0 | 0 |
| Hyphomonadaceae | Robiginitomaculum | NA | 14.774 | 4.295 | 0.709 | 28.385 | 0 | 0 |
| Rhodobacteraceae | Limibaculum | NA | 15.209 | 2.261 | 0.702 | 6.552 | 0.01 | 0.047 |
| Rhodobacteraceae | NA | NA | 18.134 | 4.292 | 0.528 | 43.637 | 0 | 0 |
| Rhodobacteraceae | NA | NA | 237.052 | 5.295 | 0.344 | 125.144 | 0 | 0 |
| Rhodobacteraceae | NA | NA | 33.738 | 5.933 | 0.586 | 57.45 | 0 | 0 |
| Rhodobacteraceae | NA | NA | 49.822 | 5.718 | 0.548 | 64.236 | 0 | 0 |
| Rhodobacteraceae | Planktomarina | NA | 7.091 | -5.703 | 1.451 | 7.493 | 0.006 | 0.032 |
| Rhodobacteraceae | Yoonia-Loktanella | NA | 8.328 | 4.319 | 1.001 | 8.13 | 0.004 | 0.024 |
| Rhodobacteraceae | Jannaschia | NA | 13.245 | -4.349 | 1.021 | 13.706 | 0 | 0.002 |
| Rhodobacteraceae | NA | NA | 16.788 | 2.021 | 0.679 | 7.572 | 0.006 | 0.031 |
| Rhodobacteraceae | NA | NA | 7.131 | -4.683 | 0.97 | 21.32 | 0 | 0 |
| Rhodobacteraceae | Octadecabacter | NA | 56.508 | 3.938 | 0.593 | 22.544 | 0 | 0 |
| Rhodobacteraceae | Octadecabacter | NA | 156.039 | -1.299 | 0.192 | 41.669 | 0 | 0 |
| Rhodobacteraceae | NA | NA | 14.799 | -3.378 | 1.237 | 6.566 | 0.01 | 0.047 |
| Rhodobacteraceae | NA | NA | 11.581 | 5.488 | 1.351 | 12.774 | 0 | 0.003 |
| Rhodobacteraceae | Tropicimonas | NA | 2.936 | -3.376 | 1.386 | 7.126 | 0.008 | 0.037 |
| Rhodobacteraceae | Planktotalea | NA | 215.331 | 0.867 | 0.205 | 16.153 | 0 | 0.001 |
| Rhodobacteraceae | Aliiroseovarius | NA | 38.776 | 6.41 | 0.873 | 16.093 | 0 | 0.001 |
| Rhodobacteraceae | Celeribacter | NA | 39.186 | -1.797 | 0.348 | 25.378 | 0 | 0 |
| Rhodobacteraceae | NA | NA | 39.828 | 3.621 | 0.655 | 19.521 | 0 | 0 |
| Rhodobacteraceae | NA | NA | 396.234 | 1.906 | 0.197 | 75.046 | 0 | 0 |
| Rhodobacteraceae | NA | NA | 34.638 | 8.145 | 0.638 | 96.542 | 0 | 0 |
| Rhodobacteraceae | NA | NA | 7.774 | 2.771 | 0.831 | 9.561 | 0.002 | 0.013 |
| Rhodobacteraceae | NA | NA | 93.88 | 2.532 | 0.314 | 47.621 | 0 | 0 |
| Rhodobacteraceae | NA | NA | 5.446 | -3.574 | 1.05 | 11.531 | 0.001 | 0.005 |
| Rhodobacteraceae | NA | NA | 9.58 | 5.864 | 1.187 | 13.246 | 0 | 0.002 |
| Rhodobacteraceae | Sulfitobacter | litoralis | 56.8 | -1.091 | 0.29 | 12.069 | 0.001 | 0.004 |
| Rhodobacteraceae | Sulfitobacter | NA | 19.103 | 5.52 | 1.101 | 13.601 | 0 | 0.002 |
| Rhodobacteraceae | Sulfitobacter | NA | 5.922 | -4.762 | 1.089 | 8.312 | 0.004 | 0.022 |
| Rhodobacteraceae | NA | NA | 7.249 | 5.253 | 1.198 | 12.154 | 0 | 0.004 |
| Rhodobacteraceae | NA | NA | 14.135 | 4.263 | 1.283 | 7.646 | 0.006 | 0.03 |
| Rhodobacteraceae | Sulfitobacter | brevis | 26.953 | -2.629 | 0.799 | 7.544 | 0.006 | 0.031 |
| Rhodobacteraceae | Sulfitobacter | NA | 8.387 | 2.842 | 1.544 | 7.172 | 0.007 | 0.036 |
| Rhodobacteraceae | Sulfitobacter | NA | 44.071 | 2.408 | 0.611 | 10.671 | 0.001 | 0.007 |
| Rhodobacteraceae | Roseobacter | litoralis | 44.601 | -1.685 | 0.479 | 10.195 | 0.001 | 0.009 |
| Rhodobacteraceae | Sedimentitalea | NA | 24.21 | -3.976 | 0.624 | 31.691 | 0 | 0 |
| Rhodobacteraceae | NA | NA | 5 | -5.515 | 1.212 | 18.105 | 0 | 0 |
| Rhodobacteraceae | NA | NA | 10.167 | 5.693 | 1.648 | 7.367 | 0.007 | 0.034 |
| Rhodobacteraceae | NA | NA | 12.148 | -3.867 | 1.096 | 6.834 | 0.009 | 0.042 |
| Rhodobacteraceae | Phaeobacter | NA | 17.193 | -4.9 | 0.682 | 39.286 | 0 | 0 |
| Rhodobacteraceae | Roseobacter | NA | 2.589 | -3.541 | 1.223 | 6.952 | 0.008 | 0.04 |
| Rhodobacteraceae | Roseobacter | NA | 5.032 | 2.999 | 1 | 7.483 | 0.006 | 0.032 |
| Rhodobacteraceae | NA | NA | 39.161 | 5 | 0.653 | 33.659 | 0 | 0 |
| Rhodobacteraceae | Roseovarius | aestuarii | 6.174 | -5.562 | 1.075 | 20.78 | 0 | 0 |
| Rhodobacteraceae | NA | NA | 18.498 | 5.922 | 0.801 | 28.482 | 0 | 0 |
| Rhodobacteraceae | NA | NA | 29.657 | 4.196 | 0.499 | 47.792 | 0 | 0 |
| Rhodobacteraceae | Sedimentitalea | NA | 72.267 | 3.72 | 0.335 | 76.604 | 0 | 0 |
| Rhodobacteraceae | NA | NA | 4.505 | -4.858 | 0.812 | 30.442 | 0 | 0 |
| Rhodobacteraceae | NA | NA | 32.365 | 4.94 | 0.469 | 70.674 | 0 | 0 |
| Rhodobacteraceae | Yoonia-Loktanella | NA | 34.968 | 3.062 | 0.741 | 13.684 | 0 | 0.002 |
| Rhodobacteraceae | NA | NA | 67.096 | 3.101 | 0.382 | 43.72 | 0 | 0 |
| Rhodobacteraceae | NA | NA | 9.982 | 4.452 | 1.113 | 11.187 | 0.001 | 0.006 |
| Rhodobacteraceae | NA | NA | 43.239 | 1.969 | 0.367 | 18.645 | 0 | 0 |
| Rhodobacteraceae | Leisingera | NA | 4.582 | -3.532 | 1.212 | 9.016 | 0.003 | 0.016 |
| Rhodobacteraceae | Yoonia-Loktanella | NA | 33.729 | 2.94 | 0.441 | 28.475 | 0 | 0 |
| Rhodobacteraceae | Yoonia-Loktanella | NA | 41.605 | 1.922 | 0.33 | 27.066 | 0 | 0 |
| Rhodobacteraceae | NA | NA | 3.688 | -2.19 | 0.853 | 8.252 | 0.004 | 0.023 |
| Rhodobacteraceae | NA | NA | 142.747 | 3.365 | 0.268 | 113.259 | 0 | 0 |
| Rhizobiaceae | Pseudahrensia | NA | 27.282 | 3.23 | 0.493 | 29.577 | 0 | 0 |
| Rhizobiaceae | Pseudahrensia | NA | 25.716 | -1.803 | 0.513 | 8.774 | 0.003 | 0.018 |
| Rhizobiaceae | Pseudahrensia | NA | 14.5 | -6.513 | 0.755 | 56.359 | 0 | 0 |
| Rhodobacteraceae | NA | NA | 3.52 | -4.799 | 0.9 | 30.766 | 0 | 0 |
| Rhodobacteraceae | NA | NA | 43.352 | -2.124 | 0.324 | 39.636 | 0 | 0 |
| Rhizobiaceae | Pseudahrensia | NA | 1.834 | -3.883 | 1.361 | 8.095 | 0.004 | 0.025 |
| Rhizobiaceae | Pseudahrensia | NA | 11.6 | -4.553 | 0.605 | 36.848 | 0 | 0 |
| Rhizobiaceae | Hoeflea | NA | 4.176 | -5.011 | 1.201 | 14.866 | 0 | 0.001 |
| Sphingomonadaceae | NA | NA | 24.79 | -2.579 | 0.665 | 12.928 | 0 | 0.003 |
| Sphingomonadaceae | Sphingorhabdus | flavimaris | 92.56 | 0.782 | 0.241 | 9.529 | 0.002 | 0.013 |
| Sphingomonadaceae | Erythrobacter | NA | 21.103 | 2.313 | 0.523 | 11.58 | 0.001 | 0.005 |
| Sphingomonadaceae | Altererythrobacter | NA | 4.57 | -3.245 | 1.07 | 8.586 | 0.003 | 0.02 |
| Sphingomonadaceae | Erythrobacter | NA | 41.883 | 1.533 | 0.452 | 8.512 | 0.004 | 0.021 |
| Saprospiraceae | NA | NA | 32.989 | 6.413 | 0.704 | 48.194 | 0 | 0 |
| Saprospiraceae | NA | NA | 21.658 | 6.067 | 0.996 | 23.925 | 0 | 0 |
| Saprospiraceae | NA | NA | 10.295 | 2.788 | 0.848 | 7.714 | 0.005 | 0.029 |
| Spirosomaceae | Taeseokella | NA | 73.479 | 3.419 | 0.353 | 66.673 | 0 | 0 |
| Cyclobacteriaceae | Ekhidna | NA | 15.733 | 2.265 | 0.559 | 11.683 | 0.001 | 0.005 |
| Cyclobacteriaceae | Fabibacter | NA | 10.48 | 3.844 | 0.782 | 10.791 | 0.001 | 0.007 |
| Flammeovirgaceae | NA | NA | 2.513 | 3.807 | 0.892 | 14.721 | 0 | 0.001 |
| Bernardetiaceae | Garritya | NA | 4.128 | 2.58 | 0.981 | 7.351 | 0.007 | 0.034 |
| Flavobacteriaceae | NS3a_marine_group | NA | 210.383 | 4.102 | 0.635 | 22.917 | 0 | 0 |
| Rhodothermaceae | NA | NA | 26.217 | 3.323 | 0.343 | 63.158 | 0 | 0 |
| Opitutaceae | Diplosphaera | NA | 2.134 | 3.574 | 0.966 | 13.002 | 0 | 0.003 |
| Desulfocapsaceae | NA | NA | 5.584 | 5.601 | 1.19 | 15.667 | 0 | 0.001 |
| Desulfocapsaceae | NA | NA | 15.71 | -1.583 | 0.53 | 6.922 | 0.009 | 0.04 |
| Desulfocapsaceae | NA | NA | 4.262 | -4.053 | 1.226 | 8.726 | 0.003 | 0.019 |
| Desulfocapsaceae | NA | NA | 56.902 | -0.982 | 0.292 | 9.961 | 0.002 | 0.011 |
| Desulfocapsaceae | NA | NA | 58.318 | -0.855 | 0.267 | 9.419 | 0.002 | 0.014 |
| Desulfocapsaceae | NA | NA | 15.688 | -1.507 | 0.506 | 7.291 | 0.007 | 0.035 |
| Fokiniaceae | MD3-55 | NA | 17.984 | 5.724 | 0.457 | 123.977 | 0 | 0 |
| Rhizobiales_Incertae_Sedis | Anderseniella | NA | 1.913 | -4.19 | 1.435 | 7.334 | 0.007 | 0.034 |
| Rhizobiaceae | NA | NA | 2.925 | -4.812 | 1.758 | 9.343 | 0.002 | 0.014 |
| Rhizobiaceae | NA | NA | 7.129 | -4.65 | 0.771 | 26.066 | 0 | 0 |
| Rhodobacteraceae | Tateyamaria | NA | 95.34 | 3.682 | 0.354 | 70.102 | 0 | 0 |
| Rhodobacteraceae | Tateyamaria | NA | 8.586 | -6.07 | 1.056 | 31.575 | 0 | 0 |
| Rhodobacteraceae | NA | NA | 9.963 | 1.942 | 0.624 | 7.465 | 0.006 | 0.032 |
| Rhodobacteraceae | NA | NA | 39.212 | 5.346 | 0.542 | 54.736 | 0 | 0 |
| Rhizobiaceae | Ahrensia | NA | 3.369 | -4.055 | 1.25 | 7.463 | 0.006 | 0.032 |
| Sphingomonadaceae | Parasphingopyxis | NA | 8.183 | -4.998 | 0.824 | 23.271 | 0 | 0 |
| Micavibrionaceae | NA | NA | 2.428 | -4.625 | 1.199 | 13.902 | 0 | 0.002 |
| Micavibrionaceae | NA | NA | 9.96 | -3.209 | 0.783 | 17.994 | 0 | 0 |
| Rickettsiaceae | Candidatus_Megaira | NA | 69.951 | 6.412 | 0.559 | 76.103 | 0 | 0 |
| Rickettsiaceae | Candidatus_Megaira | NA | 6.7 | 4.91 | 1.041 | 16.495 | 0 | 0.001 |
| Bdellovibrionaceae | Bdellovibrio | NA | 2.308 | 2.82 | 1.009 | 8.031 | 0.005 | 0.025 |

Appendix Supplemental Table 4

| **Pathway** | **log2-fold Change** |
| --- | --- |
| nitrifier denitrification | 3.23766261 |
| superpathway of polyamine biosynthesis III | 2.50832917 |
| CMP-pseudaminate biosynthesis | 2.38533794 |
| nylon-6 oligomer degradation | 1.78133925 |
| thiazole biosynthesis II (Bacillus) | 1.10325976 |
| coenzyme M biosynthesis I | 1.04456344 |
| formaldehyde oxidation I | 0.9590049 |
| formaldehyde assimilation II (RuMP Cycle) | 0.9562312 |
| methyl ketone biosynthesis | 0.9450924 |
| superpathway of thiamin diphosphate biosynthesis II | 0.92452883 |
| ectoine biosynthesis | 0.70480998 |
| ADP-L-glycero-&beta;-D-manno-heptose biosynthesis | 0.65819635 |
| L-arginine degradation II (AST pathway) | 0.63231738 |
| glucose and glucose-1-phosphate degradation | 0.60402905 |
| norspermidine biosynthesis | 0.57577995 |
| superpathway of polyamine biosynthesis I | 0.56942389 |
| superpathway of histidine, purine, and pyrimidine biosynthesis | -0.5242397 |
| thiazole biosynthesis I (E. coli) | -0.5275914 |
| fucose degradation | -0.5326074 |
| catechol degradation to &beta;-ketoadipate | -0.5359885 |
| 2-aminophenol degradation | -0.543549 |
| acetylene degradation | -0.5581539 |
| superpathway of Clostridium acetobutylicum acidogenic fermentation | -0.5613171 |
| mannan degradation | -0.5810783 |
| pyruvate fermentation to butanoate | -0.5920957 |
| pyrimidine deoxyribonucleotides de novo biosynthesis II | -0.6078529 |
| nitrate reduction VI (assimilatory) | -0.6120458 |
| superpathway of pyridoxal 5'-phosphate biosynthesis and salvage | -0.6644642 |
| superpathway of purine nucleotides de novo biosynthesis II | -0.6876697 |
| aerobactin biosynthesis | -0.6890469 |
| superpathway of sulfur oxidation (Acidianus ambivalens) | -0.7026774 |
| superpathway of salicylate degradation | -0.8300772 |
| reductive acetyl coenzyme A pathway | -0.8371843 |
| teichoic acid (poly-glycerol) biosynthesis | -0.8410264 |
| catechol degradation III (ortho-cleavage pathway) | -0.8454332 |
| aromatic compounds degradation via &beta;-ketoadipate | -0.8454332 |
| meta cleavage pathway of aromatic compounds | -0.8789704 |
| succinate fermentation to butanoate | -0.8998533 |
| adenosylcobalamin biosynthesis II (late cobalt incorporation) | -0.9365552 |
| androstenedione degradation | -0.9391163 |
| superpathway of 2,3-butanediol biosynthesis | -0.9661671 |
| superpathway of (Kdo)2-lipid A biosynthesis | -0.9733811 |
| methanogenesis from acetate | -0.9792468 |
| superpathway of demethylmenaquinol-6 biosynthesis II | -1.0619267 |
| isopropanol biosynthesis | -1.0728644 |
| superpathway of hexitol degradation (bacteria) | -1.0874184 |
| L-glutamate degradation V (via hydroxyglutarate) | -1.1359106 |
| D-galactarate degradation I | -1.1440026 |
| superpathway of D-glucarate and D-galactarate degradation | -1.1440026 |
| superpathway of (R,R)-butanediol biosynthesis | -1.1442403 |
| pyruvate fermentation to acetone | -1.1605646 |
| formaldehyde assimilation I (serine pathway) | -1.1923269 |
| factor 420 biosynthesis | -1.2158772 |
| isoprene biosynthesis II (engineered) | -1.2826466 |
| glycerol degradation to butanol | -1.3412872 |
| superpathway of N-acetylneuraminate degradation | -1.3670503 |
| superpathway of L-aspartate and L-asparagine biosynthesis | -1.4278134 |
| 1,5-anhydrofructose degradation | -1.4325517 |
| superpathway of N-acetylglucosamine, N-acetylmannosamine and N-acetylneuraminate degradation | -1.4891301 |
| coenzyme B biosynthesis | -1.6276845 |
| allantoin degradation to glyoxylate III | -1.6391955 |
| D-glucarate degradation I | -1.6718886 |
| glutaryl-CoA degradation | -1.7367827 |
| L-lysine fermentation to acetate and butanoate | -1.7522809 |
| creatinine degradation II | -1.7894301 |
| glucose degradation (oxidative) | -2.1076198 |
| mono-trans, poly-cis decaprenyl phosphate biosynthesis | -2.2966947 |
| methylaspartate cycle | -2.3760429 |
| L-lysine biosynthesis II | -2.4960889 |
| NAD salvage pathway II | -2.5786537 |
| cob(II)yrinate a,c-diamide biosynthesis I (early cobalt insertion) | -2.7391815 |
| chondroitin sulfate degradation I (bacterial) | -2.8135174 |
| L-glutamate degradation VIII (to propanoate) | -2.8832619 |
| peptidoglycan biosynthesis IV (Enterococcus faecium) | -2.9437995 |
| 3-phenylpropanoate and 3-(3-hydroxyphenyl)propanoate degradation to 2-oxopent-4-enoate | -3.0650628 |
| cinnamate and 3-hydroxycinnamate degradation to 2-oxopent-4-enoate | -3.0650628 |
| allantoin degradation IV (anaerobic) | -3.3360909 |
| superpathway of L-arginine, putrescine, and 4-aminobutanoate degradation | -3.5753343 |
| superpathway of L-arginine and L-ornithine degradation | -3.5753343 |
| nicotinate degradation I | -3.8065081 |
| starch degradation III | -5.1566723 |
| peptidoglycan biosynthesis V (&beta;-lactam resistance) | -8.1998858 |

Appendix Supplemental Table 5

|  |  |  | df | Sum Of Squares | R^2^ | F-Statistic | Pr(>F) |
| --- | --- | --- | --- | --- | --- | --- | --- |
| Based on taxonomy | Root vs Mimic | Sample Type | 1 | 2854.7 | 0.14 | 16.116 | **0.001** |
|  |  | Residual | 99 | 17536.1 | 0.86 |  |  |
|  |  | Total | 100 | 20390.8 | 1 |  |  |
|  | Mimic vs. Sediment | Sample Type | 1 | 4684.5 | 0.37082 | 35.362 | **0.001** |
|  |  | Residual | 60 | 7948.3 | 0.62918 |  |  |
|  |  | Total | 61 | 12632.8 | 1 |  |  |
|  | Root vs. Sediment | Sample Type | 1 | 6239.1 | 0.27549 | 41.447 | **0.001** |
|  |  | Residual | 109 | 16408 | 0.72451 |  |  |
|  |  | Total | 110 | 22647.1 | 1 |  |  |
| Based on predicted function | Root vs Mimic | Sample Type | 1 | 3038 | 0.0511 | 5.3312 | **0.001** |
|  |  | Residual | 99 | 56417 | 0.9489 |  |  |
|  |  | Total | 100 | 59455 | 1 |  |  |
|  | Mimic vs. Sediment | Sample Type | 1 | 8870 | 0.19271 | 14.323 | **0.001** |
|  |  | Residual | 60 | 37157 | 0.80729 |  |  |
|  |  | Total | 61 | 46028 | 1 |  |  |
|  | Root vs. Sediment | Sample Type | 1 | 12146 | 0.21179 | 29.289 | **0.001** |
|  |  | Residual | 109 | 45201 | 0.78821 |  |  |
|  |  | Total | 110 | 57347 | 1 |  |  |

Appendix Supplemental Table 6

| **Family** | **Higher on roots** | **Higher on mimics** | **Higher on mimics** | **Higher on sediment** | **Higher on roots** | **Higher on sediment** |
| --- | --- | --- | --- | --- | --- | --- |
| Acanthopleuribacteraceae | 1 | 0 | 0 | 1 | 0 | 1 |
| Anaerolineaceae | 3 | 0 | 1 | 5 | 1 | 5 |
| Arenicellaceae | 3 | 1 | 2 | 1 | 3 | 1 |
| Bacteroidetes_BD2-2 | 14 | 1 | 0 | 16 | 3 | 13 |
| Calditrichaceae | 5 | 3 | 0 | 9 | 0 | 9 |
| Cellulomonadaceae | 1 | 0 | 1 | 0 | 1 | 0 |
| Cellvibrionaceae | 3 | 0 | 2 | 0 | 3 | 0 |
| Chitinophagaceae | 1 | 0 | 1 | 0 | 1 | 0 |
| Christensenellaceae | 2 | 0 | 0 | 1 | 1 | 1 |
| Chromatiaceae | 3 | 0 | 0 | 4 | 0 | 4 |
| Crocinitomicaceae | 4 | 0 | 5 | 0 | 5 | 0 |
| Cyclobacteriaceae | 5 | 0 | 1 | 3 | 2 | 3 |
| Desulfobacteraceae | 3 | 0 | 0 | 2 | 1 | 2 |
| Desulfobulbaceae | 2 | 0 | 0 | 4 | 1 | 3 |
| Desulfocapsaceae | 24 | 0 | 6 | 15 | 14 | 10 |
| Desulfosarcinaceae | 14 | 3 | 0 | 23 | 0 | 23 |
| Desulfovibrionaceae | 5 | 0 | 0 | 1 | 5 | 0 |
| DEV007 | 3 | 0 | 3 | 1 | 4 | 0 |
| Devosiaceae | 1 | 0 | 1 | 0 | 1 | 0 |
| Ectothiorhodospiraceae | 1 | 1 | 0 | 2 | 0 | 2 |
| Flavobacteriaceae | 38 | 3 | 32 | 12 | 37 | 8 |
| Fusibacteraceae | 2 | 0 | 0 | 1 | 2 | 0 |
| Gemmatimonadaceae | 1 | 0 | 0 | 1 | 0 | 1 |
| Gimesiaceae | 1 | 0 | 1 | 0 | 1 | 0 |
| Granulosicoccaceae | 2 | 1 | 5 | 0 | 5 | 0 |
| Halieaceae | 4 | 0 | 1 | 4 | 1 | 4 |
| Halomonadaceae | 0 | 2 | 2 | 0 | 2 | 0 |
| Hungateiclostridiaceae | 3 | 0 | 0 | 3 | 1 | 2 |
| Hyphomonadaceae | 1 | 0 | 4 | 0 | 4 | 0 |
| Kiritimatiellaceae | 1 | 0 | 0 | 1 | 0 | 1 |
| Kordiimonadaceae | 1 | 0 | 1 | 0 | 1 | 0 |
| Lachnospiraceae | 6 | 0 | 2 | 1 | 6 | 0 |
| Latescibacteraceae | 0 | 2 | 0 | 3 | 0 | 3 |
| Lentimicrobiaceae | 3 | 0 | 0 | 3 | 0 | 3 |
| Magnetospiraceae | 1 | 0 | 1 | 0 | 1 | 0 |
| Marinifilaceae | 2 | 0 | 0 | 1 | 2 | 0 |
| Marinilabiliaceae | 5 | 1 | 0 | 6 | 3 | 2 |
| Marinomonadaceae | 1 | 0 | 1 | 0 | 1 | 0 |
| Melioribacteraceae | 6 | 0 | 1 | 4 | 3 | 3 |
| Methyloligellaceae | 1 | 0 | 0 | 1 | 0 | 1 |
| Methylophagaceae | 2 | 0 | 2 | 0 | 2 | 0 |
| Methylophilaceae | 4 | 0 | 4 | 0 | 4 | 0 |
| Microtrichaceae | 2 | 0 | 2 | 0 | 2 | 0 |
| MSBL8 | 1 | 0 | 0 | 2 | 0 | 2 |
| Nitrincolaceae | 2 | 0 | 1 | 1 | 2 | 0 |
| Parvibaculaceae | 1 | 0 | 1 | 0 | 1 | 0 |
| PHOS-HE36 | 2 | 0 | 0 | 3 | 0 | 3 |
| Pirellulaceae | 16 | 1 | 13 | 7 | 13 | 6 |
| Prolixibacteraceae | 7 | 0 | 2 | 4 | 6 | 2 |
| Psychromonadaceae | 1 | 0 | 1 | 0 | 1 | 0 |
| Puniceicoccaceae | 2 | 0 | 1 | 0 | 2 | 0 |
| Rhizobiaceae | 6 | 0 | 6 | 0 | 7 | 0 |
| Rhodobacteraceae | 36 | 3 | 41 | 0 | 45 | 0 |
| Rubinisphaeraceae | 5 | 0 | 5 | 0 | 5 | 0 |
| Rubritaleaceae | 3 | 0 | 3 | 0 | 3 | 0 |
| Sandaracinaceae | 1 | 0 | 0 | 1 | 0 | 1 |
| Saprospiraceae | 19 | 3 | 21 | 4 | 23 | 3 |
| SB-5 | 6 | 0 | 0 | 6 | 2 | 4 |
| Schleiferiaceae | 1 | 0 | 1 | 0 | 1 | 0 |
| Sedimenticolaceae | 2 | 2 | 1 | 5 | 1 | 5 |
| SG8-4 | 1 | 0 | 0 | 1 | 0 | 1 |
| Shewanellaceae | 0 | 1 | 1 | 0 | 1 | 0 |
| Sphingomonadaceae | 2 | 0 | 2 | 0 | 3 | 0 |
| Spirochaetaceae | 9 | 1 | 0 | 10 | 7 | 5 |
| Spirosomaceae | 1 | 0 | 2 | 0 | 2 | 0 |
| Spongiibacteraceae | 2 | 0 | 2 | 1 | 2 | 1 |
| Sulfurovaceae | 1 | 0 | 1 | 0 | 1 | 0 |
| Syntrophotaleaceae | 1 | 0 | 0 | 1 | 1 | 0 |
| Thermoanaerobaculaceae | 7 | 0 | 0 | 11 | 0 | 11 |
| Thioalkalispiraceae | 1 | 2 | 0 | 3 | 0 | 3 |
| Thiomicrospiraceae | 4 | 1 | 1 | 6 | 1 | 6 |
| Thiotrichaceae | 8 | 1 | 9 | 3 | 9 | 3 |
| Trueperaceae | 1 | 0 | 1 | 0 | 1 | 0 |
| Unknown_Family | 5 | 1 | 1 | 6 | 1 | 5 |
| Vibrionaceae | 1 | 0 | 1 | 0 | 1 | 0 |
| Woeseiaceae | 2 | 0 | 0 | 2 | 0 | 2 |

Appendix Supplemental Table 7

See file at github.org/mkardish/Mimics/AppendixASuppTab7.xlsx

Appendix Supplemental Table 8

| **Pathway** | **Root vs. Mimic** | **Mimic vs. Sediment** | **Root vs. Sediment** |
| --- | --- | --- | --- |
| &beta;-alanine biosynthesis II | NA | 5.98028867 | 6.17916013 |
| 1,4-dihydroxy-6-naphthoate biosynthesis I | 0.52453186 | -0.8382554 | NA |
| 1,4-dihydroxy-6-naphthoate biosynthesis II | NA | -0.9208415 | NA |
| 1,5-anhydrofructose degradation | NA | 0.5822825 | 0.91678643 |
| 2-amino-3-carboxymuconate semialdehyde degradation to 2-oxopentenoate | -0.5324918 | 2.10192706 | 1.56943525 |
| 2-aminophenol degradation | -1.0096637 | 2.07887674 | 1.06921309 |
| 2-methylcitrate cycle I | -0.9007436 | 0.59329576 | NA |
| 2-methylcitrate cycle II | -0.7081516 | NA | NA |
| 2-nitrobenzoate degradation I | -0.5048366 | 1.98333606 | 1.47849943 |
| 3-phenylpropanoate and 3-(3-hydroxyphenyl)propanoate degradation to 2-oxopent-4-enoate | -1.3020418 | 4.07512185 | 2.77308003 |
| 3-phenylpropanoate degradation | -1.3795198 | 7.21839716 | 5.83887733 |
| 4-coumarate degradation (anaerobic) | NA | 0.90868496 | 1.0680199 |
| 4-hydroxyphenylacetate degradation | -0.6235443 | 1.70481364 | 1.0812693 |
| 4-methylcatechol degradation (ortho cleavage) | -2.1204126 | 2.63553665 | 0.51512402 |
| adenosylcobalamin biosynthesis I (early cobalt insertion) | -0.5278635 | 2.38312108 | 1.85525759 |
| adenosylcobalamin biosynthesis II (late cobalt incorporation) | NA | 2.45438795 | 2.19260435 |
| ADP-L-glycero-&beta;-D-manno-heptose biosynthesis | NA | -1.0379013 | -1.0447171 |
| aerobactin biosynthesis | NA | 2.67822173 | 2.97740096 |
| allantoin degradation IV (anaerobic) | -3.8790422 | 11.6679446 | 7.78890236 |
| allantoin degradation to glyoxylate III | -1.0948528 | 1.20012867 | NA |
| androstenedione degradation | NA | -1.0590578 | -0.7936352 |
| aromatic biogenic amine degradation (bacteria) | NA | 0.97177605 | 0.64651199 |
| aromatic compounds degradation via &beta;-ketoadipate | -1.7048087 | 2.56052172 | 0.85571297 |
| benzoyl-CoA degradation I (aerobic) | -5.3493261 | 8.43200767 | 3.08268157 |
| benzoyl-CoA degradation II (anaerobic) | 0.81885539 | -3.1649666 | -2.3461112 |
| Bifidobacterium shunt | NA | 1.36268466 | 0.9583059 |
| biotin biosynthesis II | -2.2801293 | 5.1067305 | 2.82660117 |
| catechol degradation III (ortho-cleavage pathway) | -1.7048087 | 2.56052172 | 0.85571297 |
| catechol degradation to &beta;-ketoadipate | -0.8640468 | 2.3152049 | 1.4511581 |
| catechol degradation to 2-oxopent-4-enoate II | NA | 1.29739008 | 1.0656568 |
| chitin derivatives degradation | 0.83765223 | 1.23367148 | 2.07132371 |
| chlorophyllide a biosynthesis I (aerobic, light-dependent) | NA | 1.66916567 | 1.65806178 |
| chlorophyllide a biosynthesis II (anaerobic) | NA | 1.63793953 | 1.6191585 |
| chlorophyllide a biosynthesis III (aerobic, light independent) | NA | 1.63793953 | 1.6191585 |
| chlorosalicylate degradation | -1.5806965 | 5.29068882 | 3.70999227 |
| chondroitin sulfate degradation I (bacterial) | -3.5706214 | 2.1288075 | -1.4418139 |
| cinnamate and 3-hydroxycinnamate degradation to 2-oxopent-4-enoate | -1.3020418 | 4.07512185 | 2.77308003 |
| CMP-legionaminate biosynthesis I | NA | -1.7170811 | -1.7560234 |
| CMP-pseudaminate biosynthesis | 0.96889146 | 2.61729121 | 3.58618266 |
| cob(II)yrinate a,c-diamide biosynthesis I (early cobalt insertion) | NA | 1.95843292 | 1.70820751 |
| cob(II)yrinate a,c-diamide biosynthesis II (late cobalt incorporation) | NA | 1.06300152 | 1.22997949 |
| coenzyme B biosynthesis | NA | 6.33172398 | 5.84288679 |
| coenzyme M biosynthesis I | NA | 0.55913072 | NA |
| creatinine degradation I | NA | 1.47125265 | 1.73472301 |
| creatinine degradation II | NA | 2.02886451 | 2.48406197 |
| D-fructuronate degradation | NA | 0.84183251 | 0.54886057 |
| D-galactarate degradation I | NA | 0.5502454 | NA |
| D-galacturonate degradation I | NA | 0.68804147 | 0.60597012 |
| D-glucarate degradation I | -0.7791992 | NA | NA |
| dTDP-N-acetylthomosamine biosynthesis | NA | 0.82091307 | NA |
| ectoine biosynthesis | NA | 0.6063511 | 0.7610108 |
| enterobacterial common antigen biosynthesis | -3.9398588 | 8.08370873 | 4.1438499 |
| enterobactin biosynthesis | -1.6985363 | 1.81330954 | NA |
| ergothioneine biosynthesis I (bacteria) | -4.5808794 | 3.71990392 | -0.8609754 |
| ethylmalonyl-CoA pathway | NA | 1.63626912 | 1.93902201 |
| factor 420 biosynthesis | -3.6181246 | 7.7968298 | 4.17870521 |
| formaldehyde assimilation II (RuMP Cycle) | NA | 1.36498138 | 1.62215714 |
| formaldehyde oxidation I | NA | 1.34472111 | 1.60981698 |
| gallate degradation I | NA | 2.96335443 | 2.60732399 |
| gallate degradation II | NA | 2.9931866 | 2.60959275 |
| GDP-D-glycero-&alpha;-D-manno-heptose biosynthesis | NA | -2.336863 | -2.3533359 |
| glucose and glucose-1-phosphate degradation | -0.6154652 | 0.87514308 | NA |
| glucose degradation (oxidative) | -3.5634038 | 1.31205774 | -2.251346 |
| glutaryl-CoA degradation | -0.8744803 | -1.4823571 | -2.3568374 |
| glycerol degradation to butanol | -1.2138411 | 2.11371278 | 0.89987173 |
| glycine betaine degradation I | NA | 1.52630641 | 1.77967665 |
| glycogen degradation I (bacterial) | NA | -0.5022613 | NA |
| glycogen degradation II (eukaryotic) | NA | 1.92811354 | 1.48703839 |
| glyoxylate cycle | NA | 0.59545961 | NA |
| heterolactic fermentation | NA | 1.35594303 | 0.9030291 |
| hexitol fermentation to lactate, formate, ethanol and acetate | -2.3564217 | 2.93058383 | 0.57416213 |
| incomplete reductive TCA cycle | NA | -0.6320642 | NA |
| isoprene biosynthesis II (engineered) | NA | -1.596346 | -1.2874942 |
| isopropanol biosynthesis | NA | -0.7395089 | -0.6601573 |
| ketogluconate metabolism | -1.1100443 | 2.96513108 | 1.85508681 |
| L-arabinose degradation IV | NA | 9.80122018 | 9.49234434 |
| L-arginine degradation II (AST pathway) | -2.0305001 | 2.94130845 | 0.91080839 |
| L-glutamate degradation V (via hydroxyglutarate) | NA | -1.5896498 | -1.8924772 |
| L-histidine degradation II | NA | 1.82293122 | 1.87999775 |
| L-isoleucine biosynthesis IV | NA | -0.6183816 | NA |
| L-lysine biosynthesis II | -2.7233821 | 5.1544283 | 2.43104615 |
| L-lysine fermentation to acetate and butanoate | NA | 1.47736849 | 1.57053027 |
| L-methionine biosynthesis I | NA | 0.71356601 | 0.64118985 |
| L-methionine salvage cycle III | -5.7266642 | 7.85970156 | 2.1330374 |
| L-rhamnose degradation I | -0.5624761 | 0.61263037 | NA |
| L-tryptophan degradation IX | NA | 1.00068122 | 0.80181412 |
| L-tryptophan degradation to 2-amino-3-carboxymuconate semialdehyde | NA | 1.00325492 | 0.74569097 |
| L-tryptophan degradation XII (Geobacillus) | -0.7230692 | 1.62974146 | 0.90667225 |
| L-tyrosine degradation I | NA | 0.85425311 | 0.81798341 |
| L-valine degradation I | NA | 6.3388524 | 6.49180761 |
| lactose and galactose degradation I | -5.062184 | 8.28938471 | 3.22720075 |
| mannan degradation | NA | 0.97400331 | 0.60637761 |
| meta cleavage pathway of aromatic compounds | -1.0709935 | 2.48553208 | 1.41453861 |
| methanogenesis from acetate | 0.66802212 | -1.5177292 | -0.8497071 |
| methanol oxidation to carbon dioxide | -0.6092689 | 1.67128139 | 1.06201246 |
| methylaspartate cycle | NA | NA | 0.72955726 |
| methylgallate degradation | NA | 2.96817215 | 2.59940031 |
| methylphosphonate degradation I | NA | 1.21248246 | 1.44671709 |
| mevalonate pathway I | NA | -0.6973192 | NA |
| mevalonate pathway II (archaea) | 1.19113556 | -2.5647924 | -1.3736568 |
| mono-trans, poly-cis decaprenyl phosphate biosynthesis | -2.1669721 | 6.66242923 | 4.49545717 |
| mycothiol biosynthesis | -0.5798133 | 0.57033211 | NA |
| myo-, chiro- and scillo-inositol degradation | NA | 2.56222211 | 2.29443289 |
| myo-inositol degradation I | NA | 2.51159939 | 2.38726902 |
| NAD biosynthesis II (from tryptophan) | NA | 0.7850773 | 0.59630288 |
| NAD salvage pathway II | -3.216504 | 3.1998056 | NA |
| nicotinate degradation I | -5.9100989 | 8.01022674 | 2.1001278 |
| nitrate reduction VI (assimilatory) | NA | 1.33504703 | 0.94737502 |
| norspermidine biosynthesis | NA | 1.58842222 | 1.33114879 |
| nylon-6 oligomer degradation | NA | 0.99787962 | 0.80549182 |
| octane oxidation | NA | 1.03817344 | 1.15439139 |
| palmitate biosynthesis II (bacteria and plants) | -0.872762 | -0.8440889 | -1.716851 |
| peptidoglycan biosynthesis II (staphylococci) | -5.788931 | 12.6352847 | 6.84635369 |
| peptidoglycan biosynthesis IV (Enterococcus faecium) | -2.3037652 | 3.02148026 | 0.71771506 |
| peptidoglycan biosynthesis V (&beta;-lactam resistance) | -3.1421431 | 5.05225645 | 1.91011332 |
| phenylacetate degradation I (aerobic) | -1.5928447 | 2.04828109 | NA |
| phospholipases | -0.7007483 | 1.74447062 | 1.04372228 |
| polymyxin resistance | -2.9570821 | 1.86688576 | -1.0901963 |
| ppGpp biosynthesis | NA | 1.3237357 | 0.94719959 |
| protocatechuate degradation I (meta-cleavage pathway) | NA | 3.31323514 | 3.04051865 |
| protocatechuate degradation II (ortho-cleavage pathway) | NA | 1.43269458 | 1.45955473 |
| purine nucleotides degradation II (aerobic) | NA | 0.84745319 | 0.95184059 |
| purine ribonucleosides degradation | NA | 1.02596312 | 1.33210895 |
| pyrimidine deoxyribonucleotides biosynthesis from CTP | NA | -3.2387427 | -3.0061164 |
| pyrimidine deoxyribonucleotides de novo biosynthesis IV | NA | -3.2464337 | -3.0353371 |
| pyruvate fermentation to acetone | -0.7989368 | 0.58953572 | NA |
| pyruvate fermentation to butanoate | NA | -0.8480954 | -0.6834175 |
| reductive acetyl coenzyme A pathway | 0.56299075 | -1.0712442 | -0.5082534 |
| S-adenosyl-L-methionine cycle I | -0.5730782 | 1.49674887 | 0.92367066 |
| S-methyl-5-thio-&alpha;-D-ribose 1-phosphate degradation | -6.0971955 | 7.92182186 | 1.82462635 |
| spirilloxanthin and 2,2'-diketo-spirilloxanthin biosynthesis | NA | 2.2330273 | 2.28217566 |
| starch degradation III | NA | 7.32377146 | 7.21603178 |
| sucrose degradation II (sucrose synthase) | NA | -2.0756477 | -1.9117441 |
| sucrose degradation III (sucrose invertase) | -1.0811735 | 2.13489002 | 1.05371654 |
| superpathway of (Kdo)2-lipid A biosynthesis | NA | -0.7054507 | -1.1461552 |
| superpathway of &beta;-D-glucuronide and D-glucuronate degradation | NA | 0.85148319 | 0.6319071 |
| superpathway of aerobic toluene degradation | -0.7444703 | 1.79585096 | 1.05138063 |
| superpathway of bacteriochlorophyll a biosynthesis | NA | 1.67435543 | 1.69358424 |
| superpathway of C1 compounds oxidation to CO2 | 0.79775953 | 4.93409574 | 5.73185526 |
| superpathway of Clostridium acetobutylicum acidogenic fermentation | NA | -0.7857639 | -0.6260313 |
| superpathway of D-glucarate and D-galactarate degradation | NA | 0.5502454 | NA |
| superpathway of demethylmenaquinol-6 biosynthesis II | NA | 0.81685025 | 1.18219199 |
| superpathway of fucose and rhamnose degradation | -0.8033356 | 3.780394 | 2.97705835 |
| superpathway of geranylgeranyldiphosphate biosynthesis I (via mevalonate) | NA | -0.7049968 | NA |
| superpathway of glycerol degradation to 1,3-propanediol | NA | 2.71236968 | 2.61185182 |
| superpathway of glycol metabolism and degradation | -1.048886 | 4.77755709 | 3.72867113 |
| superpathway of hexitol degradation (bacteria) | -1.4325053 | 1.10451066 | NA |
| superpathway of hexuronide and hexuronate degradation | NA | 1.1797743 | 0.68847957 |
| superpathway of L-arginine and L-ornithine degradation | -4.2622312 | 7.80495629 | 3.54272513 |
| superpathway of L-arginine, putrescine, and 4-aminobutanoate degradation | -4.2622312 | 7.80495629 | 3.54272513 |
| superpathway of L-threonine metabolism | -5.2246486 | 8.59332633 | 3.36867768 |
| superpathway of menaquinol-8 biosynthesis II | 0.67518966 | -0.7613806 | NA |
| superpathway of methylglyoxal degradation | -1.1952535 | 3.88225935 | 2.6870059 |
| superpathway of phenylethylamine degradation | -1.3717238 | 4.68127965 | 3.30955587 |
| superpathway of polyamine biosynthesis III | 1.12820444 | NA | 1.40574398 |
| superpathway of purine deoxyribonucleosides degradation | NA | 0.99995122 | 1.24937609 |
| superpathway of pyridoxal 5'-phosphate biosynthesis and salvage | -0.6116537 | 2.45674032 | 1.84508665 |
| superpathway of pyrimidine deoxyribonucleosides degradation | NA | 0.70003531 | 1.00034021 |
| superpathway of S-adenosyl-L-methionine biosynthesis | NA | 0.51696413 | NA |
| superpathway of salicylate degradation | -1.5799859 | 2.48032888 | 0.90034295 |
| superpathway of sulfolactate degradation | NA | 1.89159187 | 2.27801115 |
| superpathway of sulfur oxidation (Acidianus ambivalens) | 0.76011508 | -1.8921344 | -1.1320193 |
| superpathway of thiamin diphosphate biosynthesis II | NA | -1.1289035 | -0.8492926 |
| superpathway of UDP-glucose-derived O-antigen building blocks biosynthesis | NA | 1.16390318 | 0.98925039 |
| superpathway of vanillin and vanillate degradation | NA | 3.40551039 | 2.99074115 |
| TCA cycle VII (acetate-producers) | NA | 0.78551358 | NA |
| teichoic acid (poly-glycerol) biosynthesis | NA | 4.22846527 | 3.957639 |
| thiazole biosynthesis II (Bacillus) | NA | -1.4247689 | -1.0254904 |
| toluene degradation III (aerobic) (via p-cresol) | -1.5757307 | 2.32241598 | 0.74668526 |
| toluene degradation IV (aerobic) (via catechol) | -1.3085971 | 2.94089177 | 1.63229471 |
| tRNA processing | NA | -0.5698891 | NA |
| UDP-2,3-diacetamido-2,3-dideoxy-&alpha;-D-mannuronate biosynthesis | NA | -0.5764018 | NA |
| urea cycle | NA | 0.52908568 | 0.76921194 |
| vanillin and vanillate degradation I | NA | 3.40551039 | 2.99074115 |
| vanillin and vanillate degradation II | NA | 3.39293769 | 2.98479856 |
| vitamin B6 degradation | -4.8624169 | 6.32522899 | 1.46281206 |
| vitamin E biosynthesis (tocopherols) | -2.7550338 | 7.24500287 | 4.48996911 |
